# Hippocampal-cortical coupling dynamics drive system consolidation of remote memory

**DOI:** 10.64898/2026.06.22.733680

**Authors:** Tao Sheng, Shaoli Wang, Jiaxi Zhang, Danqing Xing, Yi Wu, Qiao Wang, Wei Lu

**Author notes:** These authors contribute equally to this work (Tao Sheng, Shaoli Wang, Jiaxi Zhang, Danqing Xing, Yi Wu). Correspondence (Q.W.), (W.L.).

## Abstract

System consolidation transforms temporary hippocampal representation of memory into long-term storage in cortex. The underlying neural substrate, however, remains enigmatic. Here, we tracked the spatiotemporal evolution of hippocampus (HPC)-cortex local field potentials and single-neuron spikes in behaving animals during fear memory formation. During learning, HPC fast gamma exhibited a progressive phase shift relative to PFC theta oscillations, with gamma power aligning to progressively later phases of the PFC theta cycle. Strikingly, a related phase-shifted coupling pattern re-emerged during subsequent consolidation in association with hippocampal sharp-wave ripples and transient PFC spindle events during NREM sleep. Across this process, interregional interactions evolved from HPC-driven cortical gamma coherence at recent stages to PFC-mediated cortical low-frequency coherence at remote stages. Using closed-loop optogenetic perturbations, we demonstrated a stepwise causal chain of coupling events underlying remote memory formation. Our study revealed HPC-PFC coupling phase shift as a feasible substrate mediating recent-to-remote transformation of memory.

## INTRODUCTION

System consolidation is a crucial aspect of memory processing,^1–8^ during which the brain region responsible for storing the information shifts from hippocampus to cortex.^9–12^ This process engages vast number of cortical neurons for memory storage; thus, it is considered to be a foundation for ensuring the stability and capacity of remote memory storage.^13,14^ Previous work dissecting the neural substrate of system consolidation has been focusing on the synaptic and circuit mechanisms during the early phase of memory consolidation.^15–18^ However, the precise spatiotemporal profiles of neural activities during the later processes, particularly related to the formation of remote memory, are still largely unknown.^19–25^

Specifically, previous studies have identified featured neural oscillations related to mnemonic processes,^26–30^ including theta phase precession during memory encoding and sharp wave ripples (SWRs) during consolidation and retrieval.^31^ These oscillations reflect information processing in local circuit or cross-area communications,^32–36^ and are usually coupled with neural activities critical for neurocomputation. Focusing on system consolidation, one previous study reported gradually increased slow oscillation coherence between prefrontal and motor cortices for motor tasks.^22^ Nevertheless, characterization regarding neural oscillation features underlying system consolidation of remote episodic memory has been scarce. More importantly, causal evidence linking cross-area oscillatory coherence with system consolidation has been missing in the field.

In the present study, we filled in this gap and systematically examined the inter-regional oscillatory crosstalk during system consolidation of episodic memory over weeks. We achieved this with custom-designed multi-channel electrodes that enables simultaneous detection of both local field potentials (LFPs) and neuronal spiking at multiple brain regions in behaving animals.^37^ We further developed closed-loop optogenetic techniques for targeted perturbation of inter-regional coherence to test their causal involvement in memory processes. With these techniques, we captured a previously unappreciated phase shift phenomena in hippocampal-prefrontal oscillatory dialogue. We discovered that the phase shift happens during memory encoding and re-occurs throughout system consolidation. Further intervention experiment demonstrated that the phase shift represents gradual re-organizations in the HPC-cortical circuits, which are necessary for system consolidations.

## RESULTS

### Progressive HPC-PFC coupling phase shift over fear learning

To obtain a panoramic view of cross-area communication related to remote memory formation, we trained rats with a trace fear-conditioning paradigm to associate a contextual cue with freezing responses induced by electrical stimuli at the paws^38^, and performed longitudinal in vivo simultaneous recording in the dorsal CA1 region of hippocampus and multiple cortical regions including prelimbic subdivision of prefrontal cortex (PFC), anterior cingulate cortex (ACC) and posterior parietal cortex (PPC; **Figure 1A**). All these regions are engaged in mnemonic process including trace and contextual fear conditioning.^7,39–47^ Placement of recording electrodes in these regions was confirmed after the experiment (**Figure S1A**).

**Figure 1.**
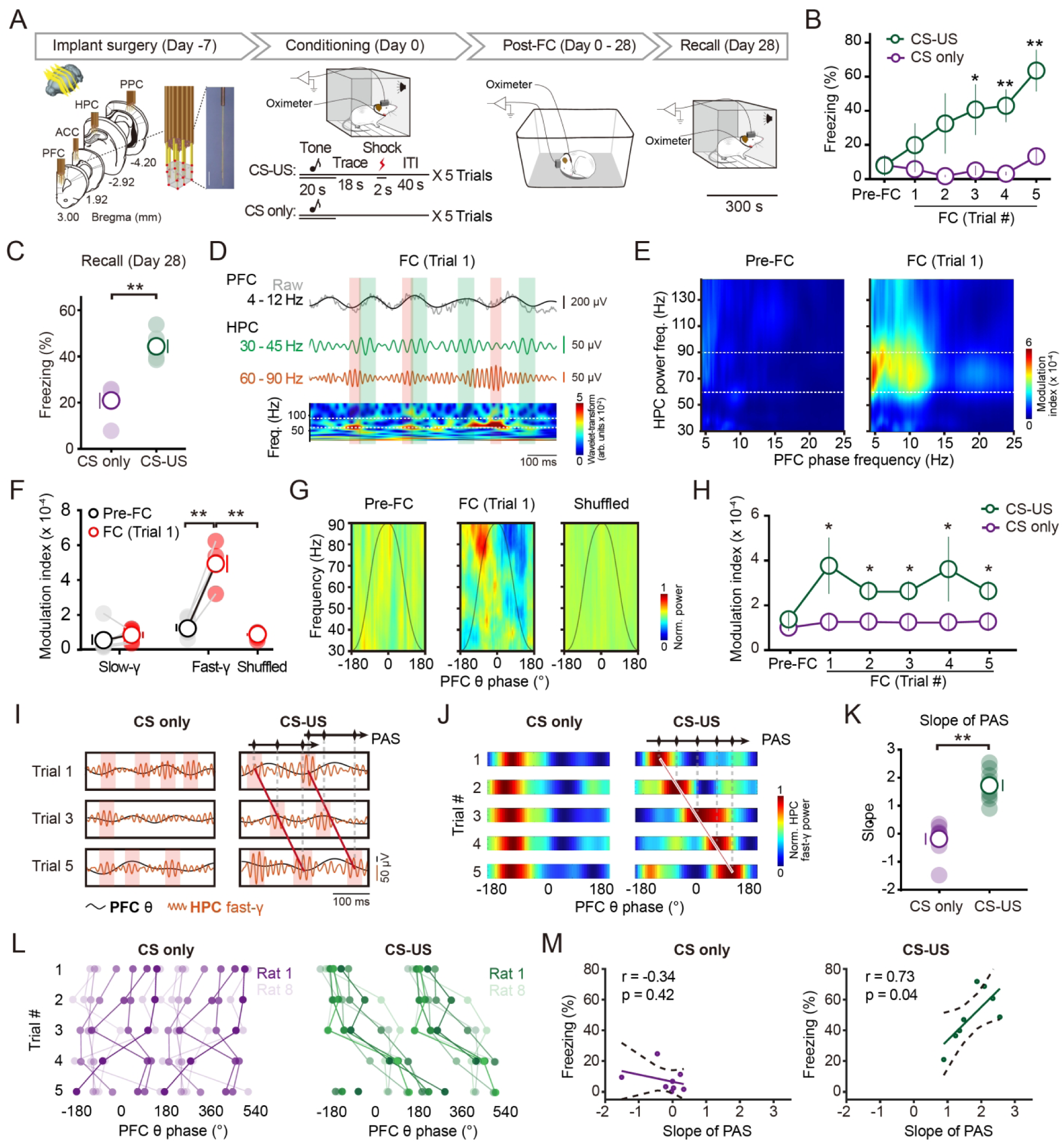
Progressive HPC-PFC coupling phase shift over fear conditioning. (A) Electrodes targeting and experimental schedule. Electrodes respectively targeted HPC and multiple cortical regions including PFC, ACC and PPC. Upper bar shows experimental timeline. Insets, close-up views of the electrodes show flanking recording tips at both lateral and vertical sites with 250 μm spacing intervals. ITI, inter-trial interval. (B) Successful formation of associative memory in CS-US group. Freezing percentage as a function of fear condition (FC) trials was displayed, which was significantly higher in CS-US group than that of CS only group (trial #5, two-way ANOVA, F = 2.88, CS only group, n = 6 rats; CS-US, n = 6 rats; \**P* < 0.05, \*\**P* < 0.01). (C) The formation of remote fear memory was validated by successful recall on day 28 post-FC in CS-US group (green) but not in CS only group (CS only: 21.0 ± 3.4, CS-US: 44.4 ± 2.8, unpaired t-test, CS only group, n = 6 rats; CS-US, n = 6 rats; \*\**P* < 0.01). Centre values denote mean ± s.e.m. (D) Sample of time-aligned LFP trace obtained after trial 1 and wavelet transform in PFC and HPC. Top, raw (gray) and θ band-passed (4-12Hz, black) trace of PFC LFP. Middle, slow (30-45 Hz; green) and fast-γ (60-90 Hz; orange) oscillations. Note that fast-γ oscillations locked to ascending phase of PFC θ oscillation, in contrast to that slow-γ oscillations appeared in different PFC θ phases. Bottom, wavelet transform (color plot) of LFP in HPC. Boxes indicate representative slow (green)/fast-γ (orange) events in each frequency band. arb., arbitrary. (E) Representative phase-amplitude comodulogram demonstrating prominent modulation of HPC fast-γ power (regions between the dotted line; y axis) by PFC θ oscillation phase (x axis) at trial 1. Pre-FC, pre-fear conditioning. Warmer colors indicate stronger modulation. (F) Statistical analysis of modulation index reveals selective modulation of HPC fast-γ (60-90 Hz) by PFC θ at trial 1 (red). n = 6 rats, Paired t*-*test, \*\**P* < 0.01. (G) Example comodulograms of PFC θ-HPC γ PAC during pre-FC (left; 30 s before FC), trial 1 (middle), and shuffled predictor of trial 1 data (right). Norm., normalized. Black lines depict PFC θ oscillation phases. (H) The evolvement of cross-frequency PAC strength in HPC-PFC circuit. PFC θ and HPC γ exhibited prominent coupling during conditioning trials in CS-US group. In contrast, no such coupling was observed in CS only animals (Bonferroni corrected, Wilcoxon signed-rank test, CS only group, n = 6 rats; CS-US, n = 6 rats; \**P* < 0.05). (I) An example showing the shift of HPC fast-γ preferred PFC θ phase across conditioning trials. Note the progressively “later” PFC θ phases of HPC fast-γ oscillations across conditioning trials. Shaded boxes indicate the preferred PFC θ phase. Inset, the rhombuses and arrows indicate the preferred phase and direction of PAS, respectively. (J) Representative comodulograms of HPC fast-γ-PFC θ coupling showing trial-to-trial shift of HPC fast-γ preferred PFC θ phase across conditioning trials. (K) Quantification of PAS with slope of phase shift. CS-US group elicited higher slope than that of CS only group (CS only: –0.23 ± 0.27; CS-US: 1.73 ± 0.26, unpaired t-test, CS only group, n = 8 rats; CS-US, n = 8 rats; \*\**P* < 0.01). (L) The cross-frequency PAC preferred phase displayed directional shift in CS-US groups. The starting point of phase shift in CS-US groups gathered in the ascending period of PFC θ phase, and following points preceded in the same direction. Each dot represents preferred phase for a single animal at one trial (CS only: n = 8 rats; CS-US: n = 8 rats). (M) Learning performance positively correlated with PAS in conditioned animals (CS-US group), as indicated by Pearson correlation coefficient of mean percentage of freezing at the end of conditioning and the slope of PAS (CS only: n = 8 rats, r = –0.34, *P* = 0.42; CS-US: n = 8 rats, r = 0.73, *P* = 0.04). Each cycle represents data from an individual animal. Colored lines show results of linear regression fit.

As hippocampus interacts with neocortical regions at the very early stage of memory formation,^16,23,40,43^ we started recordings before the conditioning stage to map out the temporal evolution of “memory-related” change in the strength of HPC-cortical functional connections, and tracked both local neural oscillations and their temporal coupling during day 0-28 after fear conditioning (thereafter termed post-FC). Recordings in the conditioning chamber were restricted to periods of fear conditioning and memory recall, whereas data acquisition before and after conditioning was conducted in the home cage.

Oscillations and oscillatory coupling are strongly dependent on behavioral states.^27,35,36,48,49^ To assess behavioral states during electrophysiological recordings, we simultaneously monitored EEG, EMG and autonomic signals including respiration and heart rates (**Figures S1B and S1C**). Based on EEG and EMG signals, combined with video monitoring, we classified the recorded data into wakefulness and sleep. As freezing is presumably associated with autonomic changes, we further monitored autonomic changes such as heart rate and respiration. This allowed us to distinguish freezing and immobility during conditioning. Through careful observation and differentiation of these behavioral states, we can clearly link the electrophysiological changes recorded thereafter to their corresponding behavioral states.

Rats subjecting to both conditioned (CS) and unconditioned stimuli (US) displayed gradually increased freezing behavior across the conditioning trials. As a control, rats trained with CS only failed to display similar responses to the contextual stimuli (**Figure 1B**). The associative memory was maintained without exposure to the conditioning context until the recall test at day 28 post-FC (**Figure 1C**), indicating the formation of remote memory.

In order to monitor oscillatory dynamics of HPC and cortical regions with precise spatiotemporal resolution, we developed a multi-channel recording system to allow simultaneous detection of LFPs and neuronal spikes in multiple brain regions from behaving rats^37^ (for details see Methods). Using this system, we monitored LFPs simultaneously in HPC and the cortical regions and measured the oscillations during the 40-second inter-trial intervals (ITIs) following each individual foot shock. These oscillations were also analyzed for the CS only group in the corresponding time frames. Pre-FC data were collected during the 3 minutes preceding the FC in each conditioning session. In recall sessions, 300-second time segments were analyzed.

We then decomposed the signal with wavelets to extract separate frequency components from the ongoing LFPs.^50,51^ This analysis revealed PFC low-frequency oscillations at theta (θ; 4-12 Hz) band and two frequency bands of HPC gamma (γ) oscillations (30-45 Hz and 60-90 Hz, thereafter termed “slow-γ” and “fast-γ”, respectively) in the 40-second ITIs after each training trial, which have been shown to reflect distinct neural and behavioral processes.^52–60^ Notably, we found that the fast-γ, but not slow-γ oscillations, locked to the ascending phase of PFC low-frequency oscillations in the first ITI after trial #1 (**Figure 1D**). Moreover, we observed an increase in the power of both HPC fast-γ and PFC θ oscillations during training, suggesting the both power increase might be correlated with fear conditioning (**Figure S1D**). Compared with post-FC resting periods, hippocampal fast-γ and prefrontal θ power were significantly increased during freezing epochs in the conditioning stage (**Figures S1E and S1F**), indicating that these training-related power enhancements are specifically associated with learning behavior rather than immobility. Furthermore, by monitoring the animals’ respiration frequency, we excluded the influence of respiration rhythm alterations on PFC theta activity (**Figures S2**).

As the amplitude of fast oscillations is often modulated by the phase of slow oscillations,^51,61–71^ a phenomenon called cross-frequency phase-amplitude coupling (PAC), we further investigated whether HPC γ rhythms were modulated by PFC θ during fear conditioning. We employed the modulation index (MI), a measure of circular unimodality, to assess the strength of PAC,^56^ and found that HPC fast-γ oscillations were strongly coupled to that of PFC θ frequencies (**Figures 1E and 1F**). In contrast, no similar PAC enhancement was observed for slow-γ or other within– or inter-regional combinations, with the exception of HPC-ACC θ-fast-γ coupling (**Figures S3 and S4**). In addition, we demonstrated that PFC θ oscillations was not volume conducted from HPC.^50,72–74^

The increase in cross-frequency coupling may be due to an increase in coincident coupling caused by independent changes within θ and γ frequency ranges. To examine this possibility, we compared our results to a shuffled data obtained by shifting the γ power relative to the θ phase by x seconds (where x is an integer value between 1 and 30). We found that the shuffled data failed to display strong patterns of cross-frequency coupling (**Figure 1G**). In addition, the PFC-HPC circuit exhibited strong cross-frequency PAC only in conditioned CS-US animals, but not in CS only animals (**Figure 1H**) or between HPC θ and PFC γ (**Figure S1G**). These results suggest that cross-frequency coupling is not caused by coincident occurrence of independent oscillations.

Strikingly, HPC fast-γ power progressively coupled with “later” phases of the PFC θ, exhibiting a gradual phase shift pattern (**Figures 1I and 1J**). Statistical analysis of the “phase slope” using circular-linear regression confirmed this observation (**Figure 1K**),^75^ suggesting the HPC-PFC cross-frequency coupling underwent linear phase shift over conditioning trials of CS-US group. This observation was further confirmed when Pearson’s correlation coefficient (R^2^) was calculated^73^ to assess the goodness-of-fit in phase-trial regression (**Figure S5A**; for details see Methods). The HPC fast-γ preferred PFC θ phase returned to the pre-training level after conditioning, indicating that the phase is effectively reset to the early-trial configuration (see **Figures S5B and S5C**). Notably, this phase shift was absent in HPC-ACC and HPC-PPC circuits (**Figures S3E-S3L**), indicative of selective communication between HPC and PFC at conditioning stage.

The phase shift was different from the conventional θ phase precession of neuronal spiking observed in hippocampal place cells, in which neuronal spikes occur progressively in earlier phases of the LFP theta oscillation.^76–81^ We termed this recessive phase shift of PAC as phase-amplitude shift (“PAS” in short here and thereafter). We found consistent PAS in all the conditioned animals (**Figure 1L**). Across eight rats, the HPC γ power initially peaked at the ascending phase of PFC θ in the early stage, then progressively occurred to the later phases. As a control, no clear phase shift was observed in CS only group, shock only group (**Figure S5D**) and any other within– or inter-regional combinations (**Figures S3 and S4**). Notably, although θ-γ coupling was observed in HPC-ACC circuit, it did not exhibit PAS during conditioning.

We next explored the brain state relevance of the PAS in HPC-PFC circuit. Combined LFP and behavioral recording analysis showed that the phase shift can be detected equally during freezing and non-freezing periods (**Figures S5E and S5F**). Interestingly, we found that the animals with prominent PAS displayed better learning performance than those with less significant PAS (**Figure 1M; Figure S5G**). Here, the learning performance was defined as mean percentage of freezing at the end of conditioning.^82^ Therefore, learning performance in the conditioned animals was positively correlated with the occurrence of PAS. Collectively, these results identify PAS as a novel neural signature in HPC-PFC circuit that may represent an association to fear memory acquisition.

### Phase shift correlates with PFC modulation of HPC spiking

Brain oscillations are important in coordinating the timing of neuronal firing.^47,51,54,83^ In the analogous phase precession phenomenon, place cell spikes occur at progressively earlier θ phases as an animal moving through a cell’s place field, and spikes’ θ phases provide spatial information.^76,77,79–81^ We asked if the learning-associated PAS is correlated with PFC modulation of HPC single unit activity during fear conditioning. To explore this, we analyzed cross-regional phase-locking of hippocampal CA1 neuronal spiking to PFC θ. We found that the counts of HPC units significantly differed in subpopulations of phase-locked and non-phase-locked CA1 neurons (**Figure 2A**), with 37.2% of all HPC units showed significant phase-locking to the PFC θ oscillations in the first ITI (**Figures 2B and 2C**). Moreover, HPC units that were phase-locked to PFC θ showed higher firing rates during FC compared to pre-FC, while CA1 neurons not phase-locked to PFC θ showed no increase (**Figure 2D**).

**Figure 2.**
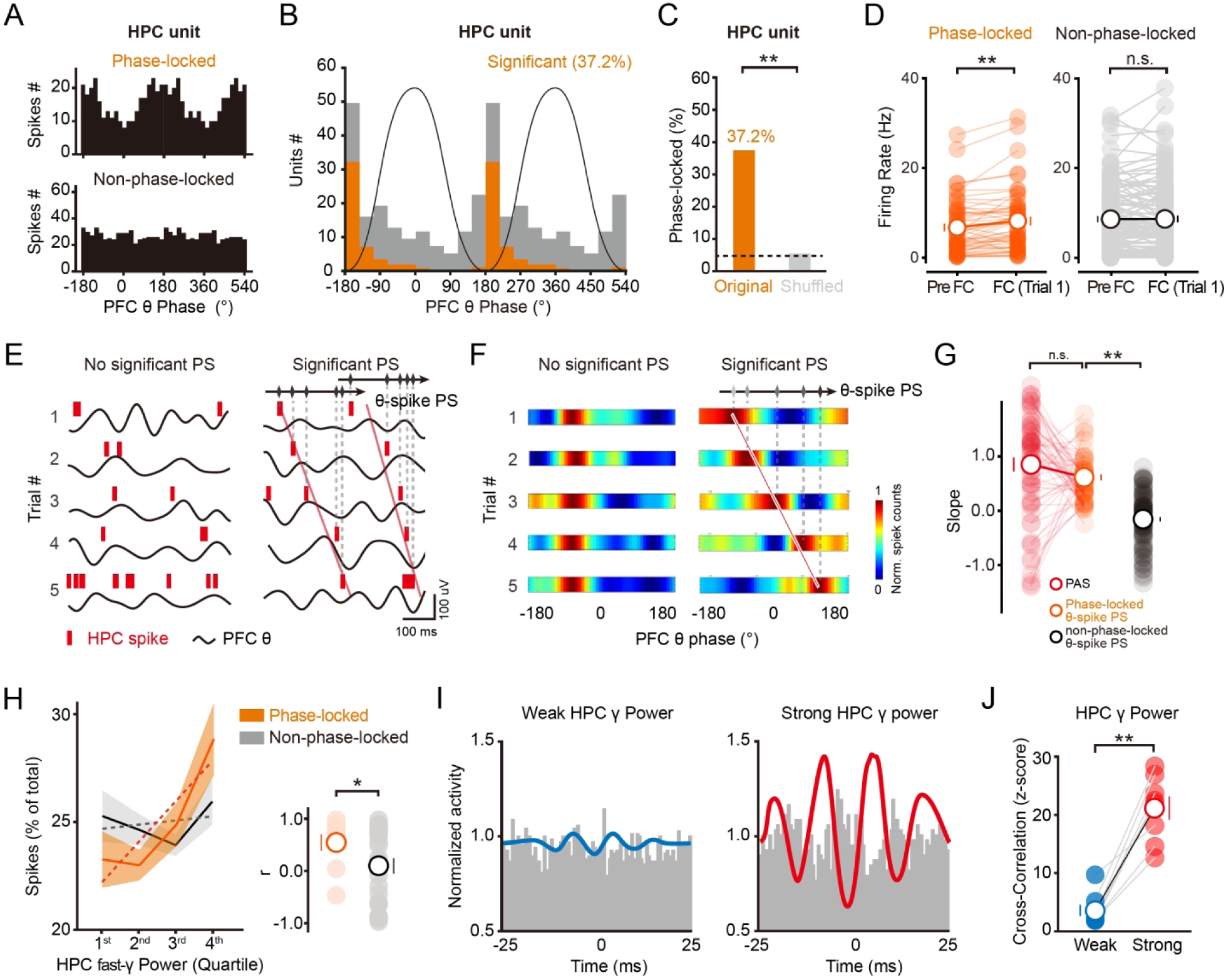
HPC– PFC coupling phase shift correlates with PFC modulation of HPC spiking. (A) Representative phases distribution histograms of single units that are phase-locked (top) or non-phase-locked (bottom) to PFC θ oscillations. Phase-locked HPC unit shows prominent modulation by PFC low-frequency oscillations. (B) Histogram of the preferred PFC θ phases for significantly (yellow) and non-significantly (gray) phase-locked HPC units. Note a prominent proportion of HPC units was phase locked to PFC θ phase (Rayleigh test for both distributions). Oscillatory cycle is repeated for clarity. (C) Percentage of HPC units phase-locked to the PFC θ oscillations. Compared to the shuffled data, original results display much higher percentage of phase-locked units. McNemar’s test, \*\**P* < 0.01. (D) The PFC θ phase-locked HPC units exhibited increased firing rate at trial 1. As a control, no such alteration was observed in other non-phase-locked units (Paired t*-*test, phase-locked HPC unit: n = 46 units from 6 rats, \**P* < 0.05; non-phase-locked: n = 104 units from 6 rats, *P* = 0.79). (E) Example units exhibiting no significant phase shift (PS; left) and significant PS (right). Units with significant PS displayed prominent circular-linear correlation of PFC θ phase with number of conditioning trials. The rhombuses and arrows at right panel indicate the preferred phase and direction of PAS, respectively. (F) Example showing progressive shift of HPC spikes preferred PFC θ phases over conditioning trials (right). In contrast, example without significant PS (left) failed to display similar trend. (G) HPC units that phase-locked to PFC θ demonstrated more prominent θ-spike PS, indicated by higher slope than non-phase-locked units (Phase-locked HPC unit: 0.43 ± 0.17, n = 46 from 6 rats; non-phase-locked: –0.10 ± 0.06, n = 104 from 6 rats; unpaired t-test, \**P* < 0.05). (H) Spike distribution as a function of HPC fast-γ power in PFC phase-locked (yellow) and non-phase-locked neurons (gray). HPC fast-γ was allocated to four evenly bins in accordance with amplitude, which was positively correlated with spike rate of both phase-locked and non-phase-locked firings. However, this correlation was significantly stronger for phase-locked firing, as indicated by the histogram on the right (average r values ± SE across cells: phase-locked, n = 46 from 6 rats, r = 0.45 ± 0.06; non-phase-locked, n = 104 from 6 rats, r = 0.22 ± 0.04; Wilcoxon rank-sum test, \**P* < 0.05). (I) Fast-γ trough-triggered histograms of spike counts. The left and right histograms correspond to HPC fast-γ oscillations at 1st and 4th quartile power levels, respectively. Colored lines, γ trough-triggered LFP. Noted that firing rate of HPC neurons were more closely correlated to strong HPC fast-γ power (red line), resulting in the exhibition of significant γ rhythms. (J) Cross-correlation of γ power triggered spikes. Significant for 8 animals; two-sided paired t-test, \*\**P* < 0.01.

Phase-locking in a subset of CA1 neurons was detected only in the first ITI. Over subsequent ITIs, the preferred phase of these neurons progressively shifted, explaining the observed phase shift despite phase-locking (**Figures 2E and 2F; Figure S6A**). Analysis before conditioning confirmed that very few CA1 neurons exhibited phase-locking to PFC theta (**Figure S6B**). Here, the θ-spike phase shift (θ-spike PS in short) was measured by computing circular-linear correlation between spike preferred phase and fear conditioning trials (**Figure 2E**). Phase-locked units showed significantly higher degree of phase shift than that of non-phase-locked neurons (**Figure 2G; Figure S6C**). The log-transformed p-values of circular-linear correlation showed consistent results (**Figure S6D**). Moreover, the firing rates of PFC theta phase-locked HPC units were positively correlated with HPC fast-γ power (**Figure 2H**). Furthermore, the activity of these HPC neurons seemed to correlated with the modulation of HPC γ power, as revealed by the γ-trough triggered distribution of HPC spiking activity (**Figures 2I and 2J**).

Neuronal spiking is less susceptible to volume conduction.^72^ Consistent with our results on the origin of PFC θ oscillations in **Figures S3**, we found that HPC neuronal spiking phase-locked to PFC θ was correlated with HPC fast γ during fear conditioning, whereas HPC neuronal spiking phase-locked to HPC θ was not (**Figure S6E-S6G**). This asymmetry provides further evidence that PFC θ is not a volume-conducted copy of hippocampal θ.

To determine the type of neurons involving in spike-θ phase shift, we classified neurons into putative pyramidal cells and interneurons using established criteria (**Figures S6H-S6J**).^84,85^ Phase-locked neurons of both cell types exhibited more prominent spike-θ phase shift compared with non-phase-locked neurons (**Figures S6K-S6P**). Together, our results suggest that the generation of PAS is correlated with PFC modulations of hippocampal spiking and fast gamma power.

### Phase shift reoccurs during memory system consolidation

The PAS over fear conditioning may reflect the initiation or early tagging of HPC-PFC interaction.^15,16,23^ As hippocampal replays in both sleeping and awake state are proposed to represent a potential neural substrate for system consolidation,^86–92^ we hypothesized that the PAS might reoccur during system consolidation of contextual fear memory. To test this, we focused on neural activities in HPC-PFC circuits over the period of system consolidation in home cage across different behavioral states (**Figure 3A**). Simultaneous recordings on EEG, EMG and autonomic signals, combined with video monitoring, enable us to distinguish different behavioral states and sleep stages (**Figure S1C**). Strikingly, cross-frequency phase-amplitude comodulograms revealed a phase-shifted coupling pattern resembling that observed during fear conditioning. This reemergence of PAS initiated at day 4 post-FC and ended around day 21 post-FC (**Figure 3B**). Notably, this phase-shifted coupling was detected specifically during NREM sleep in epochs associated with hippocampal sharp-wave ripples (SWRs, 110–250 Hz) (**Figures 3B and 3C; Figures S7, S8A, and S8B**), a network event widely implicated in memory consolidation.^93–98^

**Figure 3.**
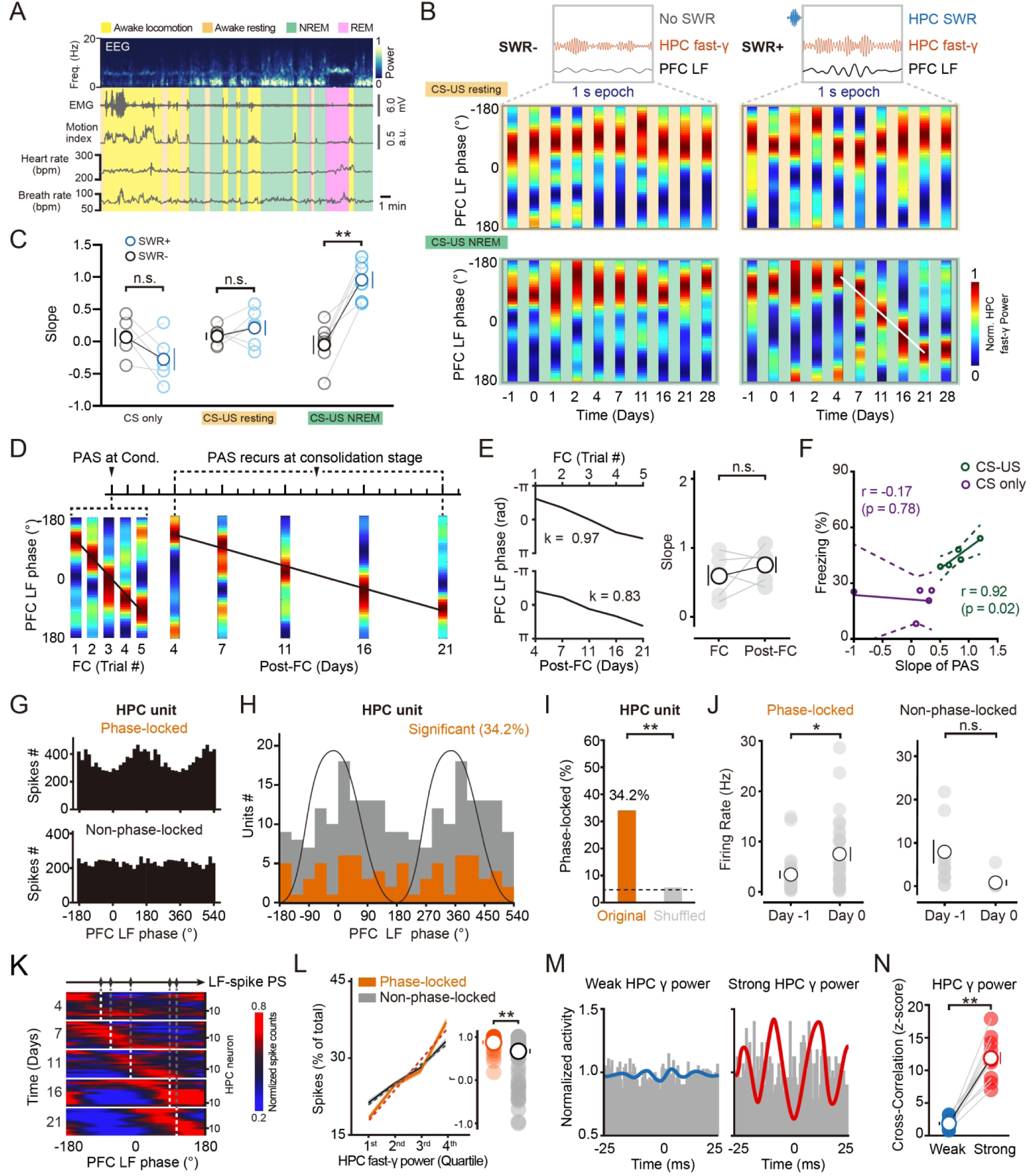
HPC-PFC coupling phase shift reoccurs during memory system consolidation. (A) Representative sample illustrating simultaneous multimodal monitoring in the home cage during memory systems consolidation. Yellow, orange, green, and pink bars indicate awake locomotion, awake resting, NREM and REM, respectively. Time-resolved power spectra and waveforms of EEG and EMG, together with motion index, blood oxygen saturation, heart rate, and respiration, exhibit distinct state-dependent amplitude profiles. (B) Cross-area comodulograms uncover the association of reoccurring PAS with HPC SWRs. The PAS observed during day 4-21 post-FC was accompanied by the occurrence of HPC SWRs during NREM sleep. LF, spindle-like low frequency activity during NREM sleep. (C) High occurrence of PAS in SWR+ CS-US NREM group during day 4-21 post-FC, as indicated by increased slope of PAS showing prominent phase shift. CS only SWR−: 0.07 ± 0.14, CS only SWR+: –0.27 ± 0.17; CS-US resting SWR−: 0.08± 0.06; CS-US resting SWR+: 0.22 ± 0.11; CS-US NREM SWR−: –0.04 ± 0.14; CS-US NREM SWR+: 0.96 ± 0.13; Repeated measures (RM) one-way ANOVA with Tukey post-hoc test, F(5,28) = 0.61, \**P* < 0.05. Mean ± s.e.m. (D) Sample shows that PAS reoccurred at consolidation stage with similar phase shifting difference to that during fear conditioning, albeit on a greatly extended time span. The sample was modified from Figure 1J **and** Figure 3B with same color-bar. (E) The PASs at conditioning and consolidation stages displayed similar phase shifting difference. The time spans of both PASs were normalized for comparing the slope at the two stages. The normalization uncovered similar phase shift distance of the two PASs (left), illustrating its consistency during conditioning and consolidation stages. Right: statistics shows no significant difference in slope of PASs at the two stages (n = 5 rats, two-sided paired t-test, *P* = 0.43, center values denote mean ± s.e.m.). (F) PAS was positively correlated with memory consolidation, as indicated by Pearson correlation coefficient of freezing percentage after conditioning and the slope of PAS (CS only: r = –0.17, CS only group, n = 6 rats, *P* = 0.78; CS-US: r = 0.92, CS-US, n = 6 rats, *P* = 0.02). Each cycle represents data from an individual animal. Colored lines show results of linear regression fit. (G-J) Same as Figures 2A**-2D**, except the results were obtained at consolidation stage (day 4-21). Similar phase-locking of HPC neuronal firing to PFC low-frequency cycle phase was observed at this stage. A subset of HPC units (I) phase-locked to the PFC low-frequency oscillations displayed increased firing rate in day 0 (J, left). As a control, no such alteration was observed in other non-phase-locked units (J, right). McNemar’s test, \*\**P* < 0.01. (K) Phase distribution of all phase-locked neurons at consolidation stage. Each row represents phase distribution of a single neuron recorded at one of the timepoints of consolidation stage. Neurons are sorted according to their preferred phase. Notice that HPC spike coupling with progressively “later” phases of the PFC low-frequency cycle. The warm colors indicate the phases that are more preferred. White lines of each day indicate the mean preferred phases. (L-N) Same as Figures 2H**-2J**, except the results were obtained at consolidation stage (day 4-21 post-FC). Both spike distribution as a function of HPC fast-γ power (L, two-sided paired t-test, t = 2.85, \*\**P* < 0.01) and γ trough-triggered histograms of spike count (M, N; two-sided paired t-test, t = 10.85, \*\**P* < 0.01) point to the notion that HPC-cortical spike-LF PS correlates with HPC γ power increment during the SWR+ 1s epochs post-FC.

Importantly, the relevant PFC low-frequency activity during these NREM epochs did not resemble a sustained, state-defining theta rhythm. Rather, it appeared as transient low-frequency events occurring in association with SWRs (**Figure S7**). On the basis of these features, we interpret them as being more consistent with sleep spindle-related cortical activity.^99–101^ Accordingly, the re-emergent PAS observed during consolidation was not a sustained feature of NREM sleep as a whole, but instead emerged selectively during SWR-associated epochs, consistent with the transient nature of the underlying PFC spindle-like events.^74^

Compared with randomly selected epochs (SWR−), 1 s epochs following the termination of SWRs (SWR+) showed more PAS events, reflected by a greater extent of phase shift and a higher percentage of cross-area channel pairs exhibiting PAS (**Figures 3B and 3C; Figures S8C and S8D**). These differences could not be explained by variation in SWR properties (**Figures S9A–S9C**). Spectral analyses time-locked to SWR events further confirmed that hippocampal SWRs and fast γ oscillations represent distinct phenomena (**Figure S9E**). In addition, no PAS was observed when HPC fast γ was replaced by ripples (**Figure S9F**), and analyses were restricted to epochs with significant low-frequency power enhancement (**Figure S8E**). PAC returned to the preexisting baseline level after conditioning and remained stable thereafter across behavioral states (**Figures S9G and S9H**). Nevertheless, the level of SWR-associated PAC was significantly increased throughout days 0-28 following training (**Figure S9I**). This training-related change in SWR-associated PAC may represent a modulation of preexisting network dynamics as a function of cognitive demand. Interestingly, although PAS during systems consolidation unfolded over weeks, the extent of phase shift was similar to that recorded during memory encoding (**Figures 3D and 3E**).

To examine whether the reported PAC is related to the 4-7 Hz PFC activity, SWRs or up and down states (UDS), we substituted PFC spindle-like low-frequency activity with HPC theta or UDS (**Figure S8F**), split the PFC low-frequency band into 4-7 Hz and 7-12 Hz (**Figures S8G-S8K**). No PAS was observed when PFC low-frequency band was replaced by hippocampal theta or UDS, whereas PAS remained evident when PFC low-frequency band was subdivided into 4-7 Hz and 7-12 Hz components. Our results indicate that PAS phenomenon does not reflect contamination by locomotion-related hippocampal theta or by PFC 4-7 Hz activity alone.

Notably, the strength of consolidated memory assessed at day 28 post-FC was highly correlated with the extent of PAS during consolidation period (**Figure 3F; Figure S8L**). These results suggest that the reoccurred PAS may represent rehearsal of formerly learned experience during fear conditioning.

To further investigate if the reoccurring PAS reflects the modulation of HPC single unit activity by PFC spindle-like low-frequency activity, we analyzed 51 well-isolated single-units from HPC. Among all the units, 34.2% displayed significant phase-locking to low-frequency activity across the consolidation phase (days 0-28 post-FC; **Figures 3G-3I**). Unlike the neurons shown in Figure 2B that are all phase-locked to PFC low-frequency activity and recorded during the first ITI of fear conditioning, the lack of a uniform phase preference in Figure 3H could be caused by the shifts in phase preference across the consolidation phase (days 0-28 post-FC). Notably, the re-emerged PAS was accompanied by the emergence of phase-locked HPC neurons, the firing of which significantly increased upon the occurrence of SWRs, while non-phase-locked neurons remained unchanged (**Figure 3J**). Considering potential electrode drifting during the longitudinal recordings, we applied average phase analysis rather than single unit phase analysis used in Figure 2.^102^ Nevertheless, we observed that HPC spike progressively coupling with “later” phases of the PFC spindle-like events (**Figure 3K; Figure S8M**). Moreover, the SWR-associated firings of these neurons were positively correlated with HPC fast-γ power (**Figure 3L**), and the phase-locked HPC neurons could be involved in the generation of HPC fast-γ oscillations (**Figures 3M and 3N**). Furthermore, we excluded the possibility that the HPC fast-γ oscillations were attributable to spectral spike leakage (**Figure S8N**). These results support the notion that reoccurring PAS is correlated with SWR-associated reactivation of HPC neuronal firings that are phase-locked to PFC spindle-like events.

### HPC-cortical coherence precedes phase shift reoccurrence

The initial appearance and subsequent recurrence of PAS occur during fear conditioning and between days 4-21 post-FC, respectively. Given the continuous nature of our observations, how do hippocampal and cortical regions interact during the intermediate period between these two phases? To address this question, we investigated the neural activities in the HPC-cortical circuits during the first week post-FC (i.e., at recent memory stage). Unlike the PAS that was only observed in the HPC-PFC circuit at learning stage, we detected a prominent cross-area oscillatory coherence between HPC and each of the three cortical regions. Notably, the HPC-cortical coherence selectively occurred at fast-γ band associated with SWR+ in conditioned CS-US animals (**Figures 4A and 4B; Figures S10A and S10B**). Specifically, compared to CS only group, the conditioned animals demonstrated increases in both coherence magnitude and percentage of channel pairs displaying γ coherence (**Figures 4C and 4D; Figure S10C**). This cross-area γ coherence initiated at day 0 and completely disappeared at day 7 post-FC. Notably, no such alterations in low-frequency coherence were observed (**Figure S10D**). Moreover, the γ coherence increase was accompanied by γ power increase selectively in the time window related to SWRs (within 1-second epoch; **Figure 4E**). Notably, this transient γ power increase did not cause the overall γ power change (**Figure S10E**), suggesting that the HPC-cortical fast-γ coherence is related to SWRs-associated fast-γ power increase.

**Figure 4.**
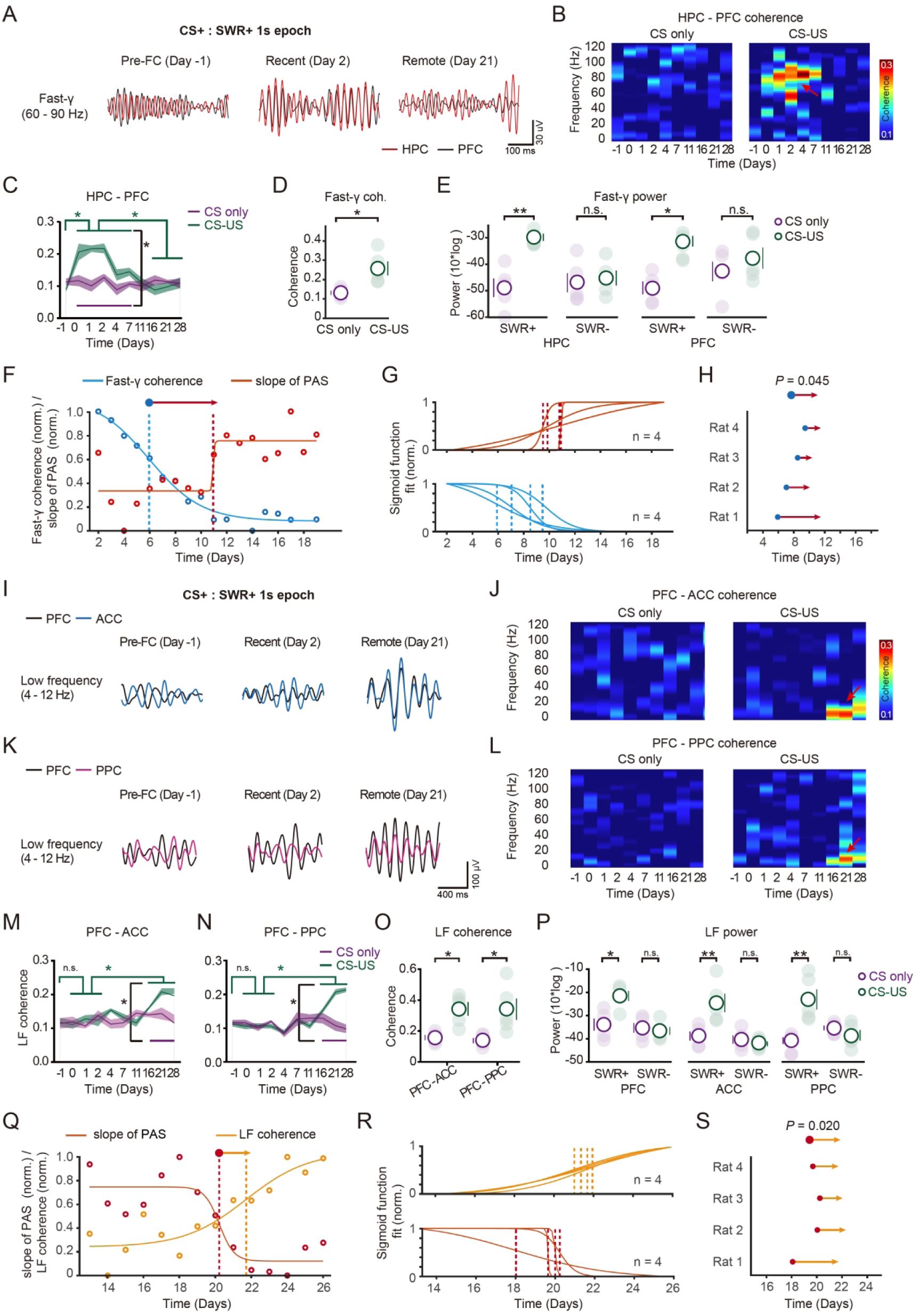
Phase shift reoccurrence bridges HPC-cortical and intercortical network coordination. (A) Overlaid examples of the filtered LFP at fast-γ band in HPC (black) and PFC (red) during 1s epochs following SWRs (SWR+) at day –1 (one day before FC), day 2 (recent stage) and day 21 (remote stage) post-FC. Note LFP coherence at day 2 post-FC. (B and C) Representative day-resolved time-frequency spectrums of CS only group (B, left) and CS-US group (B, right) and average cross-area γ coherence spectrum (C) revealed considerable HPC-PFC γ coherence at recent memory stage in CS-US group, but not in CS only group. Mean γ coherence (mean ± s.e.m.) was quantified in CS-US group (green; n= 6 rats) and CS only group (magenta; n= 6 rats). Bonferroni post-hoc test, \**P* < 0.05. CS-US group versus CS only group from day 0 and day 7. (D) Histograms show increased HPC-PFC γ coherence at recent stage in conditioned CS-US group. Mean γ coherence ± s.e.m. of 6 rats from CS only group (magenta), 6 rats from CS-US group (green), Bonferroni post-hoc test, \**P* < 0.05, CS-US group versus CS only group from d0 to d7, center values denote mean ± s.e.m.. (E) The γ power in both HPC and PFC increased selectively upon the occurrence of SWRs (within SWR+ 1s epoch) at recent stage. Unpaired t-test, n = 6 rats, \**P* < 0.05, \*\**P* < 0.01, n.s. no significant, center values denote mean ± s.e.m.). (F) Representative sigmoid function fits from a single animal showing a gradual increase in the slope of PAS alongside a gradual decrease in HPC-PFC γ coherence. (G) Sigmoid function fits for each animal (n = 4). (H) Sharpest changes marked by blue circles and red arrowheads. Significant for 4 animals; two-sided paired t-test, *P* = 0.045. (I and K) Overlaid examples of filtered low-frequency LFP at 4-12 Hz band recorded in PFC (black), ACC (blue) and PPC (pink) during 1s epochs following PFC cortical SWRs (SWR+) at day –1 (one day before FC), day 2 (recent stage) and day 21 (remote stage) post-FC. Note prominent low-frequency coherence at day 21. (J and L) Representative day-resolved time-frequency spectrums of CS only group (left) CS-US group (right) revealed considerable PFC-ACC (J) and PFC-PPC (L) low-frequency coherence at remote memory stage. (M and N) Enhanced low-frequency coherence in PFC-ACC (M) and PFC-PPC (N) circuits at remote stage, revealed by quantification of mean coherence across all the timepoints. Mean low-frequency coherence (mean ± s.e.m.) was quantified in CS-US group (green; n= 6 rats) and CS only group (magenta; n= 6 rats). Bonferroni post-hoc test, \**P* < 0.05. CS-US group versus CS only group from day 21 and day 28. LF, low frequency. (O) Histograms show increased low-frequency coherence in PFC-ACC and PFC-PPC circuits at remote stage (unpaired t-test, n = 6 rats, \**P* < 0.05, center values denote mean ± s.e.m.). (P) The power of low-frequency activity in PFC, ACC and PPC increased selectively upon the occurrence of SWRs (within SWR+ 1s epoch) at remote stage. (Unpaired t-test, n = 6 rats, \**P* < 0.05, \*\**P* < 0.01, n.s. no significant, centre values denote mean ± s.e.m.). (Q) Representative sigmoid function fits from a single animal showing a gradual increase in PFC-ACC low-frequency coherence alongside a gradual decrease in the slope of PAS. (R) Sigmoid function fits for each animal (n = 4). (S) Sharpest changes marked by red circles and orange arrowheads. Significant for 4 animals; two-sided paired t-test, *P* = 0.020.

We then formally tested the temporal relationship between HPC-cortical gamma coherence and PAS. To this end, we fit a sigmoid function to each animal’s cross-area coupling over time (**Figures 4F and 4G; Figures S10F-S10J** and Methods).^22^ This analysis revealed a series of robust stepwise events that the decrease in HPC-cortical gamma coupling preceded the PAS reoccurrence. This lag was significant across the animals (**Figure 4H**). Similar results were obtained when Pearson’s correlation coefficient (R^2^) was calculated^73^ to assess the goodness-of-fit value of PAS in phase-trial regression (**Figures S10F-S10H**). These findings indicate that the disengagement of SWR-associated HPC-cortical fast-γ coherence precedes and predicts the subsequent PAS engagement.

### Inter-cortical coherence follows phase shift reoccurrence

To further characterize interregional oscillatory crosstalk following PAS reoccurrence, we tracked the evolution of neural dynamics toward the remote memory stage (day 14-28 post-FC). Multi-site recordings obtained in home cage at different timepoints after fear learning (day 16, 21, 28 post-FC) revealed a steady increase in oscillatory coherence between PFC-ACC and PFC-PPC at the low frequency band (4-12 Hz) associated with SWR+ (**Figures 4I-4L; Figure S11A**). This increase in inter-cortical coherence started during D11-D16 post-FC and lasted over the remote stage till day 28 post-FC (**Figures 4M-4O; Figure S11B**). Notably, PFC-ACC and PFC-PPC coherence remained significantly increased when cortical low-frequency activity was subdivided into 4-7 Hz and 7-12 Hz components (**Figure S11C**). In contrast, fast-γ band did not show similar significant coherence (**Figure S11D**), indicating a specific mode of interaction. In addition, the SWRs-associated cortical low-frequency power was significantly increased during the remote stage (**Figure 4P**), while the overall low-frequency power was unchanged compared to that in the CS only group (**Figure S11E**) and across behavioral states (**Figure S11F**). Notably, with a refined classification of awake locomotion, low-frequency coherence between the PFC and HPC was found to be selectively enhanced during high-speed locomotion, exhibiting significant differences relative to other states (**Figure S11F; Table S1**). Furthermore, no increase in SWR-related low-frequency coherence was observed between ACC and PPC, suggesting PFC could be the network hub to organize the timing of low-frequency events in other cortical regions for cross-area synchrony (**Figure S11G**).

To characterize the temporal relationship between PAS and inter-cortical coherence, we fitted a sigmoid function to each animal’s cross-area coherence (**Figures 4Q and 4R; Figures S11H-S11L** and Methods).^22^ We found that during day 14-21 post-FC, a decrease in slope of PAS was followed by a sharp rise in PFC-cortical low-frequency coherence. The lag of decrease in slope of PAS and the rise in PFC-cortical coherence allowed us to estimate the transition across the animals (**Figure 4S**). Similar results were obtained when Pearson’s correlation coefficient (R^2^) was calculated^73^ to assess the goodness-of-fit value of PAS (**Figures S11H-S11J**). These analyses support the notion that disengagement of PAS predicts the subsequent occurrence of inter-cortical coherence.

Together, our results uncovered HPC-cortical coherence, phase shift and inter-cortical coherence that successively occurred in HPC-cortical network during remote memory formation. Notably, these neural dynamics were temporally overlapped and associated with different stages of remote memory formation. The “hand-in-hand” relaying style of these events helps to constitute a continuum of coordinated cross-area oscillatory dialogues potentially crucial to memory consolidation and cortical stabilization.

### Shift of HPC-cortical dialogue organizer and directionality

The above observations collectively suggest that PAS is temporally associated with the transition from HPC-cortical to inter-cortical dialogue. During the transition, HPC and PFC are potential dialogue organizers, who respectively play dominant roles in orchestrating HPC-cortical and intercortical coherence. To test this possibility, we first ask how the HPC-cortical γ coherence occurred at recent memory stage. Through multi-site recordings, we observed HPC fast-γ trough-triggered LFPs at PFC, ACC, and PPC (**Figure 5A**). Specifically, strong fast-γ-band synchronization (>1.5 S.D. above mean power) between HPC and all three cortical regions was observed. Similarly, based on changes in low-frequency coherence between PFC and other two cortical regions, we studied the possible dominant region at remote memory stage. During the epochs of strong PFC low-frequency oscillations (>1.5 S.D. above mean power),^51^ we found that PFC oscillations were phase-synchronized across cortical structures, as shown in PFC low-frequency cycle-trough-triggered LFP (**Figure 5B**). Our results suggest that at recent stage HPC fast-γ oscillation affects the temporal distribution of γ oscillations in the cortical regions, while at remote stage PFC low-frequency oscillation affects the temporal distribution of low-frequency activity in other two cortical regions.

**Figure 5.**
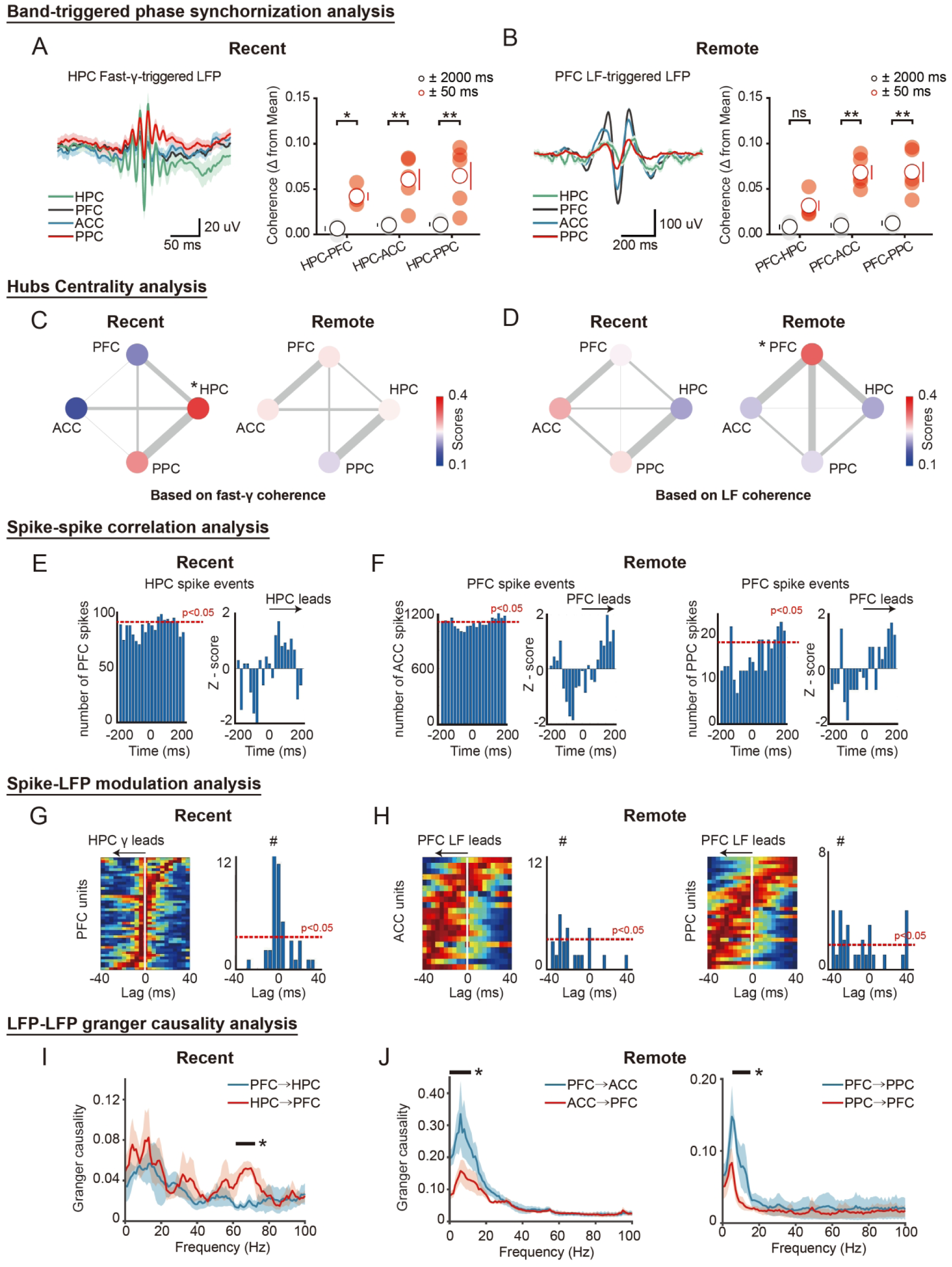
Shift of HPC-cortical dialogue organizer and temporal directionality during memory consolidation. (A and B) Trough-triggered LFP analysis reveals shift of dialogue organizer from HPC to PFC. At recent stage, significant coherence of fast-γ trough-triggered LFP was observed between HPC and cortical areas (A), while at remote stage significant coherence of LF band trough-triggered LFP was observed between PFC and other cortical areas (B). Right panel, histograms showed significantly higher coherence of trough-triggered LFP near 0 (± 50 ms) than that near 0 (± 2000 ms) in each group. Unpaired t-test, n = 6, \**P* < 0.05, \*\**P* < 0.01. LF, low frequency. (C and D) Shift of hub centrality from HPC to PFC. HPC and PFC exhibited high centrality scores selectively at recent and remote memory stage, respectively. Centrality scores were depicted by γ coherence (C) or LF coherence (D) between HPC, PFC, ACC and PPC measures that compose each network, inferred from the data in Figure 4, respectively. Both γ and low-frequency coherence measures were depicted by the lines connecting the brain regions (n = 5). (E and F) Spike-spike correlation in HPC-PFC (E) and PFC-cortical (ACC, PPC) circuits (F). The cross-area spike correlogram showed high correlation between 0 and 200 ms, inferring that HPC units preceded PFC units at recent stage and PFC units preceded other cortical units at remote stage. 201 in 340 HPC-PFC spike pairs showed significant HPC-leading interactions (E), while 53 in 108 PFC-ACC spike pairs and 47 in 120 PFC-PPC spike pairs showed significant PFC-leading interactions (F). Red dashed lines indicate significance level estimated by spike jitter test (*P* < 0.05). (G and H) Pseudo color plot of normalized pairwise phase consistency values. PFC units significantly phase-locked to HPC γ sorted by lag of maximal phase-locking at recent stage (G). By contrast, ACC and PPC units significantly phase-locked to PFC low-frequency activity at remote stage (H). Hash indicates mean lag. Right columns in G and H refer to mean normalized PPC value by lag. The mean strength of PFC unit phase-locking to HPC γ was maximal at lags in which HPC leads, whereas the mean strength of ACC and PPC unit phase-locking to PFC low-frequency cycle was maximal at lags in which PFC leads. Vertical red line, zero lag. Hash indicates mean lag. Horizontal red dashed lines, chance. (I and J) Granger causality analysis based on LFPs. HPC γ oscillations lead HPC LFPs at recent stage (I; n = 5 rats, paired t-test, HPC → PFC versus PFC → HPC: t (9) = 2.58, \**P* < 0.05), whereas PFC low-frequency oscillations lead ACC and PPC LFPs at remote stage (J; n = 5 rats; PFC → ACC versus ACC → PFC: t (9) = 5.91, PFC → PPC versus PPC → PFC: t (9) = 7.63, multiple t-tests, Black lines indicate \**P* < 0.05).

Furthermore, we performed graph-based analysis of the correlations between brain regions.^103^ The weight assigned to each brain region is determined by the strength of coherence between regions, with stronger coherence corresponding to a higher weight, whereas the hubs centrality score quantifies the total connection weight between a specific brain region and all other regions, providing a measure of its overall network connectivity. We found that at recent memory stage HPC has the highest centrality among the four brain region nodes (HPC: 0.357, Average of others: 0.214) (**Figure 5C**), whereas at the remote stage the centrality of PFC was significantly higher than that of the other three brain regions (PFC: 0.331, average of others: 0.223) (**Figure 5D**).

Pharmacological experiments with CNQX demonstrated that the HPC and PFC play a causal role in coordinating oscillations at distinct stages of memory. During the recent memory phase, CNQX injection into the HPC disrupts HPC-cortical fast-γ coherence, whereas during the remote memory phase, CNQX injection into the PFC impairs low-frequency coherence between PFC and other cortical regions (**Figure S12**). Thus, HPC and PFC are the central nodes during system consolidation, and the successive modulation by HPC fast-γ oscillation and PFC low-frequency activity may respectively contribute to the coherence in HPC-cortical and PFC-cortical circuits occurred at different stages.

This shift in the dialogue organizer may accompany or lead to a change in the direction of information transfer. We thus further analyzed temporal directionality of cross-regional dialogues in HPC-cortical circuit. Firstly, we measured precisely timed spiking offsets between regions. We found that HPC units led PFC (**Figure 5E**), ACC and PFC units (**Figure S13A**) at the recent stage, whereas PFC units led both the ACC and PPC units at the remote stage (**Figure 5F**). These findings suggested that HPC leads the cortical regions at recent memory, while PFC lead the other two cortical regions at remote memory.

Additionally, we conducted a lag analysis across the four regions. We found that a subset of PFC units was significantly phase-locked to HPC gamma oscillations (**Figure S13B**), and the distribution of phase-locked PFC unit lags were skewed in the direction where HPC gamma preceded PFC spikes (**Figure 5G**). Similar observations were found with ACC units and PPC units (**Figure S13C**). As to the remote memory stage, we determined that a subset of ACC and PPC units were significantly phase-locked to PFC low-frequency events, and the distribution of phase-locked ACC and PPC unit lags were skewed in the direction where PFC low-frequency activity preceded ACC and PPC spikes (**Figure 5H**).

Finally, we calculated Granger causality index (GCI) between pairs of brain regions. At recent stage, we found significant Granger causality in fast gamma band for the HPC→PFC direction than that for the PFC→HPC direction (**Figure 5I**). Similar phenomena were observed in HPC→ACC and HPC → PPC directions (**Figures S13D and S13E**). In sharp contrast, at remote stage we found significant Granger causality at PFC → ACC and PFC → PPC direction in low-frequency band (**Figure 5J**).

Together, these findings reveal the shift of cross-regional dialogue organizer (from HPC to PFC) and the corresponding transition of temporal directionality (from HPC→PFC to PFC→ACC and PFC→PPC) over the system consolidation of remote fear memory.

### Reoccurring phase shift mediates HPC-to-PFC transition during consolidation

PAS reoccurred upon both the transition from recent to remote memory and the shift of dialogue organizer and temporal directionality. The temporal coincidence raises the possibility that PAS is the physiological substrate that mediates memory transfer from HPC to PFC. To test this hypothesis, we injected recombinant adeno-associated viruses (rAAV) encoding the opsin (eNpHR) bilaterally into PFC to specifically express eNpHR in PFC neurons and implanted optic fiber just above PFC for optogenetic silencing of PFC neurons (**Figures 6A and 6B**). Then we carried out a stimulation protocol in which photo-stimulation was randomly delivered to disrupt the phase of PFC low-frequency at day 4-21 post-FC.^104^ This protocol had random distributions of inter-stimulation intervals ranging from 83 to 250 ms to ensure the phase intervention was operated at frequency of 4-12 Hz (**Figure 6B**). The efficacy of intervention was validated by optogenetic illumination of PFC neurons, which elicited transient disturbance of LFP and suppression of neuronal spiking (**Figures 6C and 6D**). Notably, this random intervention did not induce persistent change in PFC firing rate (**Figure S14A**). Through above opto-stimulation, we disrupted the low-frequency rhythm in the PFC, which weakened the reliability of HPC γ power peaked at a particular PFC low-frequency cycle phase in the control group (**Figures 6E and 6F**). We noted the strength of PAC before intervention was lower than that during fear conditioning. This could be due to the fact that PAC was assessed under offline state during consolidation. Over the period of PAS reoccurrence at day 4-21 post-FC, we continuously performed 2-hour optogenetic manipulations at the same time each day.^105^ The analysis of the results revealed reduction in the PAC intensity and erasure of the PAS in the conditioned animals (**Figure 6G; Figures S14B and S14C**). Moreover, the subsequent increase of inter-cortical coherence at the remote memory stage was absent (**Figure 6H; Figure S14D**). Compared to baseline data, the low-frequency phase interventions had no significant effect on the properties of SWRs (**Figures S14E-S14G**).

**Figure 6.**
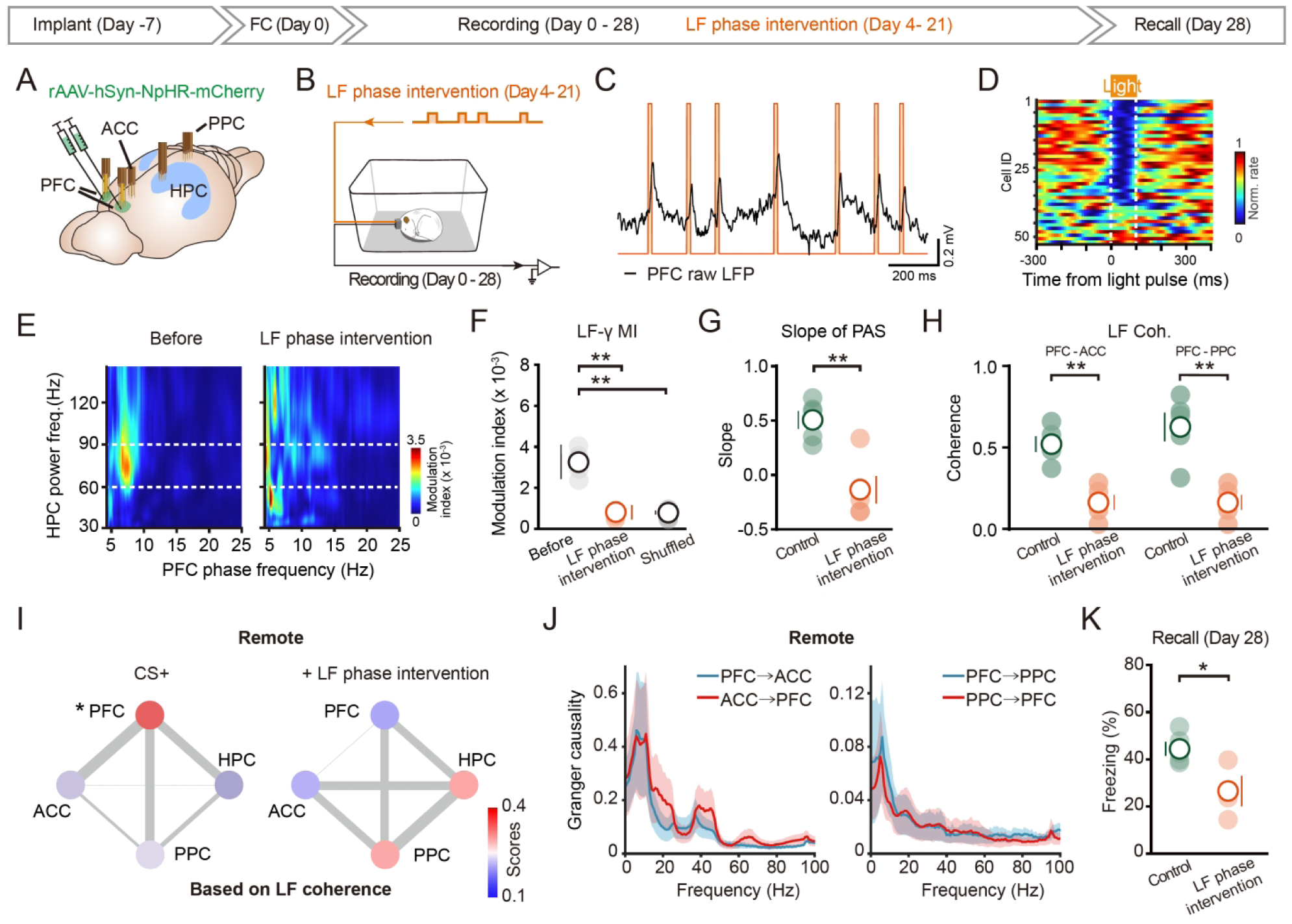
Reoccurring phase shift mediates HPC-to-PFC shift during consolidation. (A) Experimental schematics showing electrodes targeting and viral injection site. rAAV carrying eNpHR was stereotactically injected into PFC (PL of PFC) to specifically express eNpHR in PFC neurons. (B) Experimental schedule and random optogenetic intervention strategy. low-frequency pulse was randomly injected to PFC to disturb local low-frequency oscillation during day 4-21 post-FC. LF, low frequency. (C) The efficacy of low-frequency phase intervention is illustrated with a raw data example. (D) Example spiking activity of PFC neurons with respect to light delivery for LF phase intervention. 37 in 54 neurons were inhibited. (E and F) Representative Phase-amplitude comodulograms (E) and statistical histogram (F) shows significantly decreased PAC modulation index (MI) in HPC-PFC circuit of conditioned animals (Control: n = 5, 3.2 ± 0.28 x 10^-3^; LF phase intervention: n = 5, 0.82 ± 0.12 x 10^-3^, paired t-test, \*\**P* < 0.01). (G) Random LF intervention suppresses reoccurrence of PAS, indicated by decreased slope of PAS (Control: n = 5, 0.51 ± 0.08, LF phase intervention: n = 5, –0.14 ± 0.12, unpaired t-test, \*\**P* < 0.01). (H) Significantly decreased low-frequency (4-12 Hz) coherence in PFC-ACC and PFC-PPC circuits at remote stage (unpaired t-test, n = 5, \*\**P* < 0.01). (I) The shift of hub centrality from HPC to PFC was absent following random low-frequency intervention. In contrast to conditioned animals without LF phase intervention (left), PFC failed to exhibit centrality scores at remote memory stage following random LF intervention (right). (J) The shift of temporal directionality in HPC-PFC circuit was absent following random low-frequency intervention. Granger causality analysis based on LFPs shows no PFC LF oscillations leading ACC or PPC LFPs were detected at remote stage (n = 5 rats; PFC → ACC versus ACC → PFC: t (9) = 1.01, PFC → PPC versus PPC → PFC: t (9) = 0.30, multiple t-tests, *P* > 0.05). (K) Random LF intervention suppressed memory recall at remote stage (day 28 post-FC; Control, n = 5, 44.4 ± 2.8; LF phase intervention, n = 5, 29.9 ± 6.0, unpaired t-test, \**P* < 0.05).

Notably, the above PAS suppression was accompanied by failure in coupling organizer and directionality shift in HPC-cortical circuits (**Figure 6I**). Specifically, the transitions of highest centralization score (from HPC to PFC) and Granger causality (from HPC→PFC to PFC→ACC and PFC → PPC), were absent (**Figure 6J**). As a result, the animals subjected to low-frequency phase intervention showed a dramatic reduction in retention of remote contextual fear memory (**Figure 6K**). These results suggest that reoccurring PAS during system consolidation is required for the organizer/directionality shift and subsequent inter-cortical coherence at remote stage.

As the reoccurring PAS was closely associated with SWRs, we further examined if SWRs was required for the reoccurrence of PAS. Using the closed-loop intervention technique similar to former investigations,^92,94,105–108^ we found that the suppression of SWRs dramatically decreased the reoccurrence of PAS (**Figure S15**), suggesting that SWRs-associated hippocampal activity is necessary for the reoccurrence of PAS during consolidation.

### Causal role of HPC-cortical coherences in memory formation and stabilization

The above data indicate that HPC low-frequency events is necessary for the emergence of PAS and successful system consolidation. To further pinpoint the intervention time window within HPC-cortical interactions, we developed a closed-loop optogenetic intervention technique to silence neurons upon detection of the cross-area coherence (**Figure 7A**).^109,110^ Specifically, we monitored real-time LFPs and calculated cross-area coherence during the experiments. A 400-ms time window was applied to perform Fourier Transform and subsequent coherence analysis on the filtered LFPs. rAAV carrying eNpHR was specifically expressed in dorsal HPC bilaterally for optogenetic silencing of neurons based on the coherence between the HPC and all three cortical regions (**Figure 7B**).

**Figure 7.**
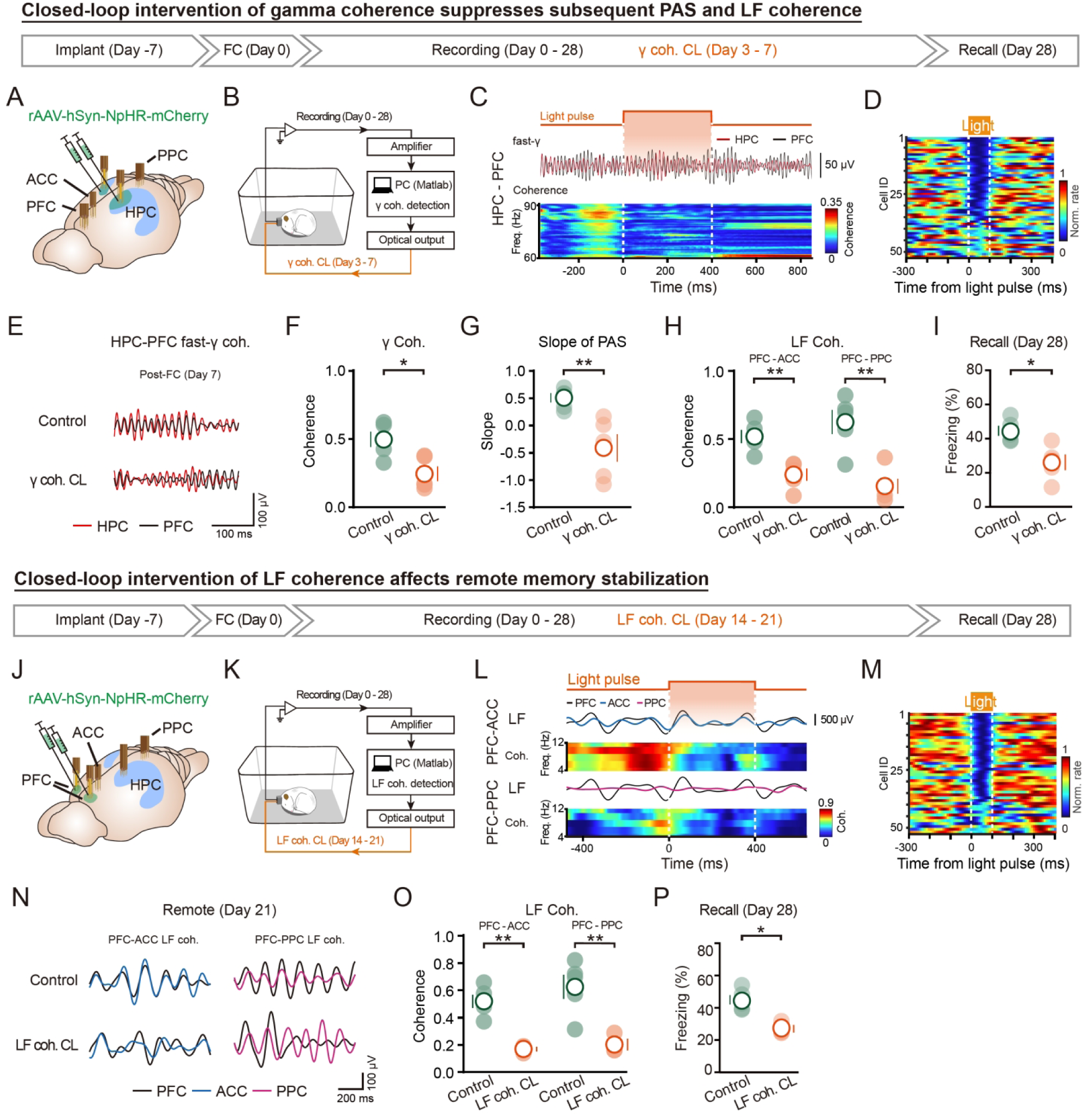
Causal links of cross-area coherences to remote memory formation and stabilization. (A) Electrodes targeting and daily experimental schedule. (B) Closed-loop intervention strategy targeting HPC-cortical γ coherence via optogenetic deactivation of HPC neurons at day 3-7 post-FC of recent memory stage. (C) Validation of real-time efficiency of closed-loop intervention. Examples of filtered LFPs and HPC-PFC coherence spectrum demonstrate suppression of HPC-PFC γ coherence upon closed-loop intervention (γ coh. CL, indicated by yellow shaded area). (D) Example spiking activity of HPC neurons with respect to light delivery for γ coh. CL. 37 in 51 neurons were inhibited). (E) Examples of filtered fast-γ LFPs recorded after γ coh. CL (day 7 post-FC) in HPC (red) and PFC (black). (F and G) Histograms show γ coh. CL implemented at recent stage suppress γ coherence and subsequent reoccurrence of PAS. Following γ coh. CL, both the increase in HPC-PFC γ coherence (F; n = 5, \**P* < 0.05, n.s. no significant, unpaired t-test) and PAS reoccurrence (G; control: n = 5, 0.51 ± 0.08, γ coh. CL: n = 5, –0.41 ± 0.25, unpaired t-test, \*\**P* < 0.01) were absent. (H) Histograms show the increase in LF coherence at remote stage was absent in HPC-ACC and HPC-PPC circuits following γ coh. CL (unpaired t-test, \*\**P* < 0.01). LF, low frequency. (I) ã coh. CL impaired memory retention and stabilization, indicated by decreased freezing behavior upon remote memory recalled at day 28 post-FC (Control: n = 5, 44.4 ± 2.8; γ coh. CL: n = 5, 26.0 ± 4.5, unpaired t-test, \**P* < 0.05). (J-M) Same as A-E, except inter-cortical low-frequency coherence CL (LF coh. CL) was implemented via optogenetic deactivation of PFC neuron during day 14-21 post-FC. Validation of LF coh. CL were displayed by examples of the filtered low-frequency LFPs recorded in PFC (black), ACC (blue) and PPC with or without CL (L), as well as the example spiking activity of PFC neurons with respect to light delivery for low-frequency coh. CL. In M, 50 in 68 neurons were inhibited. (N) Examples of filtered low-frequency LFPs recorded after LF coh. CL (day 21 post-FC) in PFC (black), ACC (blue) and PPC (pink). (O-P) Low-frequency coh. CL suppressed inter-cortical coherence (N, O; \*\**P* < 0.01, unpaired t-test) and impaired subsequent memory stabilization at remote stage (P; Control: n = 4, 44.4 ± 2.8, LF coh. CL: n = 4, 27.1 ± 2.3, unpaired t-test, *P* < 0.05).

Using this custom-developed closed-loop intervention technique, we investigated if the γ coherence detected in HPC-cortical circuit was causally involved in remote fear memory formation, using 2-hour optogenetic stimulation at the same time each day for consecutive seven days.^105^ We confirmed that closed-loop 561 nm illumination of HPC neurons disturbed γ coherence and inhibited spiking in vivo in a temporally precise and stable manner (**Figures 7C and 7D; Figure S16**). To avoid affecting the recent memory,^93^ we only conducted the optogenetics experiment at day 3-7 post-FC in HPC-PFC circuit. Indeed, optogenetic intervention at this time period failed to exhibit any influence on recent memory formation (**Figure S17A**). Compared to baseline data, the gamma coherence interventions had no significant effect on the properties of SWRs (**Figures S17B-S17D**). At the end of this optogenetic silencing of CA1 neurons at day 7 post-FC, the magnitude of γ coherence and the percentage of cross-area channel pairs exhibiting increased γ coherence were dramatically lower than that in animals without such intervention (**Figures 7E and 7F; Figures S17E and S17F**). Strikingly, the reoccurrence of subsequent PAS in HPC-PFC circuit and inter-cortical coherence were all erased. Notably, this interruption of HPC-PFC oscillatory dialogue was accompanied by impaired remote memory recalled at day 28 post-FC (**Figures 7G-7I; Figures S17G and S17H**). Moreover, control experiments using random γ delayed intervention showed no effect (**Figures S17I-S17L**). These results suggest chronic closed-loop intervention over several days at recent memory stage engenders enduring effects on both γ coherence and subsequent events.

Considering the early interaction between HPC and PFC at conditioning stage, as well as the crucial role of HPC and PFC in the transition of cross-dialogue organizer, we were motivated to further investigate whether the coherences in HPC-ACC and HPC-PPC circuits were independent or secondary to HPC-PFC γ coherence. Thus, we performed closed-loop inhibition of PFC principal neurons only based on the coherence between the HPC and PFC, rather than on the γ coherence between the HPC and all the three cortical regions (**Figures S17M and S17N**). We found that this closed-loop intervention abolished the γ coherence increase between HPC and the three cortical regions as before (**Figure S17M**) and resulted in fear recall deficits (**Figure S17N**), suggesting that the γ coherence in HPC-ACC and HPC-PPC circuits are not independent events, but secondary to HPC-PFC γ coherence. Collectively, these results indicate that HPC-PFC coherence is a prerequisite for the subsequent reoccurrence of PAS and inter-cortical coherence, thus reinforce the notion that the successive HPC-cortical events are interconnected.

We next examined the role of inter-cortical coherence that occurred at remote stage. As PFC is the dominant region that proved to organize the coherence with other cortical regions, we performed optogenetic silencing of PFC neurons upon detecting of inter-cortical coherence (**Figures 7J and 7K**). We found the closed-loop optogenetic suppression of PFC neurons based on inter-cortical coherence detection during remote stage could dampen the occurrence of inter-cortical coherence and neural spiking (**Figures 7L and 7M**). Compared to baseline data, the low-frequency coherence interventions had no significant effect on the properties of SWRs (**Figures S17O-S17Q**). Interestingly, the suppressive effect could sustain even after the termination of the closed-loop intervene, as the occurrence of inter-cortical coherence was still low even at the end of our recording at day 28 post-FC (**Figures 7N and 7O; Figures S17R and S17S**). As a result, the retention of remote memory was impaired (**Figure 7P**), indicated by decreased freezing behavior upon remote memory retrieval. Moreover, control experiments using random delayed intervention showed no effect (**Figures S17T and S17U**). These results suggest that inter-cortical coherence is required for stabilization and maintenance of remote memory. Together, our findings prove the causal role of the coordinated cross-area oscillatory coherences in formation and stabilization of remote fear memory.

## DISCUSSION

Memory system consolidation involves cross-area communication between hippocampus and cortical regions.^6,29,53^ We have observed a dynamic temporal sequence of distinct types of HPC-cortical interactions during the formation of remote memory (**Figure S18A**). This sequence starts with the occurrence of PAS over training sessions, possibly reflecting the early tagging or index of learned information from hippocampus to PFC. This is followed by an increase in HPC-cortical fast-γ coherence, potentially initiating system consolidation of remote memory. Then PAS reoccurs, extending across day 4-21 post-FC, likely sustaining the ongoing consolidation and transferring the consolidated memory to cortex as discussed below. This is followed by an increase in inter-cortical low-frequency coherence, possibly stabilizing the consolidated memory in cortex. Notably, these events are interconnected rather than independent. Each event precedes and predicts the next one. When the former event gradually faded away, the later one initiates before the former one totally disappeared. This “hand-in-hand” relaying style helps to constitute a chain of cross-area oscillatory coupling events potentially crucial to memory consolidation and cortical stabilization. As brain oscillations are known to modulate neuronal firing,^6,29,53^ they may be important in generating the precise neuronal firing, particularly related to synaptic plasticity heavily relying on spike timing.^111–113^

System consolidation is a gradual and time-consuming process. To the best of our knowledge, the present study described for the first time a gradual coupling phase shift in the HPC-cortical circuit throughout the process of system consolidation. Importantly, intervention of the PAS hindered the remote memory consolidation, providing direct evidence for the causal link between PAS and memory consolidation. Consistent with this view, we found that PAS is associated with the occurrence of SWRs and reactivation of phase-locked HPC neurons, which are likely key neurophysiological markers of memory consolidation. Furthermore, the time window of PAS reoccurrence is temporally linked with the cross-regional coherences observed at both recent and remote stages, making it a feasible substrate linking two stages for memory system consolidation. The progressive feature of reoccurred PAS may represent a neural signature of the gradual HPC-to-cortical memory transfer process.

Memory stabilization and integration during sleep are thought to rely on the reactivation of awake neural patterns, which facilitate synaptic modifications across hippocampal-neocortical networks.^48,49^ During the NREM phase of sleep, SWRs serve as a critical window for memory-related activity. SWR replay during sleep gradually consolidates memory traces within the broader context of existing memories. If SWR replay assemblies propagate mnemonic information across distributed brain circuits, a significant portion of brain activity should be temporally aligned with SWRs. In line with this, during NREM sleep in the systems consolidation phase, we observed the re-occurrence of PAS following SWR epochs. Our results suggest that both the encoding and systems consolidation of episodic memories are supported by a phase coding mechanism. This phenomenon may represent a versatile brain function that enables the gradual information transmission through inter-regional couplings.

A more specific implication of this interpretation is that the prolonged time window of PAS reoccurrence during consolidation likely reflects repeated recruitment of many brief replay-linked coordination events, rather than maintenance of a continuous phase-shifted state. Sleep spindles are transient thalamocortical events,^99–101^ and hippocampal SWRs are likewise brief replay-associated events.^114–116^ Accordingly, the consolidation-stage PAS described here is unlikely to represent a continuously maintained phase-shifted pattern throughout NREM sleep. Instead, it is more plausibly viewed as a temporally structured interaction that reappears transiently during discrete SWR-associated spindle windows and is repeatedly engaged across days 4–21 post-FC. In this framework, the extended duration of PAS reoccurrence reflects not a sustained oscillatory state, but the cumulative effect of many brief replay-associated events that progressively support systems consolidation.

Recent memory traces initially are stored in HPC-cortical connections. As the memory mature and consolidated into remote memory, they become stored in strengthened connections across distributed cortical regions. Our findings on the transition of cross-regional dialogue organizer and directionality further support this notion.^7,19^ The hippocampus and PFC alternately play dominant roles at different stages of remote memory formation. At learning stage, PFC modulates HPC neuronal spiking for the generation of PAS. This is followed by HPC and PFC successively orchestrating HPC-cortical and inter-cortical coherence at recent and remote stages. Pharmacological blockades of neural activities in HPC and PFC further confirmed these observations. The PAS reoccurred upon this recent to remote memory transition and recapitulate the PFC modulation of HPC neuronal firing.

Consistent with the findings on shift of dialogue organizer, a shift of the temporal directionality of cross-area coherent activity was also revealed using multiple analysis methods. All the analyzing results consistently demonstrate a shift of information flow directionality from HPC→PFC to PFC →ACC or PPC. It is worth mentioning that the dialogue between HPC and cortical regions may not necessarily be achieved through direct projections.^30–32^ Other regions, such as the anteromedial thalamus (AM)^19^ the medial entorhinal cortex (MEC),^23^ and the nucleus reuniens (Re),^117,118^ could mediate or gate the above functional dialogues. Collectively, our findings indicate that memory consolidation is accompanied by the shift of organizer and directionality of HPC-cortical dialogue.

Although cross-regional oscillatory communication involves in various brain function including cognitive tasks, very few studies interrogate their causal link to the mnemonic process, largely due to the technical limitation preventing simultaneous detection and intervention of the cross-area oscillatory coupling in a precise spatiotemporal manner. In the present study, we developed multiple intervention techniques, including random 4-12 Hz phase disruption and closed-loop intervention, to intervene in cross-area oscillatory dynamics in behaving animals. Using these techniques, we provide evidence to support the causal role of the HPC-cortical dialogues in memory consolidation and stabilization of remote fear memory. We propose that recent and remote memories are successively stored in the form of HPC-cortical and inter-cortical coherences (**Figure S18B**).

As oscillations and PAC are strongly dependent on behavioral and cognitive state,^36,37,48,49^ in the present study we simultaneously monitored EEG, EMG and autonomic signals including respiration and heart rates. These assessments help us to link the electrophysiological changes to their corresponding behavioral states, and thus determine whether the observed HPC-cortical coupling dynamics correlates with the animal’s locomotion state, sleep period or respiration rhythm. Another important network pattern is the ∼4Hz oscillation that dominates PFC activity during freezing.^36,83^ Since we here focus on the systems consolidation stage, during which animals remain in the home cage and freezing behavior is rarely expressed, this rhythm is not prominent in our observations. In addition, by comparing PFC– and HPC-based analyses and by examining state– and event-specific patterns of coupling, we were able to distinguish the encoding-stage PFC θ-associated PAS from the transient spindle-related low-frequency PFC events associated with consolidation-stage PAS.^119–123^ Thus, the phase-shifted coupling described here cannot be explained as a simple consequence of locomotion-related hippocampal theta, respiratory rhythm, or generalized cortical state changes.

Trace fear conditioning is a complex form of fear learning in which the CS and US are separated by a temporal gap, known as the trace interval. This form of learning requires the involvement of the HPC, PFC and other cortical regions. Previous studies have shown that both trace fear conditioning and contextual fear conditioning rely on hippocampal function. However, trace fear conditioning particularly depends on the coordinated interaction between the hippocampus and prefrontal cortex.^38,124^ As such, trace fear conditioning offers a well-defined experimental framework to investigate the role of hippocampal-cortical interactions in long-term memory consolidation.^20^

Together, our study provides a systematic characterization of the stepwise brain activity patterns over the process of system consolidation. The novel PAS phenomenon offers a unique signature of memory information transmission between HPC and PFC. Dissecting the cellular and molecular machinery underlying this electrophysiological signature will have crucial implication for understanding the fundamental process of remote memory storage and neurocomputation.

### Limitations of the study

Several limitations should be considered when interpreting the present findings. First, although we identified a robust PAS during both memory encoding and systems consolidation and demonstrated its causal contribution to remote memory formation, the cellular and circuit mechanisms underlying PAS remain unresolved. The observed phase shift may arise from multiple processes, including changes in cross-regional excitability, synaptic reweighting, and state-dependent modulation of long-range communication, but the relative contribution of these mechanisms was not directly examined in the current study. Furthermore, while the recurrence of PAS during consolidation suggests a potential relationship to replay-related information processing, the neuronal representations carried by PAS remain unknown. In particular, whether PAS is associated with spike sequences analogous to hippocampal replay has yet to be determined. Lastly, addressing this question will require stable identification of individual neurons across extended time periods. however, long-term tracking of single units over weeks remains technically challenging. Consequently, we were unable to directly compare neuronal firing patterns between encoding and consolidation stages. In addition, alternative models propose that memory representations may undergo transformation over time rather than preserving identical neuronal activity patterns. Future studies combining long-term cellular-resolution recordings with targeted circuit manipulations will be necessary to define the mechanistic basis and representational content of PAS during systems consolidation and remote memory storage.

## RESOURCE AVAILABILITY

### Lead contact

Further information and requests for reagents and resources should be directed to W. Lu.

### Material availability

This study did not generate new, unique reagents.

### Data and code availability

● All data and analyses necessary to understand and assess the conclusions of the manuscript are presented in the main text and in the supplementary materials.
● All original code has been deposited at Zenodo at [https://zenodo.org/records/20789326] and is publicly available as of the date of publication.
● Any additional information required to reanalyze the data reported in this paper is available from the lead contact upon request.

## ACKNOWLEDGEMENTS

We thank Dr. Jian-Guang Ni, Dr. Wei-Qun Fang, Dr. Lu-Yang Wang, Dr. Peng Yuan and all members of the Lu lab for discussion and support.

This work was supported by Brain Science and Brain-like Intelligence Technology-National Science and Technology Major Project (2021ZD0203502 to W.L.), National Natural Science Foundation of China (T2394531 to W.L.), Ministry of Science and Technology of China (2021YFA1101302 to W.L.), National Natural Science Foundation of China (32200835 to S.W.), and China Postdoctoral Science Foundation (2021M700847 to S.W.).

## AUTHOR CONTRIBUTIONS

Investigation: T.S., S.W., J.Z., H.Y., R.C.

Supervision: W.L., Q.W.

Funding Acquisition: W.L., S.W.

Visualization: T.S., S.W.

Writing: W.L., T.S., J.Z.

## DECLARATION OF GENERATIVE AI AND AI-ASSISTED TECHNOLOGIES IN THE WRITING PROCESS

During the preparation of this work, the author(s) used DeepSeek and ChatGPT in order to check for grammar errors in the codes and manuscripts. After using this tool or service, the author(s) reviewed and edited the content as needed and take(s) full responsibility for the content of the publication.

## SUPPLEMENTAL INFORMATION

**Document S1. Figures S1–S18, Tables S1 and S2**

## STAR METHODS

● KEY RESOURCES TABLE
● EXPERIMENTAL MODEL AND STUDY PARTICIPANT DETAILS
  ○ Animals
● METHOD DETAILS
  ○ Viral injection
  ○ Multisite electrode arrays fabrication and implantation
  ○ Anatomical and histological Analysis
  ○ Trace fear conditioning and contextual recall
  ○ Behavioral analysis
  ○ Electrophysiological recording
  ○ Sleep recording and state classification
  ○ LF phase intervention
  ○ Closed-loop optogenetic intervention based on coherence
  ○ Closed-loop optogenetic intervention based on SWR
  ○ Pharmacological interventions
● QUANTIFICATION AND STATISTICAL ANALYSIS
  ○ Spike sorting
  ○ LFP power and phase analysis
  ○ Phase-amplitude coupling analysis
  ○ Preferred phase and phase shift
  ○ LFP-spike analysis
  ○ Hippocampal SWR detection
  ○ Coherence analysis
  ○ Across-session analyses over system consolidation
  ○ Information directional analysis
  ○ Data analysis and statistics

## EXPERIMENTAL MODEL AND STUDY PARTICIPANT DETAILS

### Animals

Wildtype male Sprague Dawley rats were purchased from Vital River Laboratories (Beijing, China). Animals were group-housed with ad libitum access to food and water, maintaining a 12-hour light-dark cycle. Experiments were conducted during the light phase. All animals were aged 8-15 weeks and weighed 300-400 g at the time of electrode implantation. All animal experiments were approved by the Institutional Animal Care and Use Committee of Fudan University (Approval No. 202110027S) and conducted in compliance with the AAALAC guideline.

## METHOD DETAILS

### Viral injection

All surgeries were performed under inhaled anesthesia with isoflurane (5% induction, 1.5-2% maintenance). A heating pad was used to maintain the animals’ body temperature at 37 °C.

The recombinant adeno-associated virus (rAAV, serotype 2/9) carrying halorhodopsin (rAAV-hSyn-eNpHR3.0-mCherry) was stereotaxically injected into the HPC (3.5 mm posterior to bregma, 2.5 mm lateral to midline, 2.5 mm ventral to brain surface) or PFC (3.0 mm anterior to bregma, 0.5 mm lateral to midline, 3.0 mm ventral to brain surface) of wild-type Sprague-Dawley rats (4-5 weeks old, 190-220 g). The viral titer was 2 × 10¹² vg/mL. Virus was delivered bilaterally through a glass micropipette using a pulled glass capillary connected to a pressure microinjector. Each site received 500 nL of viral suspension at an infusion rate of 50 nL/min, with the injection needle retained for 10 min post-infusion to prevent reflux. Following surgery, incisions were closed, and animals recovered in a 37°C chamber until fully ambulatory.

### Multisite electrode arrays fabrication and implantation

Platinum-iridium (90/10) microwires with Teflon insulation were used for both grid and tetrode recordings. For grid recordings, 0.0014” diameter wires were sectioned into 5 cm electrodes. Pairs of these microwires were precisely loaded into 100 μm inner diameter silicon capillaries. Under stereomicroscopic guidance (40× magnification), the electrode tips were positioned in the same orientation with a calibrated 250 μm inter-tip separation, verified using readouts of the vernier caliper. For tetrode recordings, 0.0005” diameter Teflon-insulated Pt/Ir wires (California Fine Wire, USA) were twisted together. Briefly, four wires were folded and twisted using a motorized winding system, heat-fused with hot air, and inserted into silicon capillaries (75 μm inner diameter) for implantation. Tetrodes were used in selected experiments to increase the yield of simultaneously recorded single units. To accommodate the varying neuroanatomical diversity of brain regions, we designed a modular system by strategically stacking these capillary units, enabling cost-efficient construction of customizable 3D recording matrices optimized for target neuroanatomy (detailed in Multisite Electrode Array Implantation section). To ensure concurrent fidelity of both LFP and single-unit spikes during recording, the impedance profiles for every microelectrode channel were acquired using a NanoZ impedance tester (White Matter LLC, USA). Only the channels with impedance below 800 kΩ were selected for subsequent analysis.

Custom-built electrodes^31^ were chronically implanted into four brain regions using coordinates from the Paxinos & Watson rat brain atlas (7th ed.), referenced to bregma: PFC (AP +3.0 mm, ML +0.5 mm, DV –3.0 mm), ACC (AP +2.0 mm, ML +0.5 mm, DV –2.0 mm), HPC (AP –3.5 mm, ML +2.5 mm, DV –2.5 mm), and PPC (AP –4.2 mm, ML +4.5 mm, DV –1.5 mm). A stainless-steel skull screw over the cerebellum served as reference/ground. Animals were allowed a minimum of 7 days to recover before subsequent experiments.

In accordance with neuroanatomy, we allocated varying numbers of channels to different brain regions: 20-channel arrays for PFC, 12-channel arrays for ACC, 16-channel arrays for both HPC and PPC. Through four areas recording in training stage, we focused our interest in HPC and PFC. To capture more electrophysiology signals in interested areas, we recorded two areas (PFC and HPC), allocating 32 channels each. For optrode implantation, two optrodes was implanted into PFC (10-channel arrays + 200 μm core, 0.37 NA fibers (473 nm, 10 mW/mm²)) or HPC (8-channel array + 200 μm core, 0.37 NA fibers (473 nm, 10 mW/mm²)) bilaterally following a protocol similar to that of electrode implantation. Number of channels in other areas was set consistent to that of four areas recording. Two 1.2-mm ground screws were implanted symmetrically over the cerebellum.

For experiments involving simultaneous EEG and EMG recordings, a single-channel EEG electrode was implanted into the skull above the temporal lobe, and a two-channel EMG electrode was implanted into the posterior neck muscles. These electrodes occupied channels originally allocated to the PPC region within the 16-channel array. As a result, the number of available PPC channels was reduced to 13 in experiments with concurrent EEG/EMG recordings. All other methodological details were identical to those described above.

Fur over the neck region was removed for animals undergoing collar-based physiological recordings.

### Anatomical and histological Analysis

After remote memory recall, histology of surgical area was performed to verify electrode track and viral transduction. Rats were deeply anesthetized. Cardiac perfusion was performed with physiological saline and 4% paraformaldehyde sequentially. Brains were post-fixed in 4% PFA for 24 h at 4°C, cryoprotected in 30% sucrose-PBS at 4°C for 3-6 days until sinking, then embedded in optimal cutting temperature compound (OCT) for coronal sectioning at –20°C. Successive coronal sections of 45 μm were cut on a cryostat (Leica CM1950).

For electrode track verification, contained electrode implantation site were mounted on slides with a mounting medium containing nucleic acid dye (DAPI, 4’,6-diamidino-2-phenylindole). Electrolytic lesions were identified using confocal microscope (Nikon A1). Only data from recordings that confirmed lesions in the PFC, ACC, PPC, and HPC were included in our analyses.

For viral transduction validation, sections contained injection site (PFC or HPC) were mounted on slides with a mounting medium containing nucleic acid dye (DAPI, 4’,6-diamidino-2-phenylindole). Images were acquired using confocal microscope (Nikon A1). The location and extent of the injections or infections were visually assessed. Only those infections precisely targeting the PFC and HPC were considered for behavioral and electrophysiological analyses.

### Trace fear conditioning and contextual recall

All experimental phases (baseline habituation, conditioning training, and contextual memory retrieval) were conducted in a dedicated behavioral suite under controlled illumination. The conditioning chamber (30 × 25 × 21 cm) featured acrylic plastic front/back panels and flanked by aluminum sidewalls. The modular grid floor consisted of 36 stainless steel rods (3.2 mm diameter, 7.9 mm inter-rod spacing) interfaced with a programmable aversive stimulus generator via shielded cabling. To prevent odor-mediated contextual associations, the chamber underwent systematic decontamination between subjects using 10% ethanol followed by forced-air drying. Behavioral quantification was achieved through a ceiling-mounted camera acquiring 3.75 Hz grayscale video. All stimulus delivery and behavioral monitoring functions were synchronized through OmniPlex-64 (Plexon, USA).

Typically, electrical stimulation during conditioning introduces shock artifacts, including transient interference during stimulation and consistent 50/60 Hz line noise. Because our analyses were restricted to data segments within the inter-trial intervals (ITIs), which do not overlap with the stimulation period, the stimulator was disconnected during ITIs to eliminate any influence of shock artifacts. Specifically, an electromagnetic relay was incorporated into the stimulator circuit to electrically isolate it from the recording system. Controlled by the OmniPlex-64 system (Plexon, USA), the relay closed only during the 2-s shock delivery periods and remained open otherwise, thereby preventing any potential interference from the stimulator during non-shock intervals. In addition, the OmniPlex-64 synchronously recorded the relay control signal, allowing precise identification and removal of any residual stimulator artifacts during subsequent data analysis.

Before training, animals were gently handled for 3 minutes daily over 5 consecutive days and exposed to the training environment 24 hours prior to training. On the training day, animals were placed in the training chamber and underwent the following procedure: After a 180-second habituation period, animals received five consecutive training trials at 40-second inter-trial intervals (ITIs). Each trial began with a 20-second baseline, followed by a 20-second sinusoidal tone (2kHz, 80 dB), an 18-second trace interval, and a 2-second, 1 mA foot shock (unconditioned stimulus, US). To ensure comparability, the recording period in the CS only group was matched to that of the CS-US group by segmenting and analyzing identical time epochs aligned to the conditioning timeline. In both the CS-US and CS only groups, pre-FC data were obtained during the 180-s habituation period. Freezing behavior and neural activity were simultaneously recorded.

Recent and remote contextual memories were assessed using a contextual recall test. Animals were placed in the original training chamber without tone presentation for 600 s (comparable to the duration of the training session) on day 7 (recent memory) and day 28 (remote memory). Freezing behavior was recorded throughout the entire session for subsequent analysis.

### Behavioral analysis

Freezing behavior, defined as the absence of non-respiratory movements, was analyzed using FreezeFrame (Actimetrics), a software for automated motion detection. The criterion for defining freezing was based on the distribution of the motion index. Histograms typically exhibited a bimodal distribution, with a sharp peak near zero corresponding to freezing and a broader peak between 50 and 100 corresponding to movement. The freezing threshold was determined individually for each animal as the distinct trough between the two peaks in the motion index histogram, and typically fell within the range of 20-30. In addition, a minimum bout duration of 1 s was required for a motionless period to be classified as freezing, and this criterion was applied consistently across all experiments.

### Electrophysiological recording

The electrophysiological signals were amplified, subjected to band-pass filtering (1-1000 Hz for LFPs, 500-8000 Hz for spikes), and digitized using the Plexon Multichannel Acquisition Processor system. LFPs were digitized at a sampling rate of 1000 Hz, while spikes were digitized at 40 kHz and isolated using time-amplitude window discrimination and template matching.

Animals were habituated to the recording environment for three consecutive days before formal recordings. From 0 to 28 days after fear conditioning, electrophysiological recordings were performed consistently during the light phase of the circadian cycle each day. Animals were recorded in their home cages and allowed to move freely, during which they were predominantly resting or sleeping. To minimize electromagnetic interference, the home cages were placed inside a shielded chamber in which light intensity, temperature, and humidity were maintained at constant levels. Each recording session lasted 3-5 h to ensure the acquisition of sufficient sleep episodes.

Peripheral oxygen saturation (SpO_2_), heart rate, and respiratory rate were simultaneously monitored using a noninvasive pulse oximetry system (MouseOx, STARR Life Sciences). A collar-mounted sensor was then positioned around the neck, and physiological signals were acquired at a sampling rate of 15 Hz for further analysis. Rats were habituated to the collar-mounted sensor for 7 consecutive days prior to recording.

### Sleep recording and state classification

The protocols for sleep recording were detailed in a previous study.^119^ EEG recordings were obtained from a screw placed on the surface of the left cortex, positioned at AP –3.5 mm and ML 3 mm. Two EMG electrodes were inserted into the neck muscles. Insulated leads from the EEG and EMG electrodes were connected to a pin header, which was affixed to the skull using dental cement. All measures were taken to reduce rats’ discomfort during and after the surgical procedure. EEG/EMG recordings were performed for 2 h during either the light or dark phase, depending on the experimental design. Signals were acquired and analyzed using AccuSleep (https://github.com/zekebarger/AccuSleep_X), a compact neural network-based sleep scoring platform. Raw EEG and EMG signals were band-pass filtered (EEG: 0.5 – 100 Hz; EMG: 50 – 250 Hz), downsampled to 1000 Hz, and segmented into consecutive 5-s epochs using the integrated graphical user interface of AccuSleep.

Behavioral states were classified into wakefulness (Wake), non-rapid eye movement sleep (NREM), and rapid eye movement sleep (REM) based on standard EEG/EMG criteria. Wakefulness was characterized by low-amplitude, high-frequency EEG activity accompanied by high EMG tone. NREM sleep was identified by high-amplitude slow-wave EEG activity with reduced EMG activity, whereas REM sleep was characterized by low-amplitude, mixed-frequency EEG activity dominated by theta oscillations (6 – 9 Hz) and near-complete suppression of EMG tone. Sleep scoring was performed by trained experimenters blinded to experimental conditions.

### LF phase intervention

We utilized PlexDO (version 1.11.3, Plexon, USA) in conjunction with a MATLAB-based program to control the NI-DAQ square wave pulse generator for the delivery of optogenetic laser signals. To perturb the LF phase in the prefrontal cortex (PFC), we implemented a randomized LF frequency phase stimulation protocol. Photo-stimulation was administered at stochastic intervals specifically targeting the LF frequency range of 4-12 Hz. The inter-stimulation intervals were randomly distributed between 83 and 250 ms to introduce variability within the LF band.

### Closed-loop optogenetic intervention based on coherence

The experimental design is based on OmniPlex-64 and Master-9 (AMPI, USA). Custom MATLAB (version 2014b, MathWorks Inc, USA) scripts and MatlabClientDevelopKit (version 1.3.0, Plexon, USA) SDK are used to record and analyze real-time LFP signals. If trigger condition is detected, MATLAB outputs corresponding square wave pulses from NI-DAQ to laser through PlexDO (version 1.11.3, Plexon, USA) program and intermittently starts the laser according to the pulse signal. The laser is connected to the electrode by optical fiber.

To achieve closed-loop control, real-time LFPs should be recorded as described above and coherence needs to be calculated effectively among different brain areas during the experiment. A windowed Fourier Transform, with a window size set to 400 ms, was used on band passed LFPs to acquire real-time coherence for each pair of channels. Based on this value, closed-loop system determines the timing of optogenetic intervention.^109,110^

According to the experimental design, closed-loop control was configured to use a fixed pulse length mode. This method produces a light stimulus of a specific duration in the coherence-based closed-loop control. Coherence was calculated as mentioned previously. When the coherence from over 50% of channel pairs between two targeted brain areas significantly increased (exceeding 3× S.D. above the mean), the light stimulation persisted for a duration set to 400 ms for both γ coherence and LF coherence. The parameters, including the mean and standard deviation, were determined from a segment of data recorded before closed-loop stimulation and subsequently set in the software program.

For the fixed pulse length mode, delayed intervention experiments were established as the control groups. Delayed intervention means that when the trigger condition is detected, the system does not respond immediately but delays for a certain period before triggering. In this study, the delay time for the coherence trigger is set to 500 ms. Theoretically, the experimental results for both the delayed trigger should align closely with conventional response patterns. Since the delayed intervention method is employed in our experiments, we record the total stimulus duration of delayed trigger and conventional trigger to ensure they are the same proportion of stimulus duration under these controlled parameters.

To enhance the temporal precision of this intervention, coherence calculations for channels in different brain regions were conducted separately. The closed-loop system detected significant changes in functional connectivity within a brain region when coherence from more than 50% of channel pairs increased. Events characterized by coherence values exceeding 0.9 or oscillation amplitudes surpassing 2 mV were classified as aberrant, leading to the inhibition of optogenetic stimulation to avoid interference with head movements in vivo. To minimize software-induced delays, GPU parallel computing in MATLAB was utilized, facilitating multi-input multi-output closed-loop control requirements. In our closed-loop experiments, the optogenetic signal was activated with a delay of 20 to 40 ms from the timestamp that coherence from >50% channel pairs reached the threshold to the initiation of the stimulus, which was significantly shorter than the duration of increased coherence.

In order to evaluate the performance of real-time coherence detection, the precision, the recall, and the F1 score were calculated.

TP was defined as a significantly increased coherence event confirmed by offline analysis, while FP indicated that offline analysis identified incorrect coherence increase events. FN was used to describe instances where offline analysis failed to predict increased coherence events.

The precision (P) is defined as the ratio of true positives (TP) to the sum of true positives and false positives (TP + FP), indicating the proportion of positive predictions that were accurately identified.

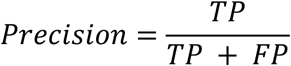

The recall (R) is calculated as the ratio of true positives (TP) to the sum of true positives and false negatives (TP + FN), reflecting the proportion of actual events that were correctly detected.

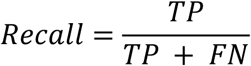

The F1 score, which is the harmonic mean of Precision and Recall, provides a balanced measure of the real-time Coherence detection performance, integrating both the accuracy of positive predictions and the completeness of event detection.

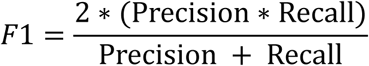

The in vivo experiment and real-time detection demonstrated the effectiveness of the proposed closed-loop control method in accurately detecting target events and responding promptly.

### Closed-loop optogenetic intervention based on SWR

We employed the same system as closed-loop control based on coherence to accomplish SWR-triggered closed-loop intervention, as previously described. Two optrodes were implanted bilaterally into HPC. A single HPC channel exhibiting the largest amplitude ripple was chosen for real-time processing of local field potentials (LFP) by custom MATLAB scripts. The root mean-square (RMS) values were calculated for the bandpass-filtered signal (within the 100-250 Hz range) using two sliding windows: a longer window spanning 2 seconds (denoted as RMS1) and a shorter window of 8 ms (denoted as RMS2). Ripples were defined as events with RMS2 exceeding 3×RMS1 for at least 8 ms. Once SWR was detected, one 60-ms square pulse was applied to HPC.

In order to evaluate the performance of real-time SWR detection, the F1 score was also calculated. Accordingly, predictions were categorized into four classes:

True Positive (TP): identified when the prediction indicated the presence of an SWR event, and the offline SWR detection confirmed this occurrence;

False Positive (FP): occurred when the model predicted an SWR event in a time window where no such event was present according to the offline SWR detection;

False Negative (FN): defined as instances where the prediction failed to indicate an SWR event that was actually present within the window;

True Negative (TN): characterized by the absence of an SWR event both in the prediction and the offline SWR detection.

The subsequent procedures for F1 score calculation adhered to the protocols detailed in the preceding methods section.

### Pharmacological interventions

Rats were implanted with guide cannulae targeting the dorsal CA1 region of the hippocampus (HPC) or the prefrontal cortex (PFC). Bilateral inactivation was achieved using CNQX (5 mM in physiological saline) due to its time-limited effects. To minimize the potential residual acute effects of CNQX, we deliberately performed the recordings prior to each CNQX application. Given the 24-hour interval between CNQX treatments, any remaining acute effects were largely diminished by the time of subsequent recordings. Specifically, CNQX was used to bilaterally inactivate the HPC from 3 to 7 days after fear conditioning in **Figures S12A and 12B**. Similarly, bilateral inactivation of the PFC was scheduled 14 to 21 days post-fear conditioning in **Figures S12C and 12D**.

## QUANTIFICATION AND STATISTICAL ANALYSIS

### Spike sorting

Spike sorting for single units was carried out using the Plexon Off-Line Spike Sorter (OFSS). Principal components 1 (PC1) and 2 (PC2) were computed for the original waveforms and plotted on a two-dimensional principal component space. Clusters of waveforms with similar characteristics were manually identified. Waveforms were considered to originate from a single neuron if they formed a distinct and isolated cluster in the principal component space and did not exhibit a refractory period shorter than 1 ms, as confirmed by autocorrelogram analysis (**Figures S6H**-**S6J**).

### LFP power and phase analysis

Spectral analysis of LFP was performed using the Chronux Package (K. Harris; http://chronux.org) with optimized parameters. We employed multi-taper spectrograms with time windows (256 samples, 1024 FFTs and 1.5 NW) for showing the changes in LFP. For specific characterization of θ oscillations, we employed time windows parameter of 2048 samples, 1024 FFTs and NW=1.5. The Morlet wavelet transform was employed to examine the dynamic changes in both power and phase within the frequency range of 1 to 150 Hz. This analysis enabled the observation of transitions between periods of slow and fast-γ oscillations, as depicted in Figure 1D. In a few recordings, a noise peak at 50 Hz was observed in the power spectrum. To remove this noise, a least squares regression was performed in the frequency domain using 800 ms windows, advanced by 200 ms, through the rmlinesmovingwinc function from the Chronux package. The 50 Hz noise was sporadic and did not exhibit any observable association with the behavior of the animals. The sporadic noise peak was thus proved to be irrelevant to the findings and conclusions of the paper. We removed trials in which any jump exceeded a z-score of 10 S.D. The cleaning process included visually inspecting every recording and manually removing any remaining artefactual trials. To determine the power envelope and phase of ongoing θ and γ oscillations, a bandpass filter was applied using a zero-phase-delay FIR filter with a Hamming window. The Hilbert transform of the bandpass-filtered signal was then computed as below (using the bzfilter function from https://github.com/buzsakilab/buzcode).

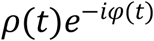

The length of the vector and the arctangent of the vector angle yield estimates for the instantaneous amplitude and instantaneous phase of the signal, respectively, at each time point. To normalize the data across different electrodes, sites, and animals, power values were calculated as the fold change from the pre-FC (30-second before FC) or day –1 (30 min recording from the day before FC) as baseline.

### Phase-amplitude coupling analysis

We evaluated the degree of cross frequency coupling across a wide range of frequency combinations in all recorded channels based on the modulation index (MI)^51^:

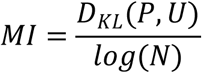

The instantaneous phase of the lower-frequency signal and the amplitude of the higher-frequency signal were extracted using the Hilbert transform. Additionally, each trial was trimmed to the final time window of interest, which was set as 35 s relative to the onset of the ITI period. This step ensured the removal of filter artifacts that could occur at the signal edges. The analysis window length remained consistent in all trials. Subsequently, the phase and amplitude signals from all trials were concatenated, and the MI was computed using 50 phase bins. The MIs were calculated separately for before FC and ITIs. To evaluate the statistical significance of cross-frequency coupling, we generated shuffled datasets by temporally shifting the γ band amplitude relative to the 4-12 Hz phase. Specifically, for each iteration, the γ power time series was shifted by an integer value between 1 and 30 seconds, effectively disrupting the temporal alignment between phase and amplitude while preserving the individual signal characteristics. This procedure was repeated for all shift values to generate a null distribution of MI. The observed MI from the original, unshifted data was then compared against this null distribution to determine whether the phase-amplitude coupling was statistically significant. All subsampled trials were included in an unbiased manner to identify significant channels displaying phase-amplitude coupling (PAC).

### Preferred phase and phase shift

The preferred phase of phase-amplitude coupling was calculated as the mean γ coupled θ phase by the MATLAB function ‘circ_mean’ from the Chronux open-source MATLAB toolbox (http://chronux.org/). Considering that the number of training blocks is linearly distributed, while the PAC preferred phase is circular data, we utilized circular-linear regression to yield a reliable estimation of the slope, fit goodness of the regression line, along with a correlation coefficient (R^2^ value and p value) suitable for phase shift.^73^ We used the linear regression method to confirm the correlation between task performance (mean freezing % of Trial 4 and Trial 5) and occurrence of phase shift (slope and R^2^ value).

### LFP-spike analysis

Phase-locking analysis: The LFP signal underwent digital band-pass filtering (4-12 Hz for θ, 60-90 Hz for γ) using a zero-phase-delay filter with an order of 5 and sample frequency operation. The phase component was determined through a Hilbert transform, enabling the assignment of a corresponding phase to each spike. Rayleigh test was used to calculate the significance of the phase-locking relationship, with significance determined at *P* < 0.05, unless specified otherwise. However, to ensure a reliable assessment of spike phase, a minimum threshold was set for all analyses. Specifically, during comparisons of phase-locking throughout the consolidation stage, only units with a minimum of 100 spikes fired on each day were included in the analysis. By employing a shuffled predictor technique, we confirmed that any potential asymmetries in the θ or γ oscillation did not generate spurious coupling. Consequently, a significant phase-locking was observed at a threshold of 5% by chance.^51^

θ-spike phase shift: The spike phase was obtained as previously described in section: LFP-spike phase-locking analysis. The preferred phase was calculated as circular mean value of spike phases by the MATLAB function ‘circ_mean’ from the Chronux open-source MATLAB toolbox (http://chronux.org/). For the purpose of plotting, the spike histograms were organized into 50 phase bins. Phase shift analysis is consistent with those in the previous method section: Preferred phase and θ-γ Phase shift.

Spike distribution in γ power: The LFP signal underwent digital band-pass filtering (60-90 Hz for γ) using a zero-phase-delay filter. The amplitude or power component of HPC fast-γ band-pass LFP was determined through a Hilbert transform, enabling the assignment of a corresponding fast-γ power to each spike. HPC fast-γ power was categorized into four distinct classes based on their amplitude values. Subsequently, each spike was assigned to one of four bins corresponding to the category of its associated fast-γ power value.

γ trough-triggered histogram: To compute γ trough-triggered histograms of spike counts for single-unit activity, custom scripts were utilized. To summarize the process, spike histograms were organized into time bins of 1 ms. Cross-correlations were then determined for spikes occurring during periods of strong fast-γ power (>1.5 standard deviations (S.D.) above the mean) compared to the weak fast-γ power (<1.5 S.D. above the mean). Considering the variation in spike number, the histograms of spike counts were normalized by the standard deviation of values at lags of –30 to –50 and +30 to +50 ms.

### Hippocampal SWR detection

Buzcode scripts (https://github.com/buzsakilab/buzcode) was employed to detect SWR in the hippocampus. The LFP average was filtered within the ripple band (110-250 Hz) using two separate zero phase-shifted filtering processes. First, a high-pass Butterworth filter (eighth order, zero phase-shifted, with a cutoff frequency of 130 Hz) was applied, followed by a low-pass Butterworth filter (tenth order, zero phase-shifted, with a cutoff frequency of 250 Hz). A smoothed envelope was generated of this signal by calculating the magnitude of its Hilbert transform and convolving it with a Gaussian window. Subsequently, we established two thresholds for SWR detection based on the mean (μ) and standard deviation (σ) of the LFP in the SWR band; the upper and lower thresholds were defined as μ + 4 × σ and μ + 1 × σ, respectively. For an epoch to be considered an SWR, the SWR power had to exceed the upper threshold for at least one sample and the lower threshold for a minimum of 50 ms. Each epoch where the SWR power surpassed the lower threshold indicated the onset and conclusion of the SWR, with the duration of each SWR determined accordingly.

We investigated the influence of SWRs on the coherence between HPC-cortex and the inter-cortical coherence. The measurement used to examine this effect was consistent across all evaluations. The analysis was conducted during two conditions: SWRs (SWR+) and random events (SWR-). In the SWR+ condition, we focused on the 1-second time intervals following the termination of SWRs. On the other hand, the SWR– condition involved selecting random events and examining the subsequent 1-second time intervals. The total number of events in the SWR-condition matched the number of SWRs in the SWR+ condition. Additionally, we investigated γ coherence, θ coherence, phase amplitude coupling, and the firing rate of individual units during SWRs.

### Coherence analysis

Coherence is often used in neuroscience to quantify the functional connections between two brain regions. To calculate coherence between two LFP signals x(t) and y(t), multi-taper method from the Chronux open-source MATLAB toolbox (http://chronux.org/) are used to get the power spectrum of the signal. For every taper, the result of the Fourier transforms of x(t), (t = 1,2, …, N) is

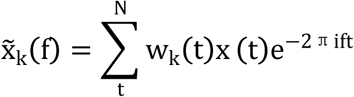

The definition of the cross-power spectrum between two time series x(t) and y(t) is

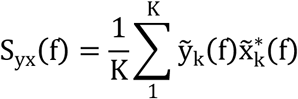

Coherence was calculated as follows:

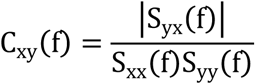

Where, S_xx_(f) and S_yy_(f) mean the self-power spectral densities of x(t) and y(t). 0 ≤ C_xy_(f) ≤ 1.

### Across-session analyses over system consolidation

In order to enhance the robustness of two-stage prediction across diverse animal data sets, the metric was individually normalized for each animal. This normalization procedure standardized the metric to a range between one and zero, representing the maximum and minimum values of the original metric, respectively. Subsequently, the convolved and normalized data were fitted to a Sigmoid function characterized by an ‘S’-shaped curve. This curve facilitated the prediction of two-stage transitions over time, distinguishing between high and low values.^22^ We utilized a standard form of the Sigmoid function, with ‘U’ and ‘L’ denoting the upper and lower boundaries, ‘xmid’ representing the midpoint parameter (symmetric point of the ‘S’-shaped curve), and ‘k’ indicating the slope parameter. The xmid parameter of the Sigmoid function played a crucial role in determining the temporal transition between the two stages. Specifically, the fittings of the Sigmoid function were focused on days 2-19 for γ coherence and the slope and R^2^ value of phase shift in Figures 4F**-4H and S10F-S10H**, as well as the slope and R^2^ value of phase shift and the average of LF coherence in Figures 4Q**-4S and S11H-S11J**. In contrast to the PAS analysis, where the slope was derived from phase data across all days, sigmoidal fitting used a five-day sliding window to estimate daily slope and R^2^ values, centered on each target day, i.e., the target day together with the two preceding and two following days. To evaluate the robustness of the sigmoid function, we performed a cross-validation procedure. In each iteration, a random subset of the original data points (10-20%) was excluded, and the remaining data were used to refit the sigmoid function. The transition point, defined as the time at which the sigmoid reached 50% of its maximal change, was extracted from each refit. The variability of the transition point across iterations was quantified using the coefficient of variation and expressed relative to the estimate obtained from the original fit. Transition points with variability below 10% were considered highly stable, while those with variability below 15% were classified as showing good stability (**Figures S10I, S10J, S11K and S11L**). These analyses were instrumental in identifying the most significant changes in γ coherence and the slope and R^2^ value of phase shift, and in comparing the sequential relationships between the increment in the average LF coherence with the subsequent decline in the slope and R^2^ value of phase shift.

### Information directional analysis

LFP trough-triggered analyses: In the γ and LF trough-triggered analyses, the troughs of γ and LF oscillations were defined as peaks in the power envelope of the 60-90 Hz or 4-12 Hz bandpass signal exceeding 1.5 S.D. from the mean power. Similar outcomes were achieved using peak-triggered averages. The neuronal activity occurring around these troughs was subsequently averaged to generate peri-event time histograms. For coherence analyses after trough-triggered data was calculated as

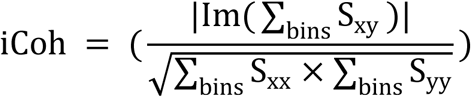

Here, S_xy_ represents the cross-spectrum, while S_xx_ and S_yy_ denote the auto-spectra. The summation is performed over the spectrogram bins that correspond to the quantified state. By maintaining the imaginary component of the normalized cross-spectrum, the coherence value is inversely weighted in relation to the time lag between the two signals. As a result, it is only responsive to time-lagged signals, while the influence of completely synchronous signals is excluded. Considering the highly synchronous characteristic of the oscillation being studied and the minimal phase lag, imaginary coherence is likely to underestimate the interaction strength.

Hubs’ centrality analysis: Hub centrality measures how important a node is by how strongly it connects to other important nodes.^100^ Hub and authority centrality analyses were performed using the MATLAB function ‘centrality’. Hub and authority scores constitute a pair of recursively defined centrality metrics for directed networks. In this framework, the hub score of a node (brain region) reflects the weighted sum of the authority scores of nodes it projects to, whereas the authority score reflects the weighted sum of the hub scores of nodes projecting to it. Directed weighted graphs were constructed, with edge weights specified by γ-band coherence for the recent stage and LF-band coherence for the remote stage. Edge weights were incorporated by specifying the ‘Importance’ parameter in the ‘centrality’ function, such that weighted sums were used instead of unweighted successor or predecessor counts. Centrality scores were rescaled within each connected component so that the sum of scores equaled 1. The same procedure was applied to 500 null networks generated by randomizing network connectivity. Node-wise centrality scores from the empirical networks were compared against the null distributions to assess statistical significance.

Spike-spike correlation: To compute spike-spike cross-correlations for simultaneously recorded multi-unit activities from two specific brain region, cross-correlograms for the spike trains were utilized. To summarize the process, pairwise spike time differences were calculated for all spikes from two specific brain region and organized into time bins of 20 ms and a time window between – 200 ms and +200 ms. In the recent stage, we use cross-correlograms with the spikes of the HPC cell serving as reference. In the remote stage, we use cross-correlograms with the spikes of the PFC cell serving as reference. If the mean z-score of –200 ms to 0 ms was negative and the mean z-score of 0 ms to +200 ms was positive, the referenced cell was determined to fire preferentially before the other cell. Conversely, the preferred firing order was in the opposite direction.

To establish the statistical significance of the lead/lag relationship, we performed a spike jitter analysis. For each spike train in the reference region, spike times were randomly jittered within a ±5 ms window, thereby disrupting precise spike timing while preserving overall firing rates. This procedure was repeated 100 times to generate a null distribution of the LFP-spike coupling metric. The coupling value obtained from the original, unshuffled data was then compared with this null distribution to determine whether the observed temporal coupling exceeded chance levels.

Spike-LFP modulation analysis: To investigate the synchronization of spikes with the oscillatory phase of LFPs, pairwise phase consistency was employed as the method of choice. Unlike other commonly used metrics for phase-locking, pairwise phase consistency is not influenced by the number of spikes, offering an unbiased evaluation.

The same spike jitter analysis was applied as described above for spike-spike correlations.

LFP-LFP granger causality analysis: Granger causality analysis was conducted utilizing the arfit toolbox in Matlab. The order for each pair of PFC-HPC LFP signals was established based on Schwarz’s Bayesian Criterion, ranging from 2 to 120 samples (equivalent to 1-60 ms). The magnitude of the Granger causality from A to B lead was then determined as GCI^A→B^ / (GCI^A→B^ + GCI^B→A^) for each animal.

### Data analysis and statistics

Data analysis was conducted using specialized routines developed in MATLAB (MathWorks, Inc., Natick, MA, USA). All effects reported as statistically significant exceeded a significance threshold of 0.05. All independence tests were conducted in a two-tailed manner. Paired t-tests were employed to analyze paired values reflecting changes across different situations. ANOVA tests involving more than two groups were followed by post-hoc tests, which used Bonferroni correction for multiple comparisons unless otherwise explicitly stated. All reported values are expressed as mean ± s.e.m. unless otherwise noted.

