## Supplemental Data 1 for "Hippocampal-cortical coupling dynamics drive system consolidation of remote memory"

**SUPPLEMENTAL INFORMATION**

**SUPPLEMENTAL FIGURES AND FIGURE LEGENDS**

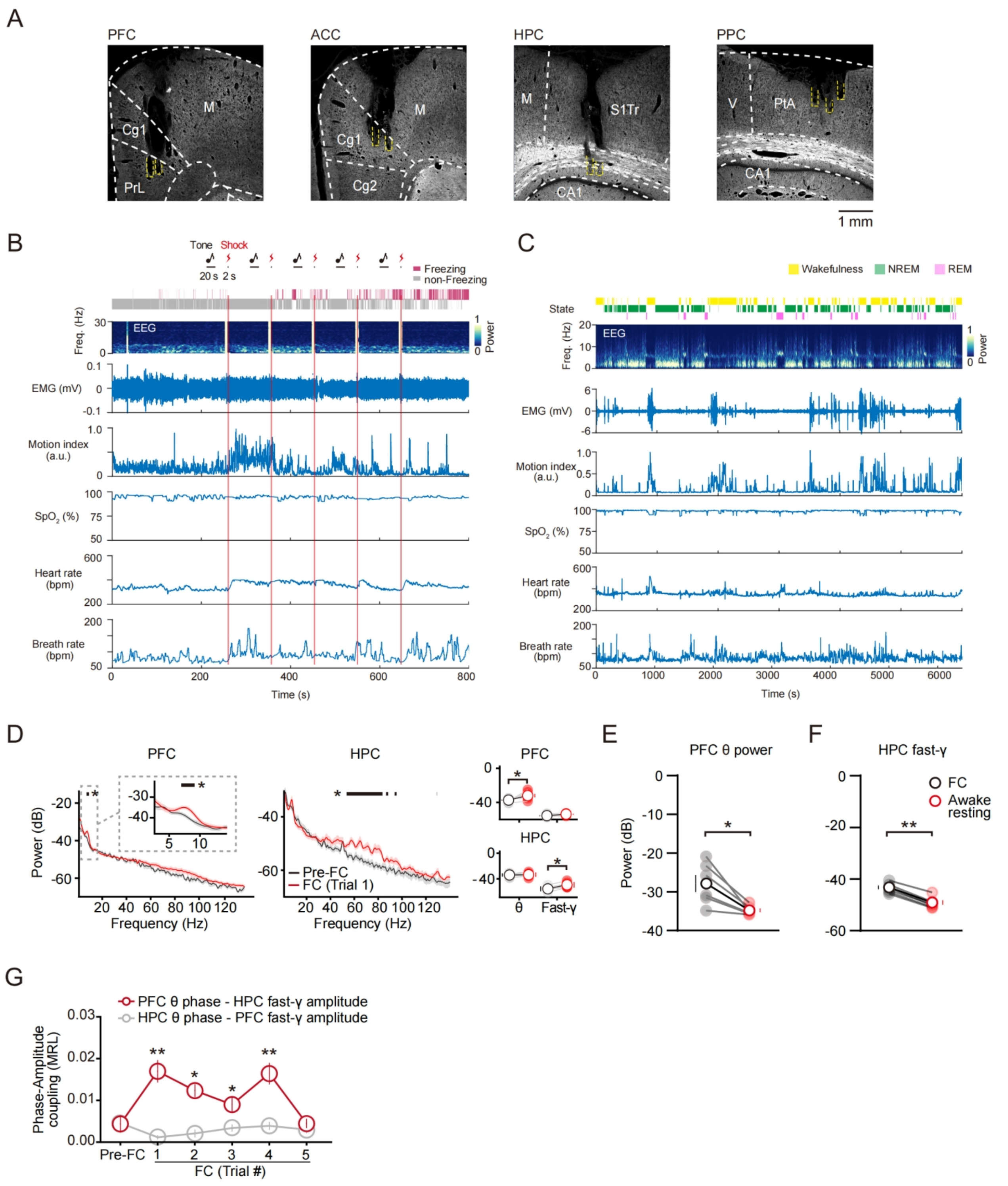

**Figure S1. [Electrode localization and multimodal recordings], related to Figure 1.**

(A) Histological examination of anatomical tracks confirmed the positions of recording electrodes after the recording experiments.

(B) Representative sample illustrating simultaneous multimodal monitoring during fear conditioning. Magenta and gray bars indicate freezing and non-freezing states, respectively. Time-resolved power spectra and waveforms of EEG and EMG, together with motion index, blood oxygen saturation, heart rate, and respiration, exhibit distinct state-dependent amplitudes profiles during FC training.

(C) Representative sample illustrating simultaneous multimodal monitoring in the home cage during the consolidation stage. Yellow, green, and pink bars indicate wakefulness, NREM and REM states, respectively. Time-resolved power spectra and waveforms of EEG and EMG, together with motion index, blood oxygen saturation, heart rate, and respiration, exhibit distinct state-dependent amplitudes profiles.

(D) LFP power spectra revealed power increased in PFC θ (4-12 Hz; left) and HPC fast-γ (middle). Black lines above the spectra indicate $*P < 0.05$, multiple t-tests. Right insets show zoom-in θ (4-12 Hz) power change. Right column shows and statistics of average power changes. n = 6, paired t test, $*P < 0.05$.

(E) The PFC θ power in fear conditioning was increasing compared to awake resting in home cage after conditioning. (n = 6, PFC θ FC: -27.88 ± 2.20; Awake resting: -34.80 ± 0.49, paired t-test, $*P < 0.05$, center values denote mean ± s.e.m.).

(F) The HPC fast-γ power in fear conditioning was increasing compared to awake resting in home cage after conditioning. (n = 6, HPC fast-γ FC: -43.53 ± 0.78; Awake resting: -49.37 ± 0.90, paired t-test, $**P < 0.01$, center values denote mean ± s.e.m.).

(G) Evolution of cross-frequency PAC strength in the HPC–PFC circuit. PFC θ and HPC fast γ exhibited prominent coupling during conditioning trials, whereas no such coupling was observed between HPC θ and PFC fast γ (n = 6 rats; Bonferroni-corrected Wilcoxon signed-rank test; $*P < 0.05$, $**P < 0.01$). MRL, mean resultant length.

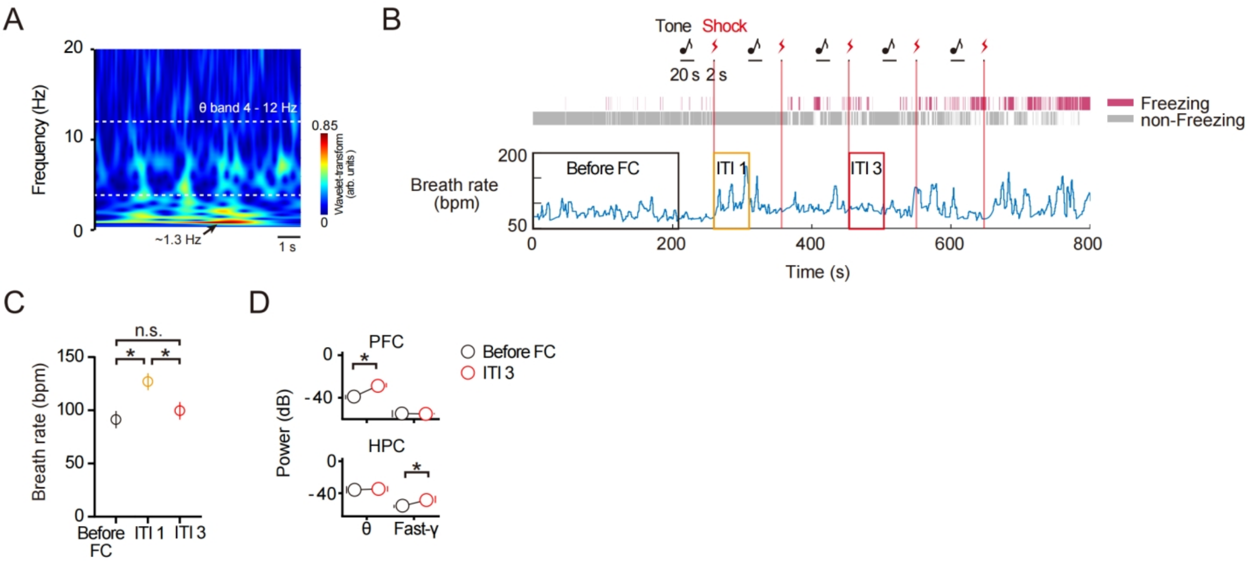

**Figure S2. [Respiratory dynamics did not account for PFC θ oscillations], related to the main text.**

(A) The characteristic respiratory rhythm during conditioning, characterized by a power spectral peak at ~1.3 Hz.

(B) Representative respiratory activity during fear conditioning, illustrating the epochs analyzed: before FC, ITI1, and ITI3. Magenta and gray bars above represent Freezing and non-freezing, respectively.

(C) Quantitative analysis of breath rate across epochs. The rate was significantly elevated in ITI1 compared to before FC and ITI3, while ITI3 returned to baseline levels. (n = 6, Before FC: 91.25 ± 8.01; ITI1: 127.00 ± 8.04; ITI3: 99.66 ± 8.13, unpaired t-test, *$P < 0.05$).

(D) Statistics of average power changes shows the increasing of PFC θ and HPC fast-γ power in ITI3 compared to the before FC baseline. (n = 6, paired t-test, *$P < 0.05$)

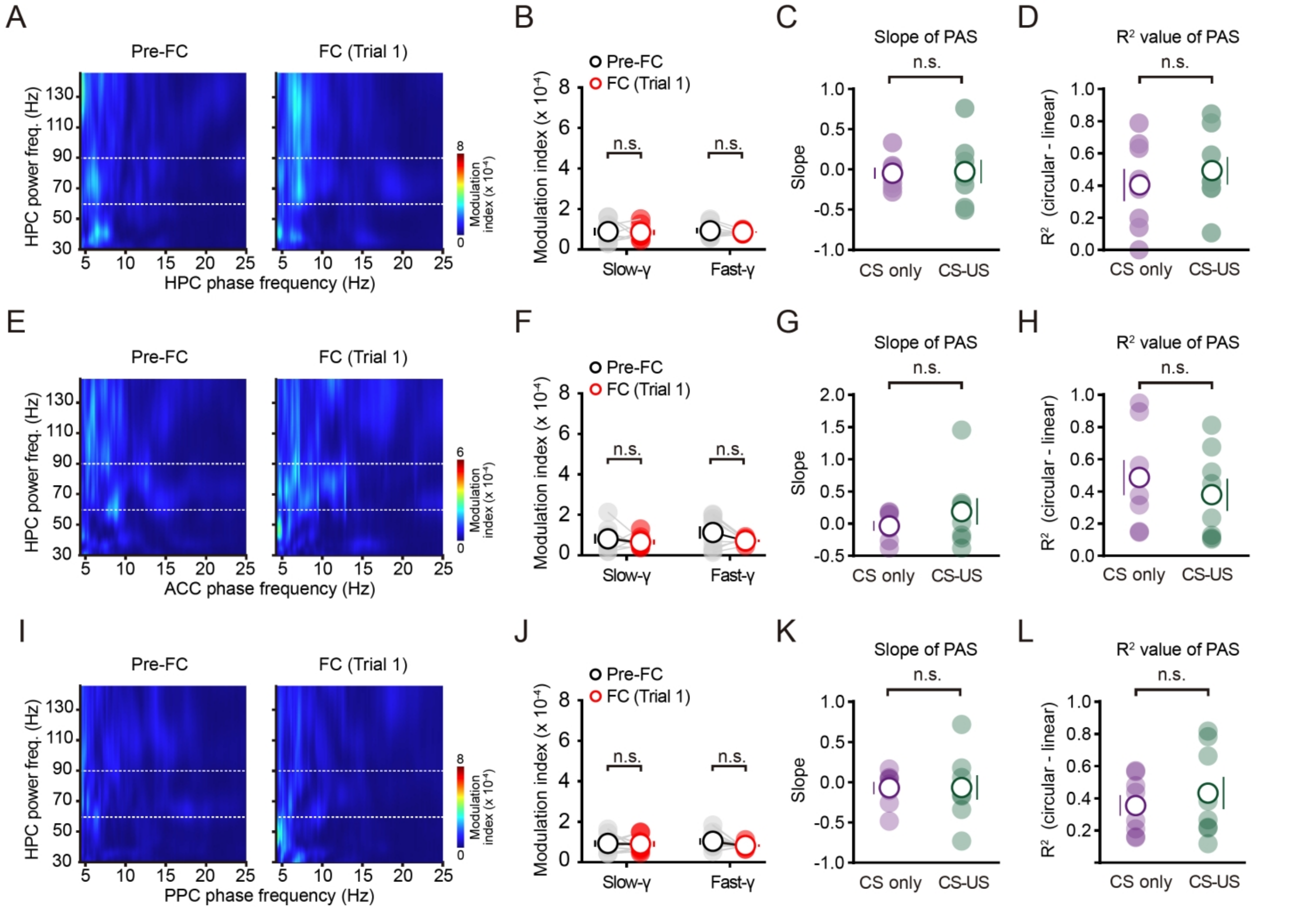

**Figure S3. [Absence of PAC and PAS in HPC-ACC, HPC-PPC, and HPC-HPC pairs], related to Figure 1.**

(A) Phase-amplitude comodulogram of representative LFPs in HPC failed to demonstrate prominent modulation of HPC fast-γ power (regions between the dotted line; y axis) by HPC θ oscillation phase (x axis) at trial 1 compared to pre-FC.

(B) Statistical analysis of modulation index reveals no stronger modulation of HPC fast-γ (60-90 Hz) by HPC θ at trial 1 (red) compared to pre-FC (gray). (n = 8, paired t-test, n.s., no significant, mean ± s.e.m.)

(C and D) Quantification of PAS with slope of PAS or mean $R^2$ value reveals no PAS of within-HPC θ-γ. Slope equaling zero indicates no significant PAS. (n = 8, unpaired t-test, n.s., no significant, mean ± s.e.m.)

(E) Phase-amplitude comodulogram of representative LFPs in HPC and ACC failed to demonstrate prominent modulation of HPC fast-γ power (regions between the dotted line; y axis) by ACC θ oscillation phase (x axis) at trial 1 compared to pre-FC (left; 30 s before FC). Warmer colors indicate stronger modulation.

(F) Statistical analysis of modulation index reveals no modulation of HPC fast-γ (60-90 Hz) by ACC θ at trial 1 (red) compared to pre-FC (gray). (n = 8, paired t-test, n.s., no significant, mean ± s.e.m.)

(G and H) Quantification of PAS with slope of PAS or mean $R^2$ value reveals no PAS in HPC-ACC circuits. Slope equaling zero indicates no significant PAS. (n = 8, unpaired t-test, n.s., no significant, mean ± s.e.m.)

(I) Phase-amplitude comodulogram of representative LFPs in HPC and PPC failed to demonstrate prominent modulation of HPC fast-γ power (regions between the dotted line; y axis) by PPC θ oscillation phase (x axis) at trial 1 compared to pre-FC.
(J) Statistical analysis of modulation index reveals no stronger modulation of HPC fast-γ (60-90 Hz) by PPC θ at trial 1 (red) compared to pre-FC (gray). (n = 8, paired t-test, n.s., no significant, mean ± s.e.m.)
(K and L) Quantification of PAS with slope of PAS or mean $R^2$ value reveals no PAS in HPC-PPC circuits. Slope equaling zero indicates no significant PAS. (n = 8, unpaired t-test, n.s., no significant, mean ± s.e.m.)

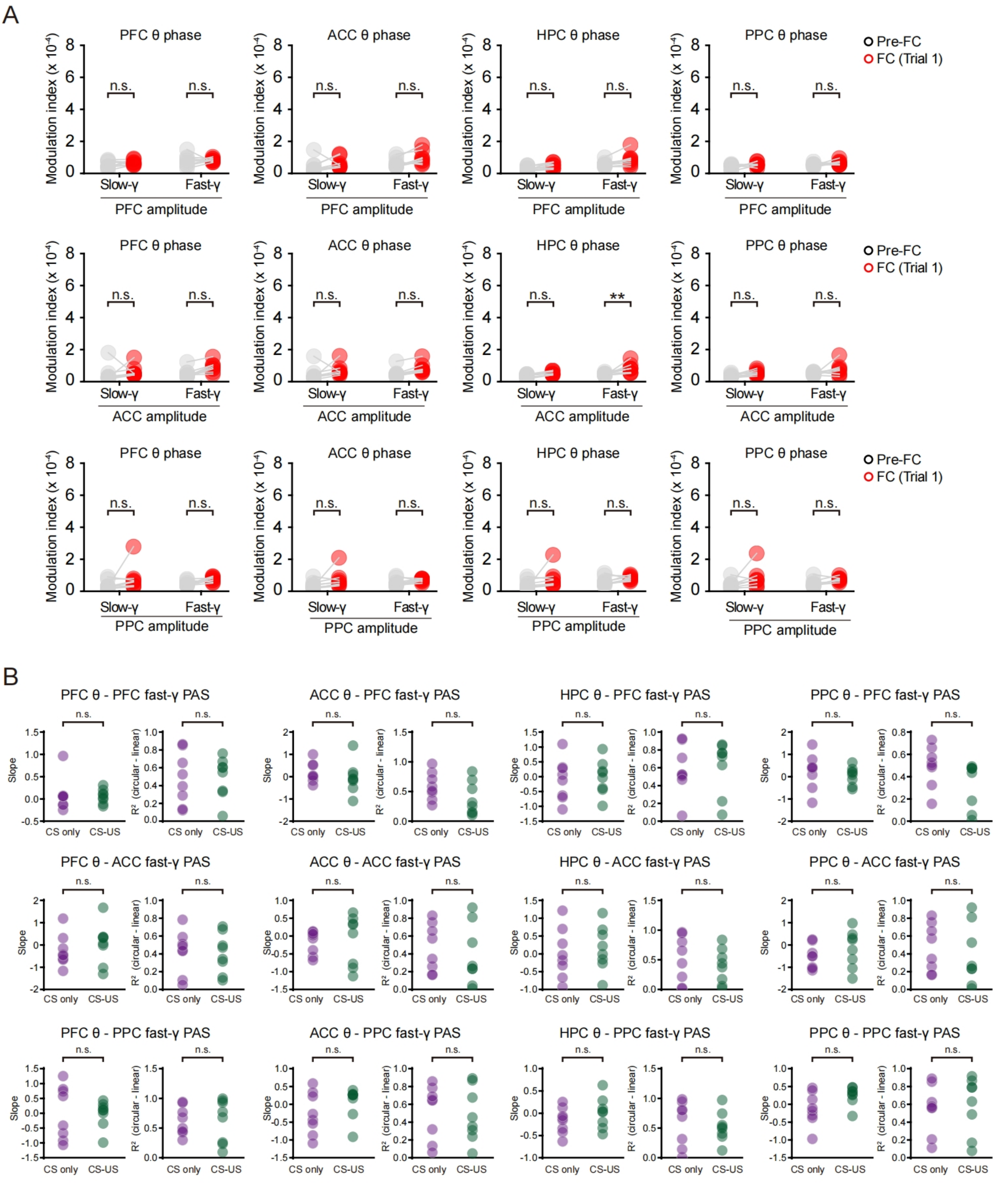

**Figure S4. [Quantification of θ-γ PAC and PAS in other within- or inter-regional combinations during conditioning], related to Figure 1.**

(A) The modulation index (MI) matrix quantifying coupling between γ (slow or fast) and θ oscillations showed no stronger modulation at trial 1 (red) compared with pre-FC (gray), with the exception of HPC-ACC θ-fast-γ coupling. (n = 8, paired t-test, $^{**}P < 0.01$, n.s., no significant, mean ± s.e.m.)

(B) Quantification of PAS using either the slope or the mean $R^2$ value, revealed no significant PAS between θ oscillations in HPC, PFC, ACC, and PPC and fast-γ activity (60-90 Hz) in PFC, ACC, and PPC. Slope equaling zero indicates no significant PAS. ($n = 8$, unpaired t-test, n.s., no significant, mean ± s.e.m.)

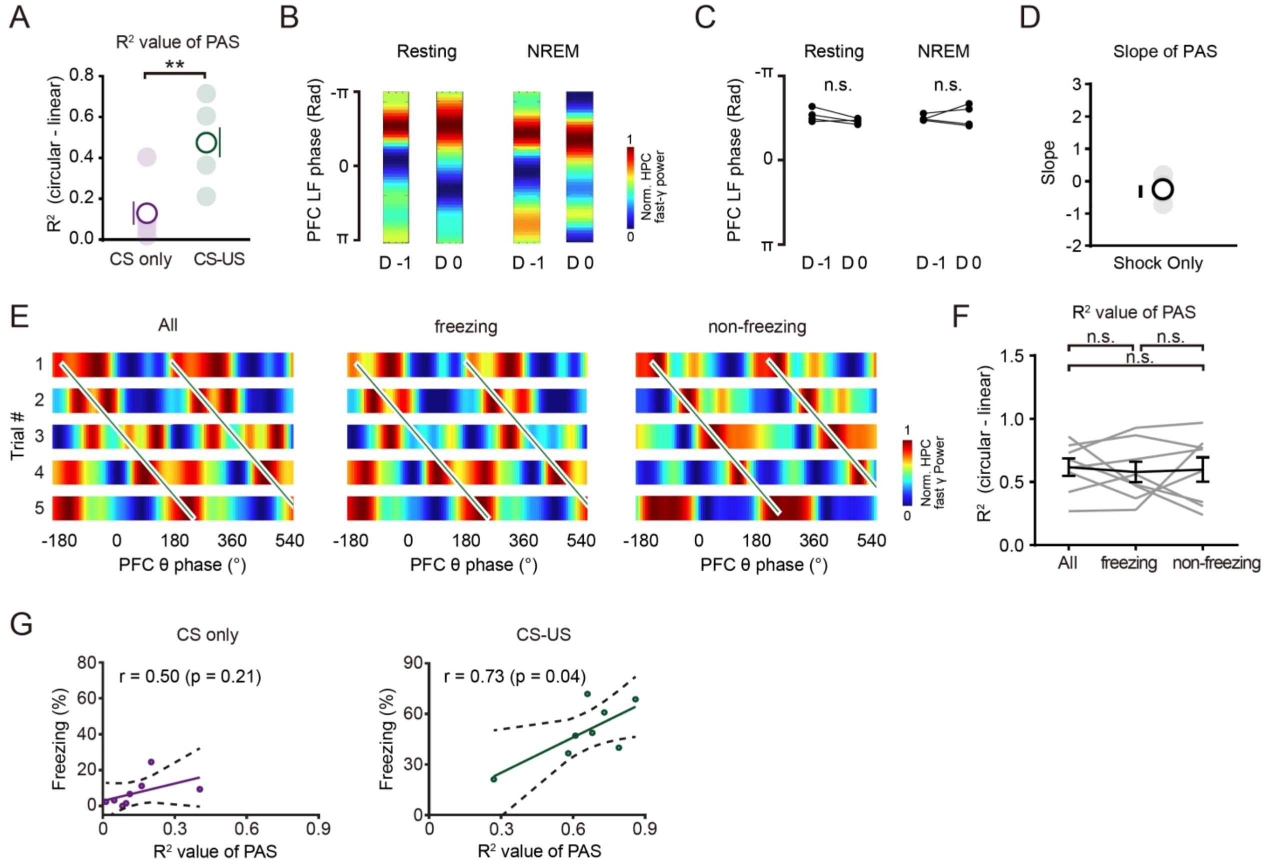

**Figure S5. [Coupling phase resetting and correlation of PAS with learning performance], related to Figure 1.**

(A) Quantification of PAS. Mean Pearson's correlation coefficient ($R^2$) was used to assess the goodness-of-fit in phase-trial regression. CS-US group elicited higher $R^2$ value than CS only group (CS only: 0.13 ± 0.06; CS-US: 0.48 ± 0.07, unpaired t test, $^{**}P < 0.01$).

(B) Representative phase distributions illustrating that the preferred coupling phase is consistent before and after training, during both resting state and non-REM (NREM) sleep.

(C) Statistical analysis quantifying the preferred phase. Data confirm no significant change following training in either resting or NREM states. paired t-test, n.s., no significant, mean ± s.e.m.

(D) Quantification of PAS with slope of PAS in HPC-PFC circuits of shock only group. Slope equaling zero indicates no significant. (Shock only: PAS: -0.25 ± 0.19).

(E) Comodulogram showing trial-to-trial phase shift of HPC fast-γ preferred PFC θ phase during the entire period (left), freezing (middle) and non-freezing (right) periods. Green lines show results of linear regression fit.

(F) Quantification of PAS with mean $R^2$ value. No difference was observed between freezing and non-freezing period. Repeated measures (RM) one-way ANOVA with Tukey post-hoc test, $F(7,14) = 2.19$, n.s., no significant, mean ± s.e.m.

(G) Learning performance was positively correlated with PAS, as indicated by Pearson correlation coefficient of learning performance and the extent of PAS that was defined as $R^2$ value of PAS (CS only: $r = 0.50$, $P = 0.21$; CS-US: $r = 0.73$, $P = 0.04$). Each cycle represents data from an individual animal.

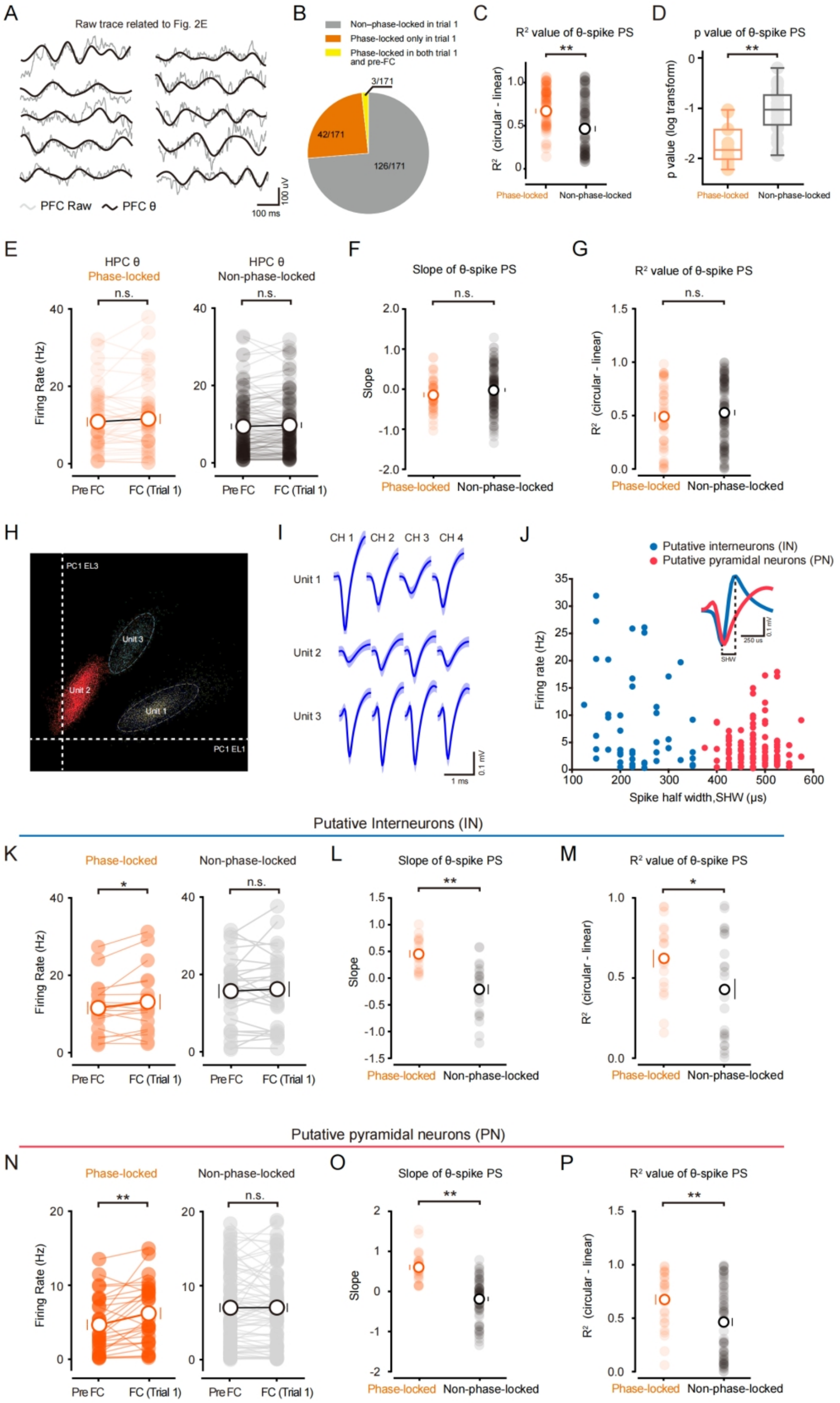

**Figure S6. [HPC units phase-locking to PFC θ demonstrated significant θ-spike PS in both INs and PNs], related to Figure 2.**

(A) Representative raw and θ band-pass-filtered trace sample related to **Figure 2E**.

(B) Pie chart shows the distribution of 171 neurons across three categories: non-phase-locked in trial 1 (n = 126, 73.7%), phase-locked only in trial 1 (n = 42, 24.6%), and phase-locked in both trial 1 and pre-FC (n = 3, 1.8%).

(C and D) Statistical analysis of $R^2$ (C) and p value (D) of PAS in phase-locked and non-phase-locked HPC units. $R^2$, Phase-locked HPC unit: 0.68 ± 0.05; non-phase-locked: 0.45 ±

0.04, unpaired t test, $^{**}P < 0.01$. D, mean log-transformed p value of the correlations shown in **Figure 2G**, unpaired t test, $^{**}P < 0.01$.
(E) The HPC θ phase-locked units in the HPC did not show a significant change in firing rate at trial 1. Similarly, no significant alteration was observed in other non-phase-locked units (unpaired t-test, phase-locked HPC unit: n = 45, $P = 0.21$; non-phase-locked: n = 126, $P = 0.13$).
(F and G) HPC units that phase-locked to HPC θ did not demonstrate a significantly different θ-spike PS compared to non-phase-locked units, as indicated by similar slopes (Phase-locked HPC unit: -0.14 ± 0.06; non-phase-locked: -0.016 ± 0.047, unpaired t-test, $P = 0.16$). and $R^2$ value (Phase-locked HPC unit: 0.49 ± 0.04; non-phase-locked: 0.53 ± 0.03, unpaired t-test, $P = 0.44$). n.s., no significant, mean ± s.e.m.
(H) Individual HPC units clustered from tetrode recordings.
(I) Mean waveforms of extracellular potentials from example units in (H).
(J) Classification of pyramidal cells and interneurons. Among the recorded neurons, 99 out of 145 units were identified as putative projection neurons (PNs, marked with red circles), while 46 out of 145 units were categorized as putative interneurons (INs, marked with blue circles). This classification was achieved using an unbiased unsupervised clustering algorithm that considered two electrophysiological characteristics: firing rate and spike half width (SHW). The inset displays the average waveform of a typical PN and IN, illustrating the measurement of SHW.
(K) The PFC θ phase-locked HPC INs exhibited increased firing rate at trial 1. As a control, no such alteration was observed in other non-phase-locked HPC INs (Paired t-test, phase-locked HPC INs: n = 18, $^{*}P < 0.05$; non-phase-locked HPC INs: n = 28, $P = 0.58$).
(L) HPC INs that phase-locked to PFC θ demonstrated more prominent θ-spike PS, indicated by higher slope than non-phase-locked units (Phase-locked HPC INs: 0.45 ± 0.07; non-phase-locked HPC INs: -0.21 ± 0.09, unpaired t-test, $^{*}P < 0.05$).
(M) Quantification of θ-spike PS. Mean Pearson's correlation coefficient ($R^2$) was used to assess the goodness-of-fit in phase-trial regression. HPC INs that phase-locked to PFC θ elicited higher $R^2$ value than non-phase-locked HPC INs (Phase-locked HPC INs: 0.62 ± 0.06; non-phase-locked HPC INs: 0.43 ± 0.06, unpaired t-test, $^{**}P < 0.01$).
(N) The PFC θ phase-locked HPC PNs exhibited increased firing rate at trial 1. As a control, no such alteration was observed in other non-phase-locked HPC PNs (Paired t-test, phase-locked HPC PNs: n = 46, $^{*}P < 0.05$; non-phase-locked HPC PNs: n = 99, $P = 0.66$).
(O) HPC PNs that phase-locked to PFC θ demonstrated more prominent θ-spike PS, indicated by higher slope than non-phase-locked units (Phase-locked HPC PNs: 0.54 ± 0.05; non-phase-locked HPC PNs: -0.20 ± 0.05, unpaired t-test, $^{*}P < 0.05$).
(P) Quantification of θ-spike PS. Mean Pearson's correlation coefficient ($R^2$) was used to assess the goodness-of-fit in phase-trial regression. HPC PNs that phase-locked to PFC θ elicited higher $R^2$ value than non-phase-locked HPC PNs (Phase-locked HPC PNs: 0.65 ± 0.04; non-phase-locked HPC PNs: 0.46 ± 0.03, unpaired t-test, $^{**}P < 0.01$).

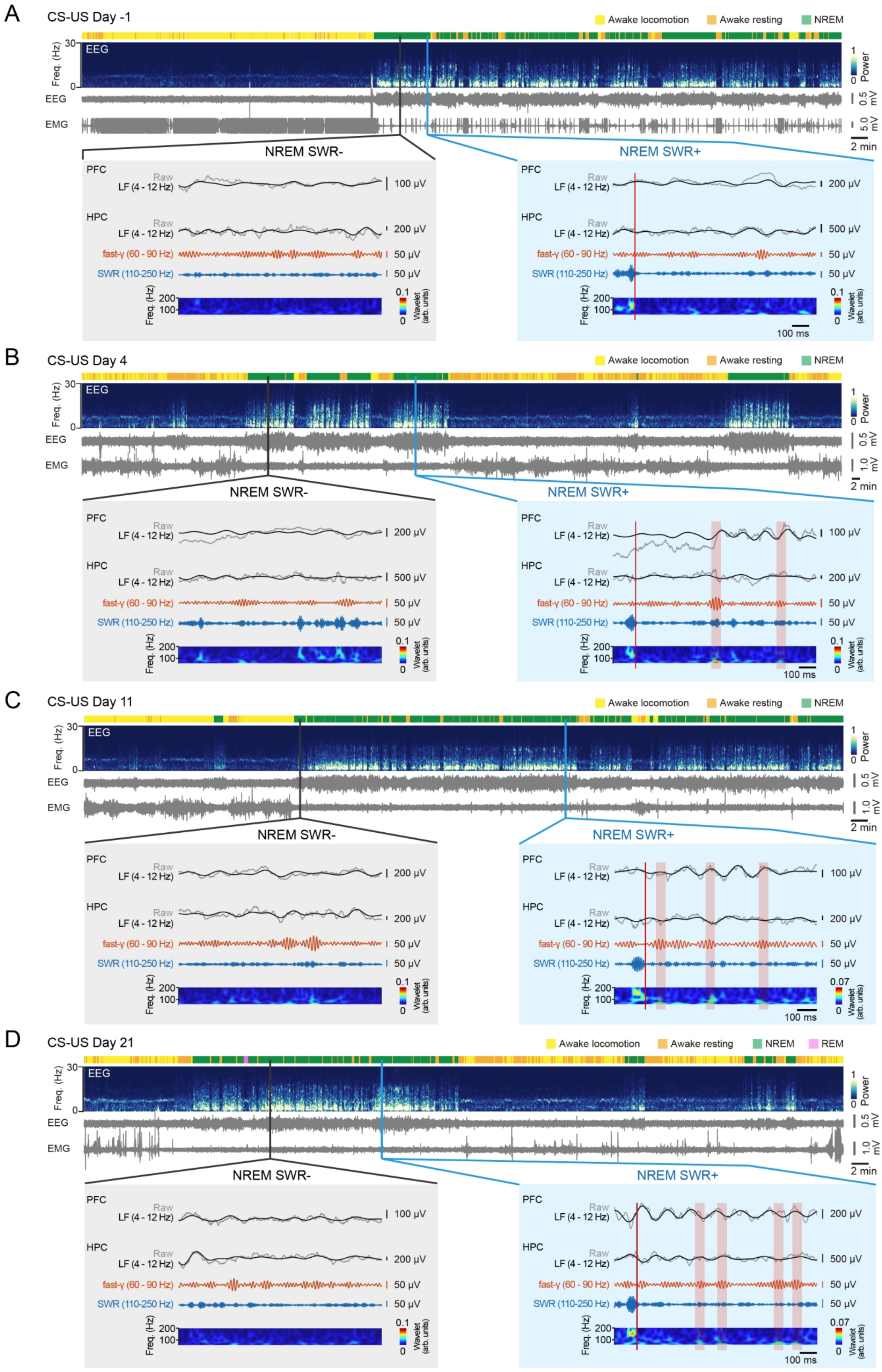

A
CS-US Day -1
Awake locomotion
Awake resting
NREM
EEG
EMG
Freq. (Hz)
Power
2 min
NREM SWR-
NREM SWR+
PFC
HPC
Raw
LF (4 - 12 Hz)
fast-γ (60 - 90 Hz)
SWR (110-250 Hz)
Wavelet (arb. units)
100 ms
B
CS-US Day 4
C
CS-US Day 11
D
CS-US Day 21
REM

**Figure S7. [Temporal SWR-associated coupling dynamics in HPC-PFC circuit revealed PAS across different timepoints during system consolidation], related to Figure 3.**

(A-D) State classification and trace example for CS-US group in Day -1 (A), Day 4 (B), Day 11(C) and Day 21 (D). (Top) Yellow, orange, green, pink (for D) bars indicate awake locomotion, awake resting, NREM and REM, respectively. (Middle) Time-resolved power spectra and waveforms of EEG and EMG. (Bottom) Sample time-aligned LFP traces and wavelet transforms recorded during NREM, showing SWR- (left) and SWR+ (right) segments in PFC and HPC. For each inset (SWR- or SWR+), traces are shown from top to bottom as follows: PFC LFP raw trace (gray) and LF band-pass (4-12 Hz, black); HPC LFP raw trace (gray), LF band-pass (4-12 Hz, black), fast-γ band-pass (60-90 Hz, orange), and ripple band-pass (110-250 Hz, blue); the corresponding power spectrum indicates the temporal positions of ripple and fast-γ activity. Orange boxes highlight representative fast-γ events. Red vertical lines indicate the end time of each SWR event. arb., arbitrary. LF, low frequency.

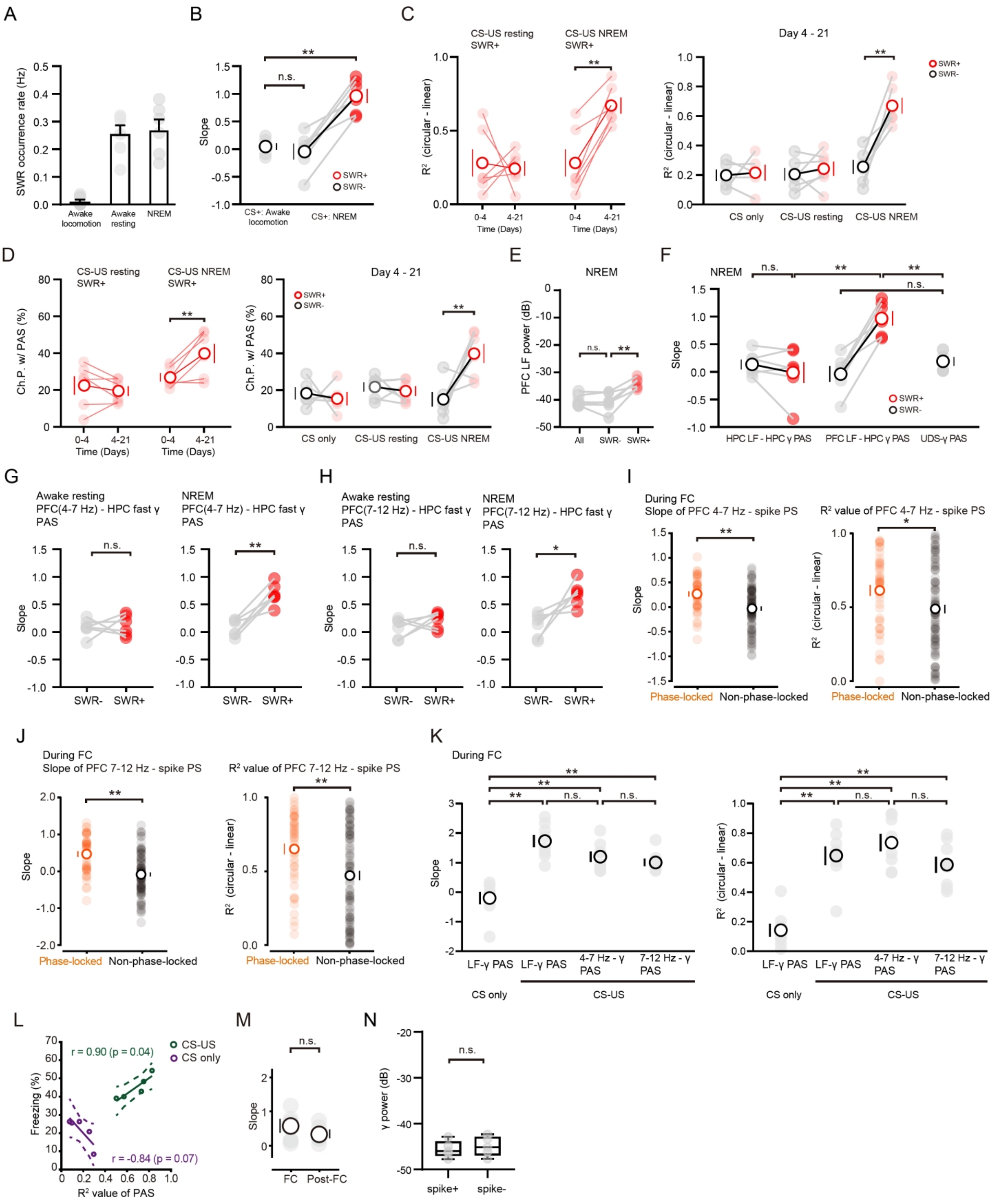

**Figure S8. [Quantification of PAS across multiple frequency bands and inter-regional pairings], related to Figure 3.**

(A) SWR occurrence rate in different states of animals (Awake locomotion:0.01 ± 0.007; Awake resting 0.25 ± 0.03; NREM: 0.27 ± 0.04).

(B) No PAS in Awake locomotion of CS-US group during day 4-21 post-FC, CS-US Awake locomotion: 0.05 ± 0.06, CS-US NREM SWR−: -0.04 ± 0.14; CS-US NREM SWR+: 0.96 ± 0.12; two-sided unpaired t-test, $^{**}P < 0.01$. n.s., no significant, mean ± s.e.m.

(C) (Left) Statistical analysis of mean $R^2$ value showing significant PAS in NREM SWR+ during day 4-21 post-FC compared to day 0-4. (Resting SWR+ day 0-4: 0.28 ± 0.09, day 4-21: 0.24 ± 0.05; two-sided paired t-test, t = 0.28, n.s., no significant, NREM SWR+ day 0-4: 0.28 ± 0.09, day 4-21: 0.67 ± 0.05; two-sided paired t-test, t = 5.59, $**P < 0.01$. Mean ± s.e.m.) (Right) Statistical analysis of mean $R^2$ value showing significant PAS in NREM SWR+ during day 4-21 post-FC compared to CS only, resting and SWR- group. (CS only SWR−: 0.20 ± 0.04, CS only SWR+: 0.26 ± 0.05; CS-US resting SWR−: 0.21 ± 0.04; CS-US resting SWR+: 0.24 ± 0.05; CS-US NREM SWR−: 0.28 ± 0.05; CS-US NREM SWR+: 0.67 ± 0.05; Repeated measures (RM) one-way ANOVA with Tukey post-hoc test, $F(5,28) = 0.98$, $**P < 0.01$. Mean ± s.e.m.)
(D) (Left) Statistical analysis of percentage of channel pairs showing significant PAS in NREM SWR+ during day 4-21 post-FC compared to day 0-4. (Resting SWR+ day 0-4: 22.46 ± 4.89%, day 4-21: 19.47 ± 2.60%; two-sided paired t-test, t = 0.92, n.s., no significant, NREM SWR+ day 0-4: 26.89 ± 2.20, day 4-21: 39.84 ± 5.11; two-sided paired t-test, t = 3.25, $**P < 0.01$. Mean ± s.e.m.) (Right) Statistical analysis of percentage of channel pairs showing significant PAS in NREM SWR+ during day 4-21 post-FC compared to CS only, resting and SWR- group. (CS only SWR−: 18.28 ± 3.32%, CS only SWR+: 15.39 ± 3.60%; CS-US resting SWR−: 21.74 ± 2.50%; CS-US resting SWR+: 19.47 ± 2.6%; CS-US NREM SWR−: 15.04 ± 3.93; CS-US NREM SWR+: 39.84 ± 5.11; Repeated measures (RM) one-way ANOVA with Tukey post-hoc test, $F(5,28) = 0.96$, $**P < 0.01$. Mean ± s.e.m.)
(E) Histograms show increasing of PFC LF power in NREM SWR+ compared to NREM SWR- after conditioning. While overall LF power did not change. All: -40.35 ± 0.86, SWR−: -40.78 ± 1.42, SWR+: -34.03 ± 0.76; two-sided paired t-test. $**P < 0.01$, n.s., no significant, mean ± s.e.m. LF, low frequency.
(F) No within-HPC PAS in the CS-US NREM and No UDS-γ PAS. (Within-HPC PAS SWR-: 0.13 ± 0.09, Within-HPC PAS SWR+: -0.02 ± 0.19; CS-US NREM SWR−: -0.04 ± 0.14; CS-US NREM SWR+: 0.96 ± 0.12; USD-γ PAS: 0.14 ± 0.08; Repeated measures (RM) one-way ANOVA with Tukey post-hoc test, $F(5,20) = 5.434$, $**P < 0.01$. Mean ± s.e.m.)
(G) (Left) No PFC (4-7 Hz)- HPC fast-γ PAS in awake resting of CS-US group during day 4-21 post-FC, SWR-: 0.08 ± 0.06, SWR+: 0.12 ± 0.08; two-sided paired t-test. (Right) High occurrence of PFC (4-7 Hz)- HPC fast-γ PAS in NREM SWR+ of CS-US group during day 4-21 post-FC, as indicated by increased slope of PAS showing prominent phase shift. SWR-: 0.07 ± 0.06, SWR+: 0.69 ± 0.08; two-sided paired t-test. $**P < 0.01$.
(H) (Left) No PFC (4-7 Hz)- HPC fast-γ PAS in Awake resting of CS-US group during day 4-21 post-FC, SWR-: 0.07 ± 0.07, SWR+: 0.21 ± 0.06; two-sided paired t-test. (Right) High occurrence of PFC (4-7 Hz)- HPC fast-γ PAS in NREM SWR+ of CS-US group during day 4-21 post-FC, as indicated by increased slope of PAS showing prominent phase shift. SWR-: 0.12 ± 0.09, SWR+: 0.56 ± 0.17; two-sided paired t-test. $*P < 0.05$. n.s., no significant, mean ± s.e.m.
(I) (Left) HPC units that phase-locked to PFC 4-7 Hz demonstrated more prominent PFC 4-7 Hz-spike PS, indicated by higher slope than non-phase-locked units (Phase-locked HPC units: 0.35 ± 0.07; non-phase-locked: -0.04 ± 0.05, unpaired t-test, $**P < 0.01$). (Right) Quantification of PFC 4-7 HZ-spike PS. Mean Pearson's correlation coefficient ($R^2$) was used to assess the goodness-of-fit in phase-trial regression. HPC units that phase-locked to PFC 4-7 HZ elicited higher $R^2$ value than non-phase-locked units (Phase-locked HPC units: 0.62 ± 0.04; non-phase-locked: 0.49 ± 0.03, unpaired t-test, $*P < 0.05$).

(J) (Left) HPC units that phase-locked to PFC 7-12 Hz demonstrated more prominent PFC 7-12 HZ-spike PS, indicated by higher slope than non-phase-locked units (Phase-locked HPC units: 0.45 ± 0.07; non-phase-locked: -0.10 ± 0.05, unpaired t-test, $^{**}P < 0.01$). (Right) Quantification of PFC 7-12 HZ-spike PS. Mean Pearson's correlation coefficient ($R^2$) was used to assess the goodness-of-fit in phase-trial regression. HPC units that phase-locked to PFC 7-12 HZ elicited higher $R^2$ value than non-phase-locked units (Phase-locked HPC units: 0.65 ± 0.04; non-phase-locked: 0.47 ± 0.03, unpaired t-test, $^{*}P < 0.05$).
(K) (Left) Quantification of PAS with slope of phase shift. CS-US group, which included three subgroups (PAS, 4-7 Hz-γ PAS and 7-12 Hz-γ PAS), elicited higher slope than that of CS only group (CS only: -0.20 ± 0.21; PAS: 1.73 ± 0.2; 4-7 Hz-γ PAS: 1.20 ± 0.17; 7-12 Hz-γ PAS: 1.00 ± 0.12, Repeated measures (RM) one-way ANOVA with Tukey post-hoc test, $F(7,21) = 0.59$, $^{**}P <$ 0.01. n.s., no significant, mean ± s.e.m.). (Right) Statistical analysis of mean $R^2$ value. CS-US group, which included three subgroups (PAS, 4-7 Hz-γ PAS and 7-12 Hz-γ PAS), elicited significant PAS than that of CS only group (CS only: 0.14 ± 0.04; PAS: 0.65 ± 0.06; 4-7 Hz-γ PAS: 0.74 ± 0.06; 7-12 Hz-γ PAS: 0.59 ± 0.05, Repeated measures (RM) one-way ANOVA with Tukey post-hoc test, $F(7, 21) = 0.60$, $^{**}P < 0.01$. n.s., no significant, mean ± s.e.m.).
(L) The consolidated memory assessed at day 28 post-FC was highly correlated with reoccurring PAS in the CS-US group, as indicated by Pearson correlation coefficient of recalled freezing behavior and the extent of PAS defined as $R^2$ value of PAS (CS only: $r = -0.84$, $P = 0.07$; CS-US: $r = 0.90$, $P = 0.04$). Each cycle represents data from an individual animal.
(M) No significant difference between slopes of LF-spike PS at conditioning (data from Figure 2F) and at consolidation stages (Conditioning: n = 5, post-FC: n = 5; unpaired t-test, $P = 0.36$, center values denote mean ± s.e.m.).
(N) Histograms show no significant difference of HPC fast-γ power between segments surround spike or without spike (Spike+: -47.57 ± 0.82; Spike−: -44.94 ± 0.96, unpaired t-test, $P = 0.63$).

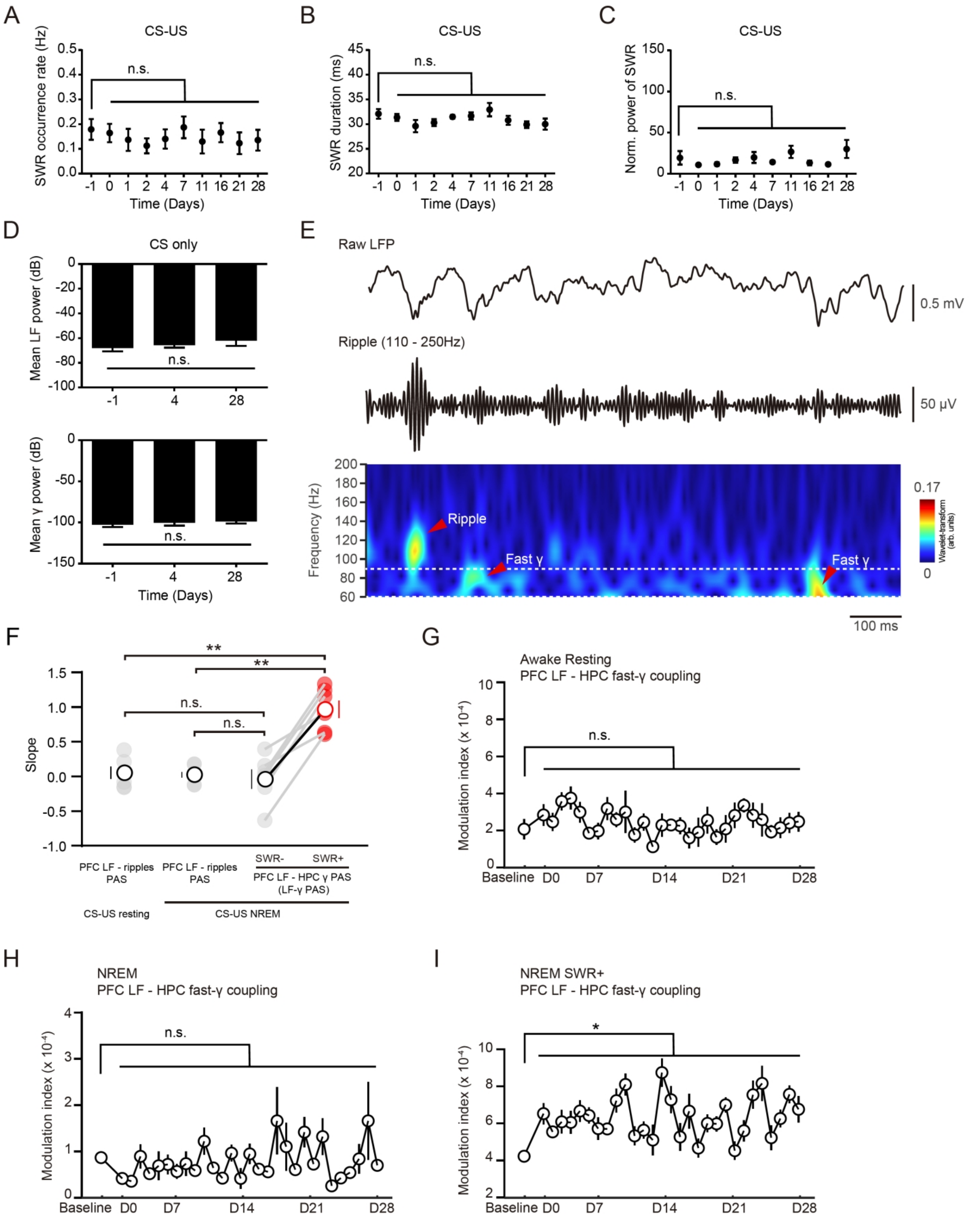

**Figure S9. [Properties of SWR and its associated oscillatory activity], related to the main text.** (A-C) Time courses of SWR frequency (A), SWR duration (B), SWR power (normalized, C). No significant difference across days indicated stable recording quality. n = 4, mean ± s.e.m

(D) Mean LF and γ LFP power of CS only group recording from day -1, day 4 and day 28. n.s., no significant; n = 5, mean ± s.e.m. LF, low frequency.

(E) Raw LFP trace from the HPC region after training (above), filtered ripple trace (110-250 Hz middle) and the corresponding power spectrum shows the temporal positions of ripple and fast-γ (bottom). arb., arbitrary.

(F) No PFC LF-ripples PAS both in the CS-US NREM and CS-US resting period (CS-US resting PFC LF-ripples PAS: 0.05 ± 0.09, CS-US NREM PFC LF-ripples PAS: 0.02 ± 0.04; CS-US NREM SWR−: -0.04 ± 0.14; CS-US NREM SWR+: 0.96 ± 0.12; Repeated measures (RM) one-way ANOVA with Tukey post-hoc test, $F(5,15) = 2.97$, $^{**}P < 0.01$. Mean ± s.e.m).

(G and H) During the resting state, the modulation index of PFC-HPC LF-γ PAC level across consolidation remains unchanged. (Repeated measures (RM) one-way ANOVA with Tukey post-hoc test, resting (G): $F(5, 145) = 1.371$. NREM (H): $F(5, 145) = 1.148$. n.s., no significant, mean ± s.e.m.)

(I) the level of PAC associated with SWR was significantly increased throughout days 0-28 following training. Repeated measures (RM) one-way ANOVA with Tukey post-hoc test, $F(29, 145) = 2.30$, $^{*}P < 0.05$. n.s., no significant, mean ± s.e.m.

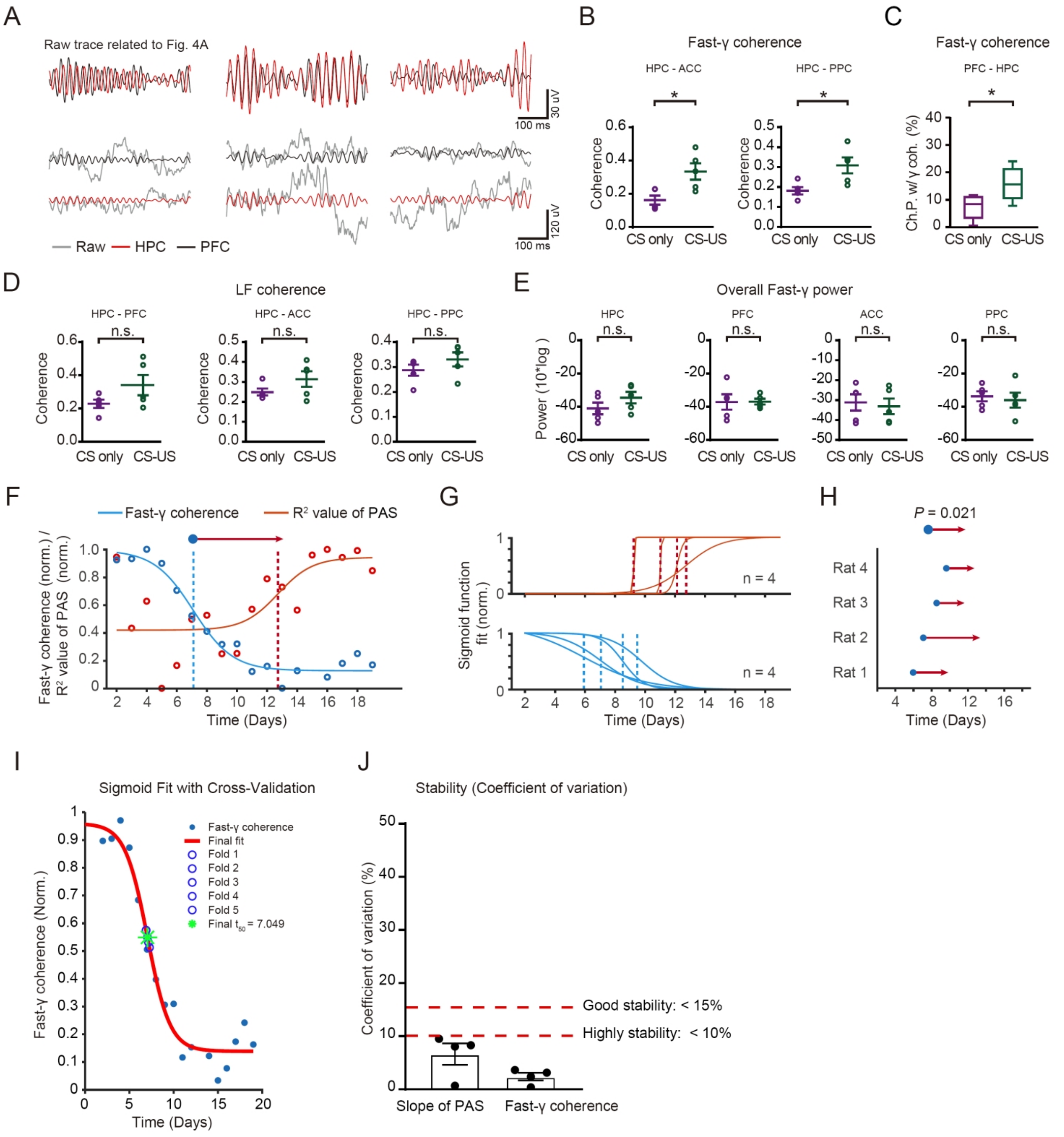

**Figure S10. [Statistical analysis of HPC-cortical coherence and γ power at recent stage], related to Figure 4.**

(A) Representative raw and fast-γ band-pass-filtered trace sample related to **Figure 4A**.

(B) Histograms show increased fast-γ coherence in HPC-ACC and HPC-PPC circuits at recent stage, two-sided unpaired t-test, *$P < 0.05$.

(C) Histograms show increased percentage of channel pairs showing HPC-PFC γ coherence (Ch.P. w/γ coh.) at recent stage in conditioned CS-US group. Mean γ coherence ± s.e.m. of 6 rats from CS only group (magenta), 6 rats from CS-US group (green), Bonferroni post-hoc test, *$P < 0.05$, CS-US group versus CS only group from d0 to d7, center values denote mean ± s.e.m.

(D) Histograms show no significant difference of LF coherence in HPC-PFC, HPC-ACC and

HPC-PPC circuits between control (CS only) and conditioned (CS-US) animals at recent stage. LF, low frequency.

(E) Histograms show no significant difference of fast-γ power in HPC, PFC, ACC and PPC between control (CS only) and conditioned (CS-US) animals at recent stage.

(F) Comparison of gradual increase of $R^2$ value of PAS with gradual drop in HPC-PFC γ coherence in single animal.

(G) Sigmoid function fits for each animal (n = 4).

(H) Sharpest changes marked by blue circles and red arrowheads. Significant for 4 animals; two-sided paired t-test, $P = 0.021$.

(I) Sample sigmoid fitting curves from the validation set of fast-γ coherence. Blue open circles: $t_{50}$ values calculated from folds 1-5; green stars: $t_{50}$ for the complete dataset.

(J) The coefficients of variation for the slope of PAS and fast-γ coherence are both below 15%, indicating good stability.

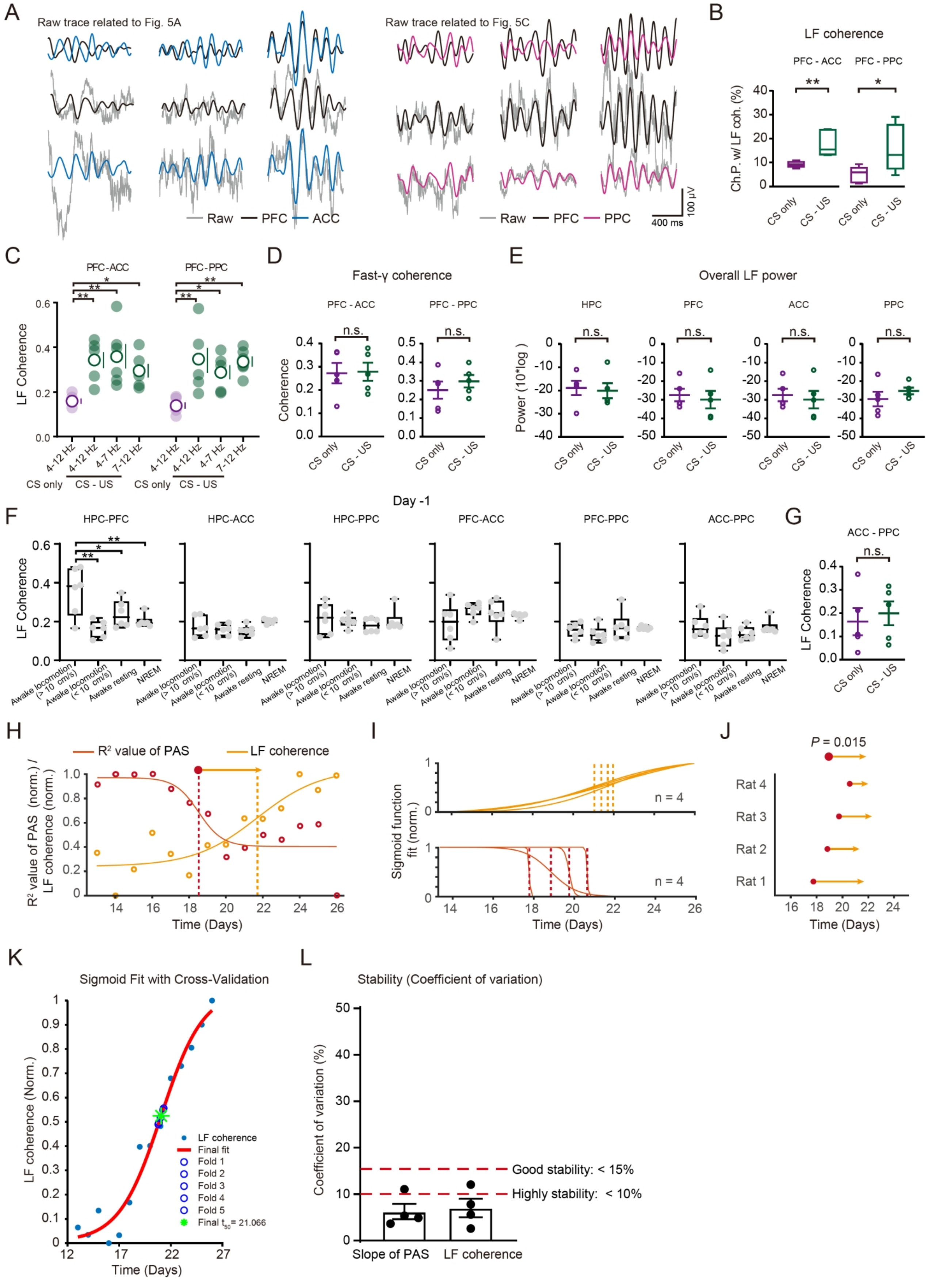

**Figure S11. [Statistical analysis of inter-cortical coherence and low-frequency power at remote**

**stage], related to Figure 4.**

(A) Representative raw and LF band-pass-filtered trace sample related to **Figure 4I** (left) and **Figure 4K** (right).

(B) Histograms show increased percentage of channel pairs showing PFC-ACC and PFC-PPC low-frequency coherence (Ch.P. w/LF coh.) at remote stage in conditioned CS-US group. Mean γ coherence ± s.e.m. of 6 rats from CS only group (magenta), 6 rats from CS-US group (green), Bonferroni post-hoc test, $*P < 0.05$, CS-US group versus CS only group from d21 to d28, center values denote mean ± s.e.m.. LF, low frequency.

(C) Histograms show increased PFC-ACC, PFC-PPC LF coherence at recent stage in conditioned CS-US 3 groups (PAS, 4-7 Hz-γ PAS and 7-12 Hz-γ PAS). Mean LF coherence ± s.e.m. of 6 rats from CS only group (magenta), 6 rats from CS-US group (green) (PFC-ACC: CS only: 0.16 ± 0.01; PAS: 0.34 ± 0.04; 4-7 Hz-γ PAS: 0.36 ± 0.05; 7-12 Hz-γ PAS: 0.30 ± 0.03, $F(5,15) = 0.80$; PFC-PPC: CS only: 0.14 ± 0.01; PAS: 0.34 ± 0.06; 4-7 Hz-γ PAS: 0.29 ± 0.03; 7-12 Hz-γ PAS: 0.34 ± 0.02, $F(5,15) = 2.99$; Repeated measures (RM) one-way ANOVA with Tukey post-hoc test, $*P < 0.05$, $**P < 0.01$. Mean ± s.e.m.).

(D) Histograms show no significant difference of γ coherence in PFC-ACC and PFC-PPC circuits between control (CS only) and conditioned (CS-US) animals at remote stage.

(E) Histograms show no significant difference of LF power in HPC, PFC, ACC and PPC between control (CS only) and conditioned (CS-US) animals at remote stage.

(F) Histograms show no significant difference of LF coherence in ACC-PPC circuits between control (CS only) and conditioned (CS-US) animals at remote stage.

(G) Histograms show no significant difference of LF coherence in HPC-PFC, HPC-ACC, HPC-PPC, PFC-ACC, PFC-PPC and ACC-PPC circuits between different states (Awake locomotion, Awake resting and NREM) at day -1.

(H) Comparison of gradual increase of PFC-ACC LF coherence with gradual drop in $R^2$ value of PAS in single animal.

(I) Sigmoid function fits for each animal (n = 4).

(J) Sharpest changes marked by red circles and orange arrowheads. Significant for 4 animals; two-sided paired t-test, $P = 0.015$.

(K) Sample sigmoid fitting curves from the validation set of LF coherence. Blue open circles: $t_{50}$ values calculated from folds 1-5; green stars: $t_{50}$ for the complete dataset.

(L) The coefficients of variation for the slope of PAS and LF coherence are both below 15%, indicating good stability.

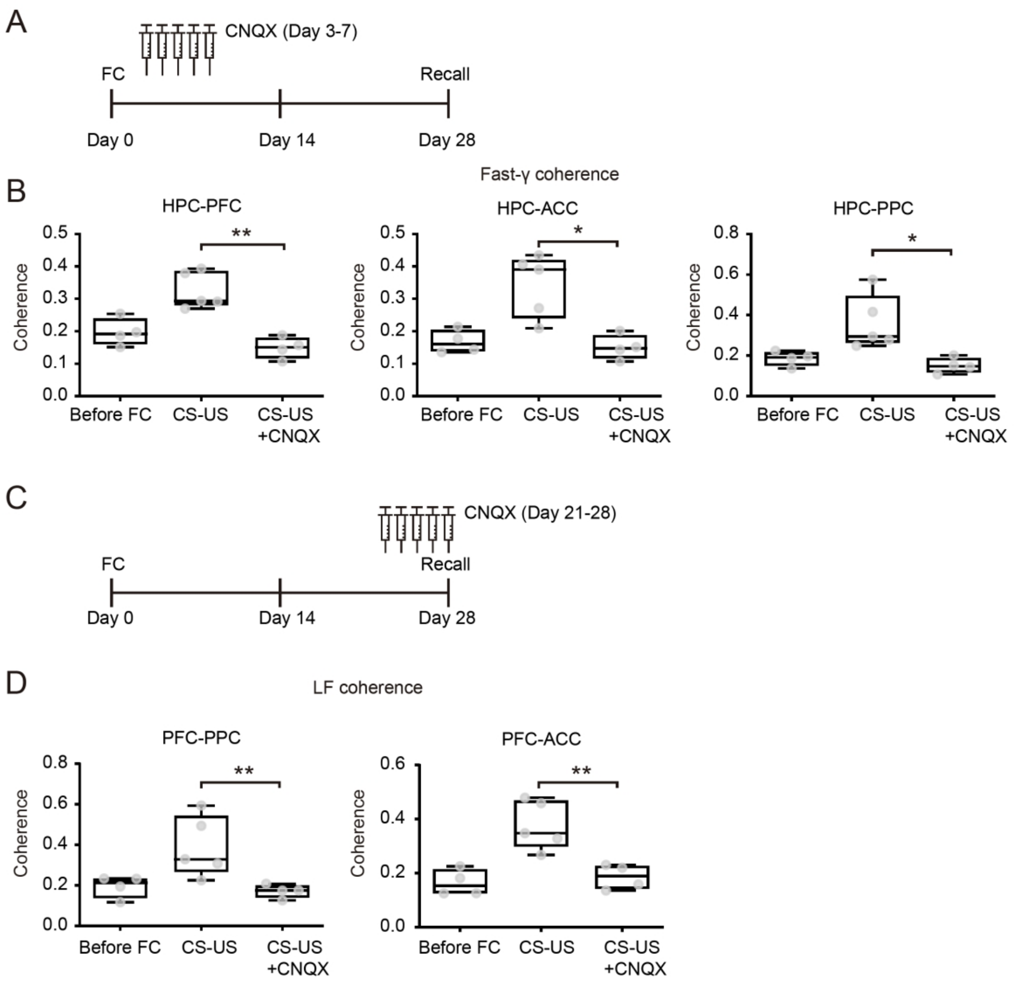

**Figure S12. [Pharmacological inactivation experiments validated HPC and PFC's role as the cross-area dialogue organizers], related to the main text and STAR Methods.**

(A) Experimental timeline for pharmacological inactivation of HPC with CNQX during day 3-7 post-FC.

(B) Application of CNQX in HPC decoupled fast-γ coherence in HPC-PFC, HPC-ACC, and HPC-PPC circuits during day 3-7 post-FC. (Left, HPC-PFC) Before FC: 0.20 ± 0.02; CS-US: 0.33 ± 0.03; CS-US + CNQX: 0.15 ± 0.02, $F(2, 10) = 17.64$. (Middle, HPC-ACC) Before FC: 0.17 ± 0.02; CS-US: 0.34 ± 0.04; CS-US + CNQX: 0.15 ± 0.02, $F(2, 10) = 11.31$. (Right, HPC-PPC) Before FC: 0.19 ± 0.02; CS-US: 0.36 ± 0.06; CS-US + CNQX: 0.15 ± 0.02, $F(2, 10) = 7.29$. Repeated measures (RM) one-way ANOVA with Tukey post-hoc test, *$P < 0.05$, **$P < 0.01$. Mean ± s.e.m.

(C) Experimental timeline for pharmacological inactivation of PFC with CNQX during day 21-28 post-FC.

(D) Application of CNQX in PFC decoupled LF coherence in PFC-ACC and PFC-PPC circuits during day 14-21 post-FC. (Left, PFC-PPC) Before FC: 0.19 ± 0.03; CS-US: 0.39 ± 0.07; CS-US + CNQX: 0.17 ± 0.02, $F(2, 10) = 6.51$. (Right, PFC-ACC) Before FC: 0.16 ± 0.02; CS-US: 0.38 ± 0.04; CS-US + CNQX: 0.19 ± 0.02, $F(2, 10) = 13.68$. Repeated measures (RM) one-way ANOVA with Tukey post-hoc test, **$P < 0.01$. Mean ± s.e.m. LF, low frequency.

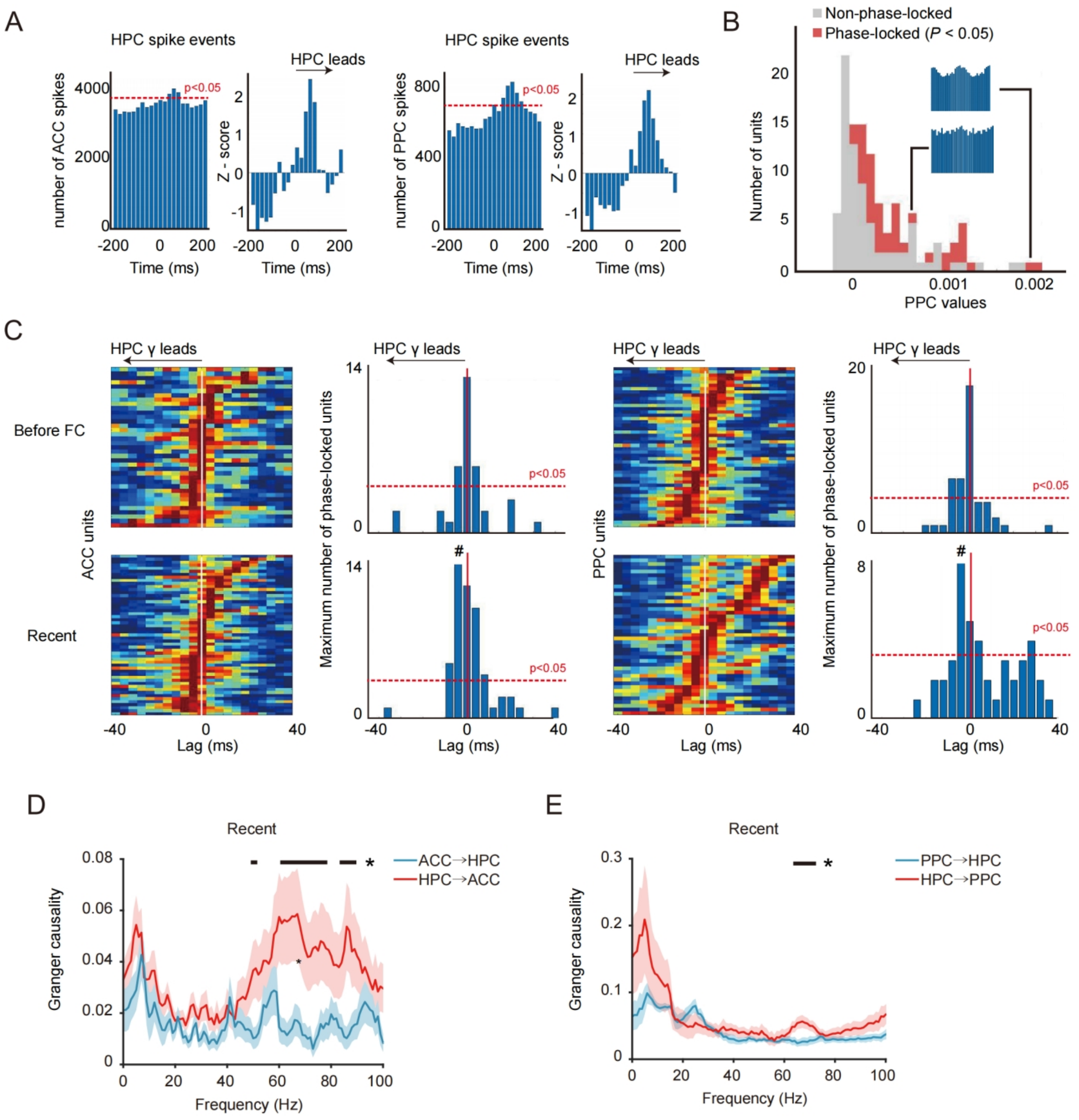

**Figure S13. [Spike-spike correlation, pairwise phase consistency and granger causality analyses of HPC-ACC and HPC-PPC circuits], related to Figure 5.**

(A) Spike-spike correlation in HPC-ACC and HPC-PPC circuits at recent stage. A total of 289 HPC-ACC and 323 HPC-PPC spike pairs were analyzed, with significant HPC-leading interactions observed in 157 and 191 pairs, respectively. Red dashed lines indicate significance level estimated by spike jitter test ($P < 0.05$).

(B) Distribution of phase locking values for all units from spikes recorded, colored by significance (Rayleigh's test, $P < 0.05$). Insets, HPC γ phase histograms from example units.

(C) Pseudocolour plot of normalized pairwise phase consistency values of ACC (left) and PPC (right). ACC and PPC units significantly phase-locked to HPC γ sorted by lag of maximal phase-locking during recent stage after conditioning. Vertical red line, zero lag. Hash indicates mean lag. Horizontal red dashed lines, chance.

(D and E) Granger causality analysis based on LFPs in HPC-ACC (D) and HPC-PPC (E) circuit at recent stage. n = 5 rats, multiple t-tests, Black lines indicate $*P < 0.05$.

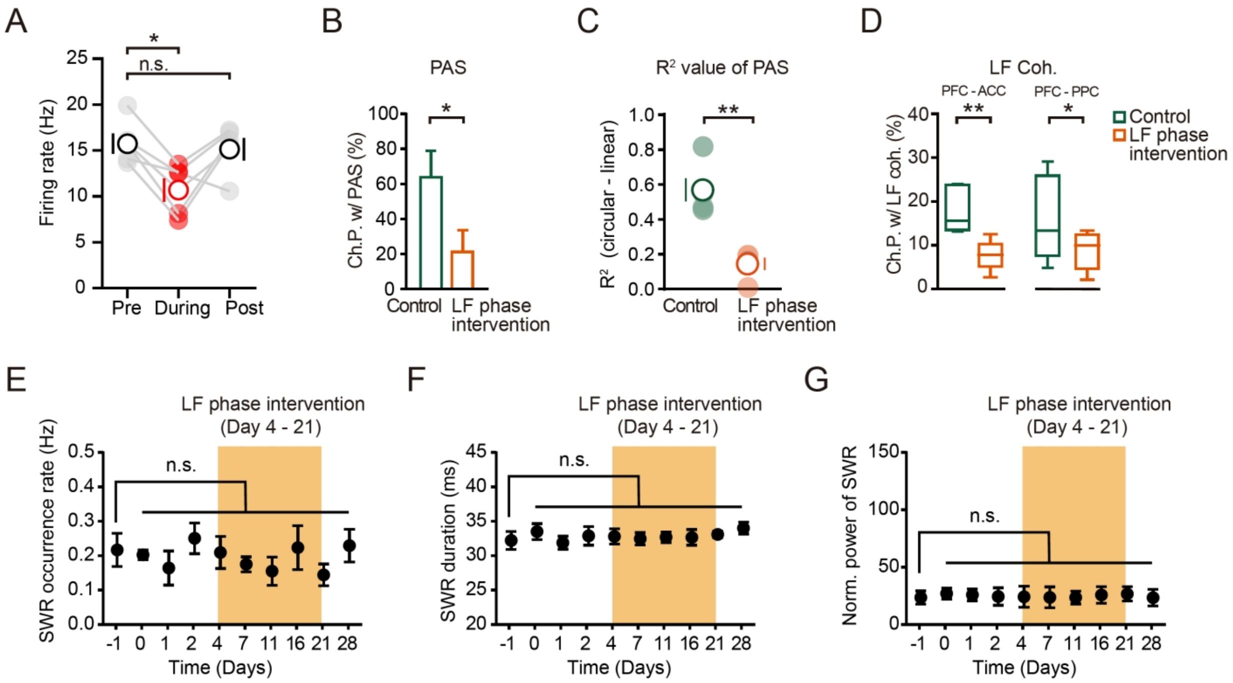

**Figure S14. [Effects of random LF phase intervention and SWR properties validation], related to Figure 6.**

(A) PFC neuronal firing was decreased during opto-stimulation, but soon recovered post stimulation. Repeated measures (RM) one-way ANOVA with bonferroni post-hoc test, F(4,8) = 1.64, paired t-tests, $*P < 0.05$.

(B) Random LF intervention suppresses reoccurrence of PAS, as indicated by decreased percentage of channel pairs showing PAS (Ch.P. w/PAS; Control: 63.3 ± 14.6, LF phase intervention, 21.2 ± 12.6, unpaired t-test, $*P < 0.05$). LF, low frequency; Ch.P. channel pairs.

(C) Random LF intervention suppresses reoccurrence of PAS, as indicated by decreased $R^2$ (Control: 0.57 ± 0.07, LF phase intervention, 0.14 ± 0.03, unpaired t-test, $**P < 0.01$).

(D) Significantly decreased channel pairs showing PAS (Ch.P. w/PAS) in PFC-ACC and PFC-PPC circuits at remote stage (unpaired t-test, $*P < 0.05$, $**P < 0.01$).

(E-G) Time courses of SWR frequency (E), SWR duration (F), SWR power (normalized, G). No significant difference across days before and after PFC LF intervention (day 4-21).

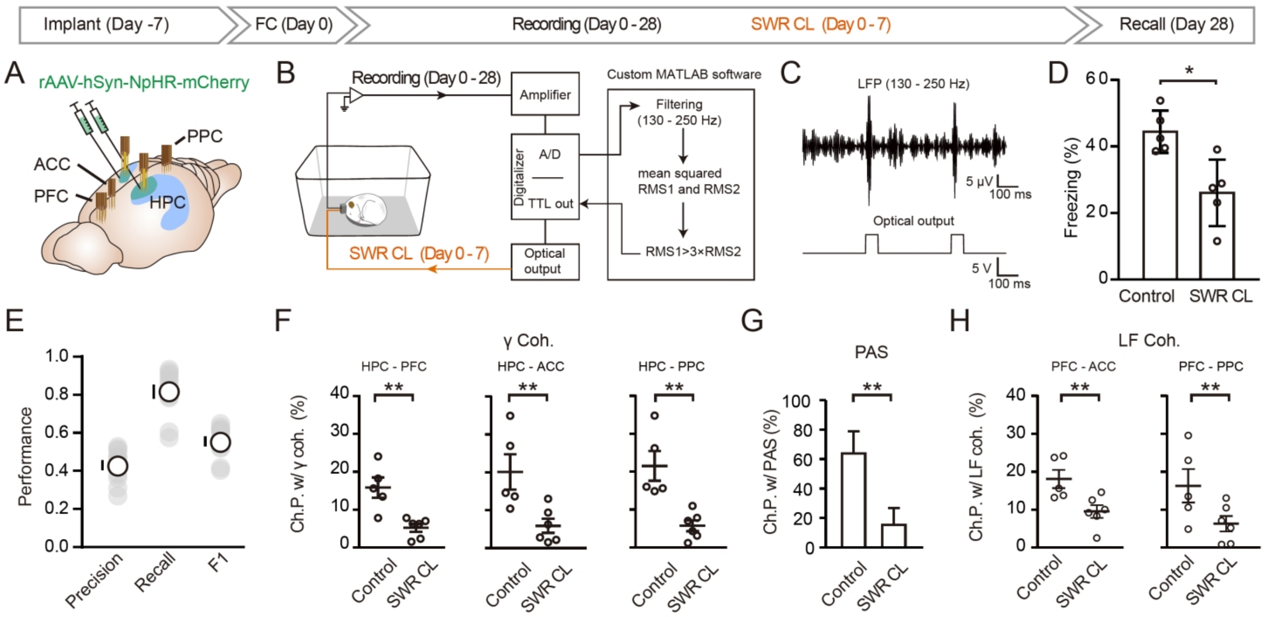

**Figure S15. [Hippocampal SWR is required for HPC-cortical γ coherence, PAS, PFC-cortical LF coherence and remote memory consolidation], related to the main text.**

(A) Electrodes targeting and daily experimental schedule.

(B) Closed-loop strategy to intervene SWR (SWR CL) at recent stage via optogenetic deactivation of HPC neurons.

(C) SWR online detection and signal triggering diagram.

(D) The closed-loop SWR intervention caused impaired remote memory (*$P$ < 0.05, unpaired t-test).

(E) The closed-loop SWR intervention was validated using precision (the ratio of correct predictions to total predictions), recall (the ratio of correct predictions to all offline-detected events) and F1 score (the harmonic mean of precision and recall). n = 11 sessions from four rats.

(F) Histograms show SWR closed-loop intervention implemented at recent stage suppress HPC-cortical γ coherence (unpaired t-test, **$P$ < 0.01).

(G) SWR closed-loop intervention suppresses subsequent reoccurrence of PAS (unpaired t-test, **$P$ < 0.01).

(H) Histograms show decreased LF coherence following SWR closed-loop intervention in HPC-ACC and HPC-PPC circuits at remote stage (unpaired t-test, **$P$ < 0.01).

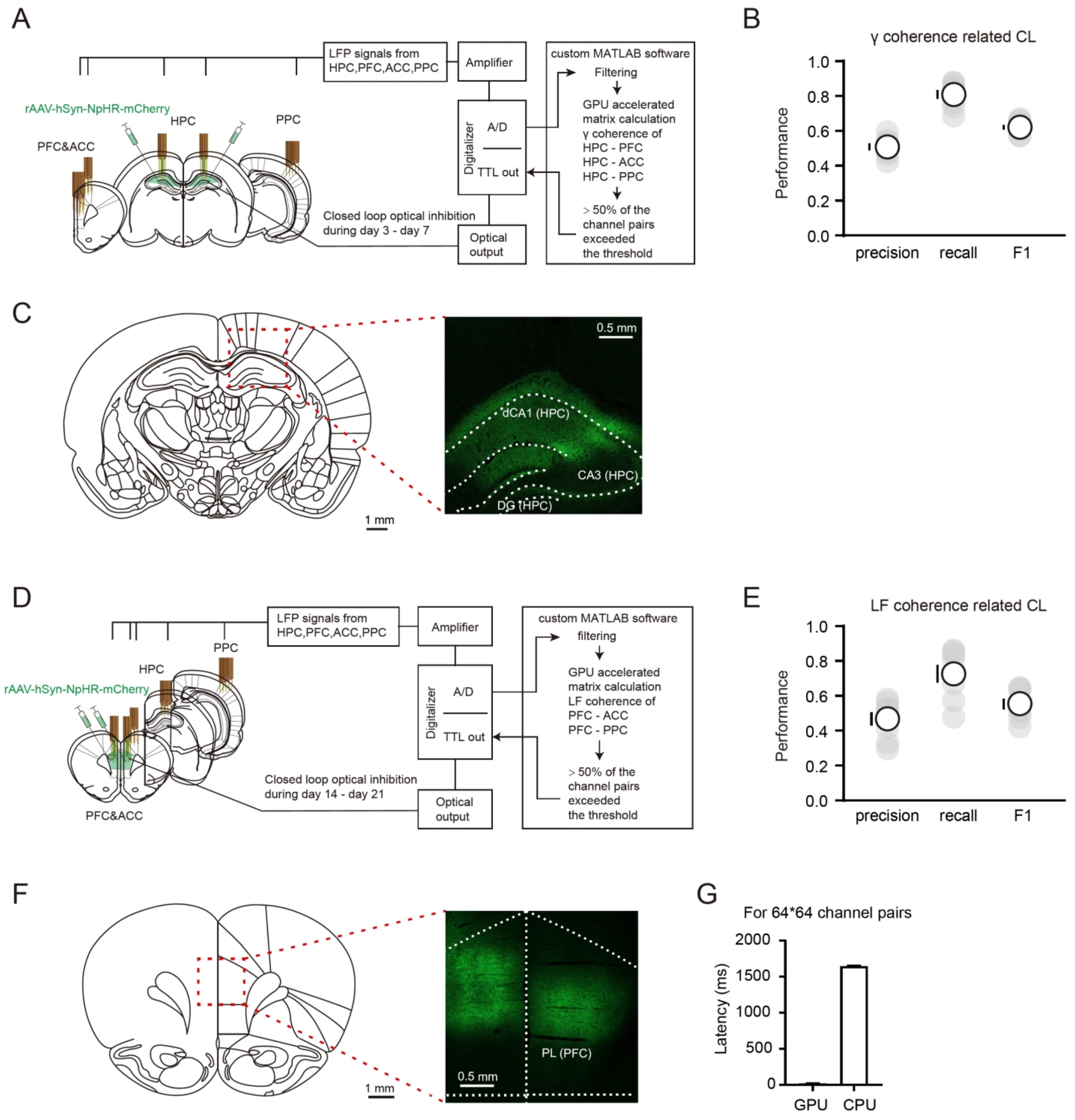

**Figure S16 [Validation of inter-regional coherence-driven closed-loop intervention], related to Figure 7 and STAR Methods.**

(A) Closed-loop strategy to intervene HPC-cortical γ coherence at recent stage via optogenetic inhibition of HPC.

(B) The closed-loop γ coherence intervention was validated using precision (0.51 ± 0.02; the ratio of correct predictions to total predictions), recall (0.81 ± 0.03; the ratio of correct predictions to all offline-detected events) and the F1 score (0.62 ± 0.01; the harmonic mean of precision and recall) (n = 8 sessions from four rats).

(C) Representative micrograph of HPC eNpHR expression. Dashed box in schematic coronal section indicated the dorsal CA1 of HPC corresponding to the fluoresce image.

(D) Closed-loop strategy to intervene PFC-cortical LF coherence at recent stage via optogenetic inhibition of PFC. LF, low frequency.

(E) The closed-loop LF coherence intervention was validated using precision (0.46 ± 0.04), recall (0.73 ± 0.05) and the F1 score (0.55 ± 0.03) (n = 8 sessions from four rats).
(F) Representative micrograph of PFC eNpHR expression. Dashed box in schematic coronal section indicated PL of PFC corresponding to the fluoresce image. PL, prelimbic cortex.
(G) Histograms for GPU-based and CPU-based parallel computing for 64*64 channel pairs (GPU: 20.2 ± 0.7 ms; CPU: 1649 ± 1.0 ms).

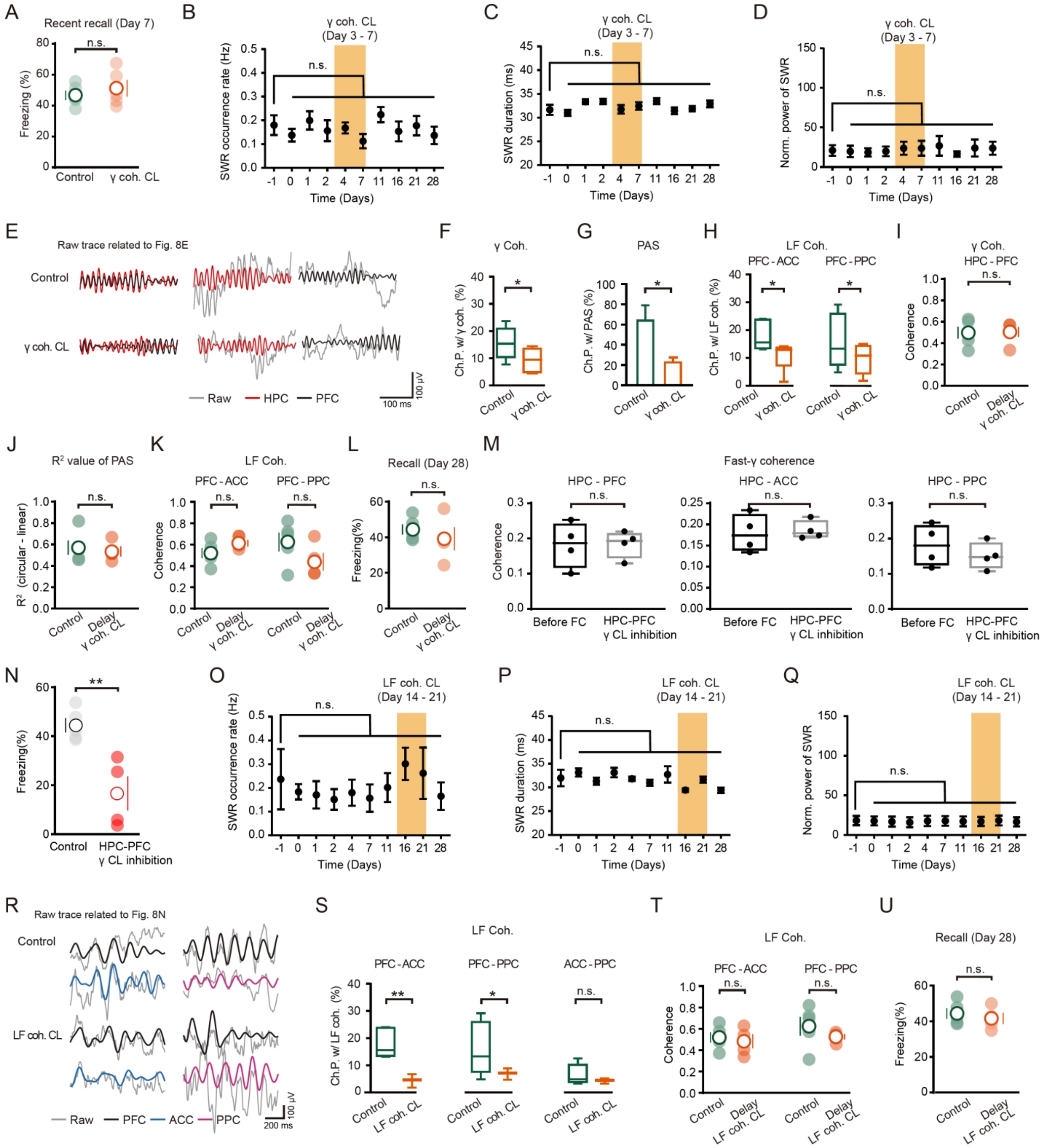

**Figure S17. [Effects of coherence-based closed-loop interventions on retention of fear memory, SWR properties, PAS and inter-regional coherence], related to Figure 7.**

(A) Statistical histogram shows close-loop γ coherence intervention did not affect recent memory. n.s., no significant, unpaired t-test.

(B-D) Time courses of SWR frequency (B), SWR duration (C), SWR power (normalized; D). No significant difference across days before and after HPC-cortical γ coherence related closed loop inhibition (Day 3-7).

(E) Representative raw and fast-γ band-pass-filtered trace sample related to **Figure 7E**.

(F) Histograms show decreased percentage of channel pairs exhibiting γ coherence (Ch.P. w/γ coh.) following γ coh. CL (recorded at day 7 post-FC, unpaired t-test, **P* < 0.05).

(G) Histograms show decreased percentage of channel pairs exhibiting PAS reoccurrence (Ch.P. w/ PAS) following γ coh. (Control: 63.3 ± 14.6%, γ coh. CL: 22.4 ± 4.9, unpaired t-test, *$P < 0.05$).
(H) Histograms show decreased percentage of channel pairs exhibiting inter-cortical coherence (Ch.P. w/LF coh.) following γ coh. CL (unpaired t-test, *$P < 0.05$). LF, low frequency.
(I) Histograms show delayed γ coherence (γ coh) closed-loop intervention failed to affect HPC-cortical γ coherence at recent stage.
(J) PAS reoccurrence was unaffected by delayed γ coherence closed-loop intervention.
(K) Inter-cortical LF coherence was unaffected by delayed γ coherence closed-loop intervention.
(L) Statistical histogram shows delayed γ coherence intervention did not affect remote memory.
(M) Histograms show no significant difference of fast-γ coherence in HPC-PFC, HPC-ACC and HPC-PPC circuits after closed-loop inhibition triggered by HPC-PFC γ coherence detection at remote stage.
(N) Closed-loop inhibition triggered by HPC-PFC γ coherence detection caused impaired remote memory (day 28 post-FC; Control, 44.4 ± 2.8; HPC-PFC γ CL inhibition, 16.6 ± 7.0, unpaired t-test, **$P < 0.01$).
(O-Q) Time courses of SWR frequency (O), SWR duration (P), SWR power (normalized, Q). No significant difference across days before and after inter-cortical LF coherence related closed loop inhibition (Day 14-21). The yellow shading represents the time period during which optogenetic intervention is conducted, and the corresponding recordings are made after the intervention ends each day. n.s., no significant, mean ± s.e.m.
(R) Representative raw and LF band-pass-filtered trace sample related to **Figure 7N**.
(S) Closed-loop LF coherence intervention significantly decreased the percentage of channel pairs exhibiting inter-cortical LF coherence (Ch.P. w/LF coh.) at remote stage (unpaired t-test, *$P < 0.05$, **$P < 0.01$).
(T) Inter-cortical LF coherence was unaffected by delayed LF coherence (LF coh.) intervention.
(U) Statistical histogram shows delayed LF coherence intervention did not affect remote memory. n.s., no significant, unpaired t-test.

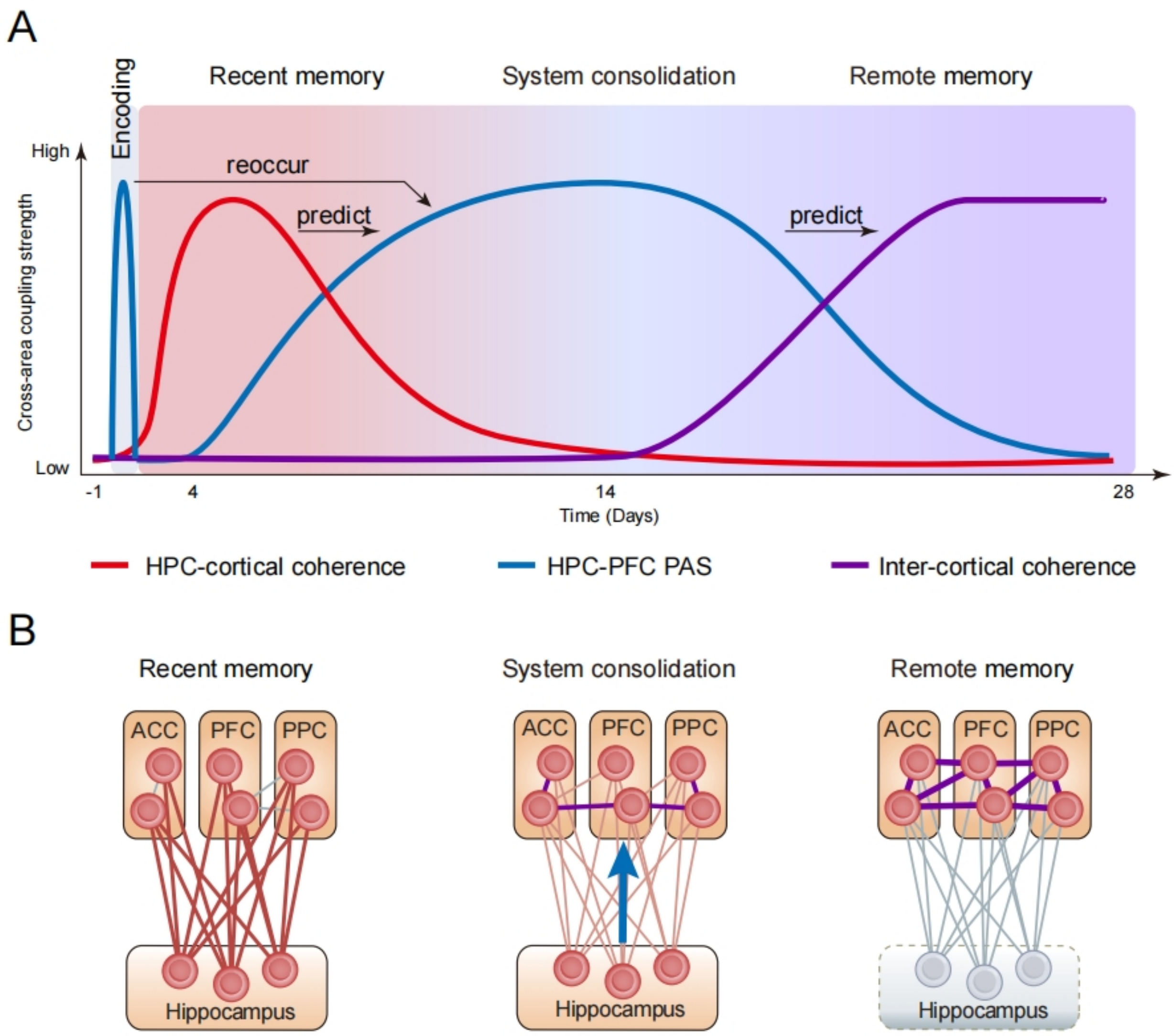

**Figure S18. [Stepwise cross-area oscillatory dialogues drive remote memory formation], related to the main text.**

(A) Schematic illustration showing the temporal sequence of consecutive cross-area oscillatory events during the development of remote memory formation. These events are interconnected and constitute a chain of cross-area oscillatory events potentially crucial to memory consolidation and stabilization. The colored shaded area shows the gradual transition from recent to remote memory.

(B) Working model for memory system consolidation. Memories are successively stored in the form of HPC-cortical coherence (left) and inter-cortical coherence (right), respectively. During memory system consolidation (middle), the reoccurring PAS drives the HPC→cortical memory transfer with subsequent hippocampal independence. Blue arrowhead shows the direction of memory transfer.

**Table S1. [GLM analysis of speed- and learning-dependent modulation of local power and cross-regional coherence], related to Figure S11F.**

| 4-12 Hz power ~ 1 + Speed + Learning + Speed*Learning + (1 \| Animal) | | | | | | | | | | | | |
|---|---|---|---|---|---|---|---|---|---|---|---|---|
| | **PFC** | | | **ACC** | | | **HPC** | | | **PPC** | | |
| | Estimate | t | p | Estimate | t | p | Estimate | t | p | Estimate | t | p |
| Name | -6.04 | -204.52 | 0 | -6.27 | -212.18 | 0 | -2.21 | -59.86 | 0 | -6.12 | -195.27 | 0 |
| Intercept | 1.04 | 12.06 | 2.14e-33 | 0.19 | 2.29 | 0.02 | 1.00 | 3.31 | 9.32e-4 | 0.17 | 1.89 | 0.059 |
| Learning | 0.053 | 6.387 | 1.71e-10 | 0.079 | 10.93 | 8.98e-28 | 0.04 | 5.94 | 2.83e-9 | 0.085 | 10.66 | 1.63e-26 |
| Speed×Learning | 0.027 | 0.24 | 0.80 | 0.27 | 2.62 | 0.0088 | -0.076 | -0.24 | 0.81 | -0.093 | -0.80 | 0.42 |

| 60-90 Hz power ~ 1 + Speed + Learning + Speed*Learning + (1 \| Animal) | | | | | | | | | | | | |
|---|---|---|---|---|---|---|---|---|---|---|---|---|
| | **PFC** | | | **ACC** | | | **HPC** | | | **PPC** | | |
| | Estimate | t | p | Estimate | t | p | Estimate | t | p | Estimate | t | p |
| Name | -9.86 | -362.24 | 0 | -9.28 | -337.97 | 0 | -6.85 | -204.93 | 0 | -9.09 | -172.85 | 0 |
| Intercept | 1.75 | 19.93 | 1.45e-87 | 0.66 | 8.48 | 2.41e-17 | 0.78 | 13.30 | 3.08e-40 | 1.30 | 24.47 | 9.05e-131 |
| Learning | -0.023 | -1.95 | 0.052 | -0.13 | -11.96 | 7.34e-33 | 0.055 | 8.53 | 1.58e-17 | 0.13 | 22.54 | 1.64e-111 |
| Speed×Learning | -0.48 | -1.93 | 0.053 | 0.15 | 0.69 | 0.49 | 0.15 | 1.60 | 0.11 | 0.16 | 1.86 | 0.063 |

| 4-12 Hz coherence ~ 1 + Speed + Learning + Speed*Learning + (1 \| Animal) | | | | | | | | | |
|---|---|---|---|---|---|---|---|---|---|
| | **PFC - ACC** | | | **PFC - PPC** | | | **ACC - PPC** | | |
| | Estimate | t | p | Estimate | t | p | Estimate | t | p |
| Name | 0.43 | 30.08 | 4.63e-96 | 0.19 | 22.83 | 1.38e-69 | 0.26 | -32.54 | 1.99e-104 |
| Intercept | 0.79 | 1.69 | 0.092 | 0.38 | 1.40 | 0.16 | 0.33 | 1.29 | 0.20 |
| Learning | -0.081 | -3.60 | 3.66e-4 | -0.0094 | -0.72 | 0.47 | -0.076 | -6.12 | 2.72e-9 |
| Speed×Learning | -0.73 | -1.10 | 0.27 | -0.40 | -1.03 | 0.30 | -0.20 | -0.55 | 0.58 |
| | **HPC - PFC** | | | **HPC - ACC** | | | **HPC - PPC** | | |
| | Estimate | t | p | Estimate | t | p | Estimate | t | p |
| Name | 0.20 | 22.00 | 2.20e-66 | 0.21 | 26.86 | 1.29e-84 | 0.42 | 46.76 | 1.32e-146 |
| Intercept | 0.46 | 5.43 | 5.63e-8 | 0.27 | 1.04 | 0.30 | 0.32 | 0.91 | 0.36 |
| Learning | 0.079 | -1.95 | 7.06e-8 | -0.022 | -1.77 | 0.079 | -0.017 | -0.98 | 0.33 |
| Speed×Learning | -0.43 | 1.02 | 0.31 | -0.17 | -0.46 | 0.64 | -0.094 | -0.18 | 0.85 |

| 60-90 Hz coherence ~ 1 + Speed + Learning + Speed*Learning + (1 \| Animal) | | | | | | | | | |
|---|---|---|---|---|---|---|---|---|---|
| | **PFC - ACC** | | | **PFC - PPC** | | | **ACC - PPC** | | |
| | Estimate | t | p | Estimate | t | p | Estimate | t | p |
| Name | 0.27 | 47.40 | 2.77e-148 | 0.12 | 40.18 | 2.54e-128 | 0.12 | 40.27 | 1.39e-128 |
| Intercept | 0.11 | 0.58 | 0.57 | 0.026 | 0.27 | 0.79 | 0.0066 | 0.066 | 0.95 |
| Learning | 0.0086 | 0.95 | 0.34 | -0.018 | -3.78 | 1.88e-4 | -0.0093 | -1.93 | 0.055 |
| Speed×Learning | -0.069 | -0.26 | 0.80 | -0.056 | -0.40 | 0.69 | -0.046 | -0.32 | 0.75 |
| | **HPC - PFC** | | | **HPC - ACC** | | | **HPC - PPC** | | |
| | Estimate | t | p | Estimate | t | p | Estimate | t | p |
| Name | 0.11 | 41.65 | 1.45e-132 | 0.11 | 37.72 | 6.09e-121 | 0.12 | 45.81 | 4.44e-144 |

| | | | | | | | | | |
|---|---|---|---|---|---|---|---|---|---|
| Intercept | 0.032 | 0.38 | 0.71 | 0.04 | 0.44 | 0.66 | -0.0093 | -0.11 | 0.91 |
| Learning | 0.0057 | 1.39 | 0.17 | -0.0049 | -1.04 | 0.30 | -0.0093 | -2.25 | 0.025 |
| Speed×Learning | -0.058 | -0.48 | 0.63 | -0.131 | -0.94 | 0.35 | -0.010 | -0.082 | 0.93 |

**Table S2. [Overview of animals used across experiments and analyses], related to STAR Methods.**

| Animal ID | Gender | Group | Number of units/channels | | | | EEG and EMG (Y/N) | Oximeter (Y/N) | Contribution (figures) |
|---|---|---|---|---|---|---|---|---|---|
| | | | PFC | ACC | HPC | PPC | | | |
| #250624-1 | m | CS-US | 0/20 | 0/12 | 0/16 | 0/13 | Y | N | Fig.3A-3D; Fig. S1; Fig. S2; Fig. S5; Fig. S7; Fig. S8; Fig. S9; Table S1; |
| #250624-2 | m | CS-US | 0/20 | 0/12 | 0/16 | 0/13 | Y | N | |
| #250819-1 | m | CS-US | 0/20 | 0/12 | 0/16 | 0/13 | Y | Y | |
| #250819-2 | m | CS-US | 0/20 | 0/12 | 0/16 | 0/13 | Y | Y | |
| #251010-1 | m | CS-US | 0/20 | 0/12 | 0/16 | 0/13 | Y | Y | |
| #251010-2 | m | CS-US | 0/20 | 0/12 | 0/16 | 0/13 | Y | Y | |
| #240706 | m | CS only | 0/20 | 0/12 | 0/16 | 0/16 | N | N | Fig.1; Fig.2; Fig.3D-3F; Fig.5; Fig. S1; Fig. S5; Fig. S8L; Fig. S8M; Fig. S13; |
| #240716-1 | m | CS only | 0/20 | 0/12 | 0/16 | 0/16 | N | N | |
| #240726 | m | CS only | 0/20 | 0/12 | 0/16 | 0/16 | N | N | |
| #241111-1 | m | CS only | 0/20 | 0/12 | 0/16 | 0/16 | N | N | |
| #241128-1 | m | CS only | 0/20 | 0/12 | 0/16 | 0/16 | N | N | |
| #240702-1 | m | CS only | 0/20 | 0/12 | 0/16 | 0/16 | N | N | |
| #240705-1 | m | CS only | 0/20 | 0/12 | 0/16 | 0/16 | N | N | |
| #240714-1 | m | CS only | 0/20 | 0/12 | 0/16 | 0/16 | N | N | |
| #231115 | m | CS-US | 9/20 | 0/12 | 5/16 | 0/16 | N | N | |
| #231209 | m | CS-US | 4/20 | 0/12 | 2/16 | 0/16 | N | N | |
| #231215 | m | CS-US | 2/20 | 0/12 | 5/16 | 0/16 | N | N | |
| #240114-1 | m | CS-US | 1/20 | 0/12 | 8/16 | 0/16 | N | N | |
| #240519 | m | CS-US | 9/20 | 0/12 | 2/16 | 0/16 | N | N | |
| #240716-2 | m | CS-US | 26/20 | 0/12 | 6/16 | 0/16 | N | N | |
| #241111-2 | m | CS-US | 9/20 | 0/12 | 11/16 | 0/16 | N | N | |
| #241128-2 | m | CS-US | 12/20 | 0/12 | 4/16 | 0/16 | N | N | |
| #220819 | m | CS-US | 4/20 | 0/12 | 3/16 | 0/16 | N | N | Fig.2; FIg.3G-3N; Fig.5; Fig. 6I; Fig. S3; Fig. S4; Fig. S13; |
| #220829 | m | CS-US | 0/20 | 4/12 | 1/16 | 5/16 | N | N | |
| #220830 | m | CS-US | 8/20 | 2/12 | 5/16 | 4/16 | N | N | |
| #220905 | m | CS-US | 5/20 | 5/12 | 4/16 | 1/16 | N | N | |
| #220927 | m | CS-US | 6/20 | 1/12 | 3/16 | 0/16 | N | N | |
| #230404 | m | CS-US | 1/20 | 0/12 | 0/16 | 2/16 | N | N | |
| #230421 | m | CS-US | 2/20 | 1/12 | 3/16 | 0/16 | N | N | |
| #230427 | m | CS-US | 4/20 | 4/12 | 4/16 | 4/16 | N | N | |
| #230501 | m | CS-US | 4/20 | 2/12 | 8/16 | 3/16 | N | N | |
| #230508 | m | CS-US | 0/20 | 0/12 | 0/16 | 0/16 | N | N | |
| #230519 | m | CS-US | 1/20 | 1/12 | 2/16 | 0/16 | N | N | |
| #230520 | m | CS-US | 3/20 | 4/12 | 2/16 | 3/16 | N | N | |
| #230526 | m | CS-US | 0/20 | 5/12 | 0/16 | 1/16 | N | N | |
| #231026 | m | CS-US | 2/20 | 3/12 | 6/16 | 1/16 | N | N | |
| #240106 | m | CS-US | 14/20 | 8/12 | 6/16 | 5/16 | N | N | |

| | | | | | | | | | |
|---|---|---|---|---|---|---|---|---|---|
| #240114-2 | m | CS-US | 6/20 | 3/12 | 2/16 | 5/16 | N | N | |
| #CS_Only_1 | m | CS only | 0/20 | 0/12 | 0/16 | 0/16 | N | N | Fig.4; Fig. S10; Fig. S11; |
| #CS_Only_2 | m | CS only | 0/20 | 0/12 | 0/16 | 0/16 | N | N | |
| #CS_Only_3 | m | CS only | 0/20 | 0/12 | 0/16 | 0/16 | N | N | |
| #CS_Only_4 | m | CS only | 0/20 | 0/12 | 0/16 | 0/16 | N | N | |
| #CS_Only_5 | m | CS only | 0/20 | 0/12 | 0/16 | 0/16 | N | N | |
| #CS_Only_6 | m | CS only | 0/20 | 0/12 | 0/16 | 0/16 | N | N | |
| #CS_US_1 | m | CS-US | 0/20 | 0/12 | 0/16 | 0/16 | N | N | |
| #CS_US_2 | m | CS-US | 0/20 | 0/12 | 0/16 | 0/16 | N | N | |
| #CS_US_3 | m | CS-US | 0/20 | 0/12 | 0/16 | 0/16 | N | N | |
| #CS_US_4 | m | CS-US | 0/20 | 0/12 | 0/16 | 0/16 | N | N | |
| #CS_US_5 | m | CS-US | 0/20 | 0/12 | 0/16 | 0/16 | N | N | |
| #CS_US_6 | m | CS-US | 0/20 | 0/12 | 0/16 | 0/16 | N | N | |
| #240510 | m | CS-US+ LF interv. | 0/20 | 0/12 | 0/16 | 0/16 | N | N | Fig. 6; Fig. S14; |
| #240702-2 | m | CS-US+ LF interv. | 0/20 | 0/12 | 0/16 | 0/16 | N | N | |
| #240705-2 | m | CS-US+ LF interv. | 0/20 | 0/12 | 0/16 | 0/16 | N | N | |
| #240714-2 | m | CS-US+ LF interv. | 0/20 | 0/12 | 0/16 | 0/16 | N | N | |
| #240715 | m | CS-US+ LF interv. | 0/20 | 0/12 | 0/16 | 0/16 | N | N | |
| #CS_US_GmCL_1 | m | CS-US_GmCL | 0/20 | 0/12 | 0/16 | 0/16 | N | N | FIg.7A-7I; Fig. S16; Fig. S17; |
| #CS_US_GmCL_2 | m | CS-US_GmCL | 0/20 | 0/12 | 0/16 | 0/16 | N | N | |
| #CS_US_GmCL_3 | m | CS-US_GmCL | 0/20 | 0/12 | 0/16 | 0/16 | N | N | |
| #CS_US_GmCL_4 | m | CS-US_GmCL | 0/20 | 0/12 | 0/16 | 0/16 | N | N | |
| #CS_US_GmCL_5 | m | CS-US_GmCL | 0/20 | 0/12 | 0/16 | 0/16 | N | N | |
| #CS_US_RpCL_1 | m | CS-US_RpCL | 0/20 | 0/12 | 0/16 | 0/16 | N | N | Fig. S15; |
| #CS_US_RpCL_2 | m | CS-US_RpCL | 0/20 | 0/12 | 0/16 | 0/16 | N | N | |
| #CS_US_RpCL_3 | m | CS-US_RpCL | 0/20 | 0/12 | 0/16 | 0/16 | N | N | |
| #CS_US_RpCL_4 | m | CS-US_RpCL | 0/20 | 0/12 | 0/16 | 0/16 | N | N | |
| #CS_US_RpCL_5 | m | CS-US_RpCL | 0/20 | 0/12 | 0/16 | 0/16 | N | N | |
| #CS_US_RpCL_6 | m | CS-US_RpCL | 0/20 | 0/12 | 0/16 | 0/16 | N | N | |
| #CS_US_ThCL_1 | m | CS-US_ThCL | 0/20 | 0/12 | 0/16 | 0/16 | N | N | FIg.7J-7P; Fig. S16; Fig. S17; |
| #CS_US_ThCL_2 | m | CS-US_ThCL | 0/20 | 0/12 | 0/16 | 0/16 | N | N | |
| #CS_US_ThCL_3 | m | CS-US_ThCL | 0/20 | 0/12 | 0/16 | 0/16 | N | N | |
| #CS_US_ThCL_4 | m | CS-US_ThCL | 0/20 | 0/12 | 0/16 | 0/16 | N | N | |
| #Shock only_1 | m | Shock only | 0/20 | 0/12 | 0/16 | 0/16 | N | N | Fig. S5D; |
| #Shock only_2 | m | Shock only | 0/20 | 0/12 | 0/16 | 0/16 | N | N | |
| #Shock only_3 | m | Shock only | 0/20 | 0/12 | 0/16 | 0/16 | N | N | |
| #Shock only_4 | m | Shock only | 0/20 | 0/12 | 0/16 | 0/16 | N | N | |
| #Tetrodes-E1 | m | CS-US | 76/32 | 0/0 | 68/32 | 0/0 | N | N | Fig. S6; |
| #Tetrodes-E2 | m | CS-US | 42/32 | 0/0 | 77/32 | 0/0 | N | N | |
| #CNQX_1 | m | CNQX_D3-7 | 0/20 | 0/12 | 0/16 | 0/16 | N | N | Fig. S12; |
| #CNQX_2 | m | CNQX_D3-7 | 0/20 | 0/12 | 0/16 | 0/16 | N | N | |
| #CNQX_3 | m | CNQX_D3-7 | 0/20 | 0/12 | 0/16 | 0/16 | N | N | |
| #CNQX_4 | m | CNQX_D3-7 | 0/20 | 0/12 | 0/16 | 0/16 | N | N | |

| | | | | | | | | |
|---|---|---|---|---|---|---|---|---|
| #CNQX_5 | m | CNQX_D3-7 | 0/20 | 0/12 | 0/16 | 0/16 | N | N |
| #CNQX_6 | m | CNQX_D21-28 | 0/20 | 0/12 | 0/16 | 0/16 | N | N |
| #CNQX_7 | m | CNQX_D21-28 | 0/20 | 0/12 | 0/16 | 0/16 | N | N |
| #CNQX_8 | m | CNQX_D21-28 | 0/20 | 0/12 | 0/16 | 0/16 | N | N |
| #CNQX_9 | m | CNQX_D21-28 | 0/20 | 0/12 | 0/16 | 0/16 | N | N |
| #CNQX_10 | m | CNQX_D21-28 | 0/20 | 0/12 | 0/16 | 0/16 | N | N |
